# Differential roles of neural crest- and endothelial-derived FOXC2 in trabecular meshwork and Schlemm’s Canal in glaucomatous pathology

**DOI:** 10.1101/2022.02.20.481184

**Authors:** Pieter R. Norden, Lisa Beckmann, Raymond Fang, Naoto Ujiie, Zhen Cai, Xian Zhang, Junghun Kweon, Ting Liu, Kazushi Aoto, Susan E. Quaggin, Hao F. Zhang, Tsutomu Kume

**Affiliations:** Feinberg Cardiovascular and Renal Research Institute, Department of Medicine, Feinberg School of Medicine, Northwestern University, Chicago, Illinois, USA; Department of Biomedical Engineering, Northwestern University, Evanston, Illinois, USA; Department of Biochemistry, Hamamatsu University School of Medicine, Hamamatsu, Japan; Division of Nephrology and Hypertension, Northwestern University Feinberg School of Medicine, Northwestern University, Chicago, Illinois, USA; Department of Ophthalmology, Northwestern University, Chicago, Illinois, United States

## Abstract

Impaired development and maintenance of the Schlemm’s Canal (SC) is associated with perturbed aqueous humor outflow regulation and glaucoma progression. Key molecular mechanisms, such as ANGPT/TIE2, PROX1, and VEGF-C/VEGFR-3 regulate SC development and maintenance, but mechanisms of paracrine signaling from neighboring tissues, including the trabecular meshwork (TM) are poorly understood. Here, we show *Foxc2* is critical within the neural crest (NC)-derived TM and SC endothelium for development of the aqueous humor outflow pathway. In mice, NC-specific deletion of *Foxc2* results in abnormal anterior eye segment development, including impaired SC morphogenesis and functional maintenance, loss of SC identity, and impaired maintenance of intraocular pressure (IOP). Visible light optical coherence tomography angiography analysis also demonstrated functional impairment of the SC in response to changes in IOP in NC-*Foxc2^-/-^* mice, suggesting increased TM stiffness. Utilization of single-cell RNA-sequencing (scRNA-seq) analysis then identified that this phenotype is predominately characterized by transcriptional changes associated with extracellular matrix organization and stiffness in TM-associated cell clusters, including increased matrix metalloproteinase (MMP) expression, which can generate soluble TIE2 that acts as an ANGPT trap. As FOXC2 is also critically involved in development of the lymphatic vasculature in other tissues, we also show that endothelial-specific deletion of *Foxc2* resulted in impaired SC morphogenesis due to loss of TIE2 expression, which was rescued by deletion of the TIE2 phosphatase VE-PTP. Thus, NC-*Foxc2* is critical for development of the TM, and both NC- and endothelial-*Foxc2* are key for maintenance of SC identity and its morphogenesis.

## Introduction

Glaucoma is the second leading cause of visual impairment, affecting 3.6 million adults aged 50 or older among the 33.6 million recorded cases of visual impairment in 2020 (1, 2), and it is estimated that approximately 76 million people suffer from glaucoma globally as of 2020 (3). Primary congenital glaucoma (PCG) is characterized as developmental glaucoma occurring before three years of age due to obstructed drainage of aqueous humor via the conventional outflow pathway without overt structural defects of the eye. In contrast, developmental glaucoma occurs secondarily to observed malformations of the anterior segment of the eye (iridocorneal angle, ciliary muscle, etc.) (4). Aqueous humor functions to provide nourishment to the tissues of the anterior segment and to maintain pressure and the proper shape of the eye. It is secreted by the ciliary body, circulated into the anterior chamber, and returned to the circulation in part by the conventional outflow pathway consisting of flow through the trabecular meshwork (TM) and Schlemm’s Canal (SC) (5). Intraocular pressure (IOP) naturally results from outflow resistance generation within the conventional outflow pathway and elevated IOP is recognized as a critical risk factor contributing to optic neuropathy and the pathophysiology of glaucoma, including PCG and developmental glaucomas (4, 6, 7). Since elevated IOP is the primary and only modifiable risk factor for glaucoma, current treatments focus on lowering IOP by topical drugs, lasers, or surgical intervention.

Axenfeld-Rieger (AR) malformations refer to autosomal dominant developmental abnormalities of the anterior eye segment associated with mutations in the transcription factors paired-like homeodomain transcription factor 2 (*PITX2*) and forkhead box (*FOX*)*C1* and often results in the progression of glaucomatous blindness (8). In contrast to mutations in *FOXC1*, *FOXC2* mutations are predominately associated with lymphatic vascular dysfunction and the progression of the autosomal dominant lymphedema-distichiasis syndrome (9–11). However, during development both *Foxc1* and *Foxc2* share overlapping expression patterns and function cooperatively and complementary to one another in various aspects of tissue development, including blood and lymphatic vascular growth and maintenance (12–14). Interestingly, our group reported that neural crest (NC)-specific deletion of *Foxc2* reproduces similar, yet more mild, phenotypes associated with abnormal anterior eye segment development and corneal neovascularization observed in NC-*Foxc1* knockout mice. Yet, NC-*Foxc2* knockout mice (NC-*Foxc2^-/-^*) are viable, survive into adulthood, and are fertile (15). Supporting the pathophysiological importance of the role of *FOXC2* in ocular function and maintenance, it was recently reported that *FOXC2* variants were identified as modifier factors in a subset of congenital glaucoma patients (16).

It has previously been reported that SC malformations, or its complete absence, were observed in mice with global *Foxc1* and *Foxc2* haploinsufficiency (17). The SC is a large, hybrid vasculature with features characteristic of both lymphatic and venous vasculature (18–21). In contrast to the limbal and conjunctival lymphatics which originate from emergent lymphatic vessels on the nasal side of the developing eye, SC morphogenesis is initiated from the blood limbal and radial vascular plexuses during postnatal development (19, 22). Several key lymphatic vascular signaling pathways directly regulate SC morphogenesis and maintenance, such as vascular endothelial growth factor (VEGF)-C/ VEGF receptor (VEGFR)-3 (18), PROX1 (20), and angiopoietin (ANGPT)/TIE2 (21, 23–26). Of clinical importance, *TIE2* mutations have been previously identified in a subset of patients with PCG (23). Recently, TIE2 has emerged as a popular target for therapeutic intervention in PCG as TIE2 activation and the use of small-molecule inhibitors of negative TIE2 regulation have shown beneficial effects of improved SC morphology, increased outflow facility, and reduction of IOP in animal models of glaucoma (27, 28).

As NC-*Foxc2* transcriptional function has been identified to control ocular development and formation of the anterior eye segment in mice, we hypothesized that this model (15) would be valuable for investigating paracrine signaling mechanisms contributing to SC morphogenesis as the SC endothelium is not formed from NC-derived mesenchymal cells but adjacent to their derivatives in the TM (19). Here, we report that *Foxc2*+ cell descendants are present not only in NC-derived tissues within the anterior eye segment, but also within the SC endothelium. For the first time to our knowledge, we also demonstrate morphological and functional impairment of the SC in a transgenic model of anterior segment maldevelopment through utilization of a visible light optical coherence tomography (vis-OCT) system. Immunohistochemical analysis of SC morphology in NC-*Foxc2^-/-^* mice revealed that these individuals develop hypoplastic SC vasculature during morphogenesis with reduced expression of key lymphatic markers such as VEGFR-3, PROX1, and TIE2. Moreover, the SC is absent in individuals with severe anterior segment developmental defects. Single-cell RNA-sequencing (scRNA-seq) analysis of the anterior eye segment identified that transcriptional changes in TM cell cluster populations associated with NC-specific deletion of *Foxc2* were characterized by reduction of pro-angiogenic factors as well as increased expression of extracellular matrix (ECM) remodeling genes, including matrix metalloproteinases (MMPs). We then show that chemical inhibition of MMP activity in cultured human lymphatic endothelial cells (LECs) impaired cleavage of the TIE2 ectodomain and the production of soluble TIE2 (sTIE2), which is capable of binding angiopoietins and preventing them from activating TIE2 (29), suggesting a novel regulatory mechanism of TIE2 signaling activity in the SC. Similarly, early postnatal endothelial-specific deletion of *Foxc2* resulted in hypoplastic SC vasculature with reduced TIE2 expression.

However, conditional endothelial specific-deletion of one allele of *Ptprb*, encoding vascular endothelial protein tyrosine phosphatase (VE-PTP) was able to rescue this phenotype. Finally, using a conditional knock-in mouse line, we show that *Foxc2* can functionally substitute for *Foxc1* in the proper development of the anterior segment and SC vasculature. Collectively, our data show tissue-specific functions for *Foxc2* in the regulation of proper SC morphogenesis during development.

## Results

### *Foxc2*^+^ cell descendants are observed in the trabecular meshwork and Schlemm’s Canal Vasculature

Lineage tracing analysis performed using a tamoxifen-inducible *Foxc2^CreERT2^; R26R* reporter mouse line identified that descendants of mesenchymal *Foxc2*-expressing cells undergo division and proliferation to generate cells within the periocular and corneal mesenchyme during embryonic ocular development (30). We sought to first investigate how descendants of *Foxc2*-expressing cells contribute to the development of tissues comprising the conventional outflow pathway, including the TM and SC vasculature, beginning at early postnatal development when SC morphogenesis is initiated. To accomplish this, we crossed *Foxc2^CreERT2^* knock-in mice with the *ROSA^mT/mG^* reporter strain to generate a tamoxifen-inducible *Foxc2^CreERT2^; mTmG* strain and performed tamoxifen administration daily from postnatal day (P)1 to P5 during early SC morphogenesis, which occurs from P1 to approximately P15 – P17 where the SC vasculature has reached its mature morphology (19). Immunostaining analysis of CD31 expression in the limbal region of flatmounted tissue from *Foxc2^CreERT2^; mTmG* mice showed *Foxc2-Cre* mediated, eGFP-positive expression (indicative of *Foxc2^+^* cell descendants) in the CD31+ SC vasculature and adjacent trabecular meshwork (TM) at both P7 and P21 (**Figure 1, A and B**). Similarly, cryosection immunostaining analysis of the iridocorneal angle from *Foxc2^CreERT2^; mTmG* mice identified *Foxc2-Cre* mediated, eGFP-positive expression in the TM (**Figure 1C**) as well as the SC endothelium where select eGFP-positive cells also exhibited positive endomucin and VEGFR-3 expression (**Figure 1, D and E**). Together, these lineage-tracing observations demonstrate that *Foxc2^+^* cell descendants contribute to the maturation and development of the SC and TM during postnatal development (31, 32).

**Figure 1.**
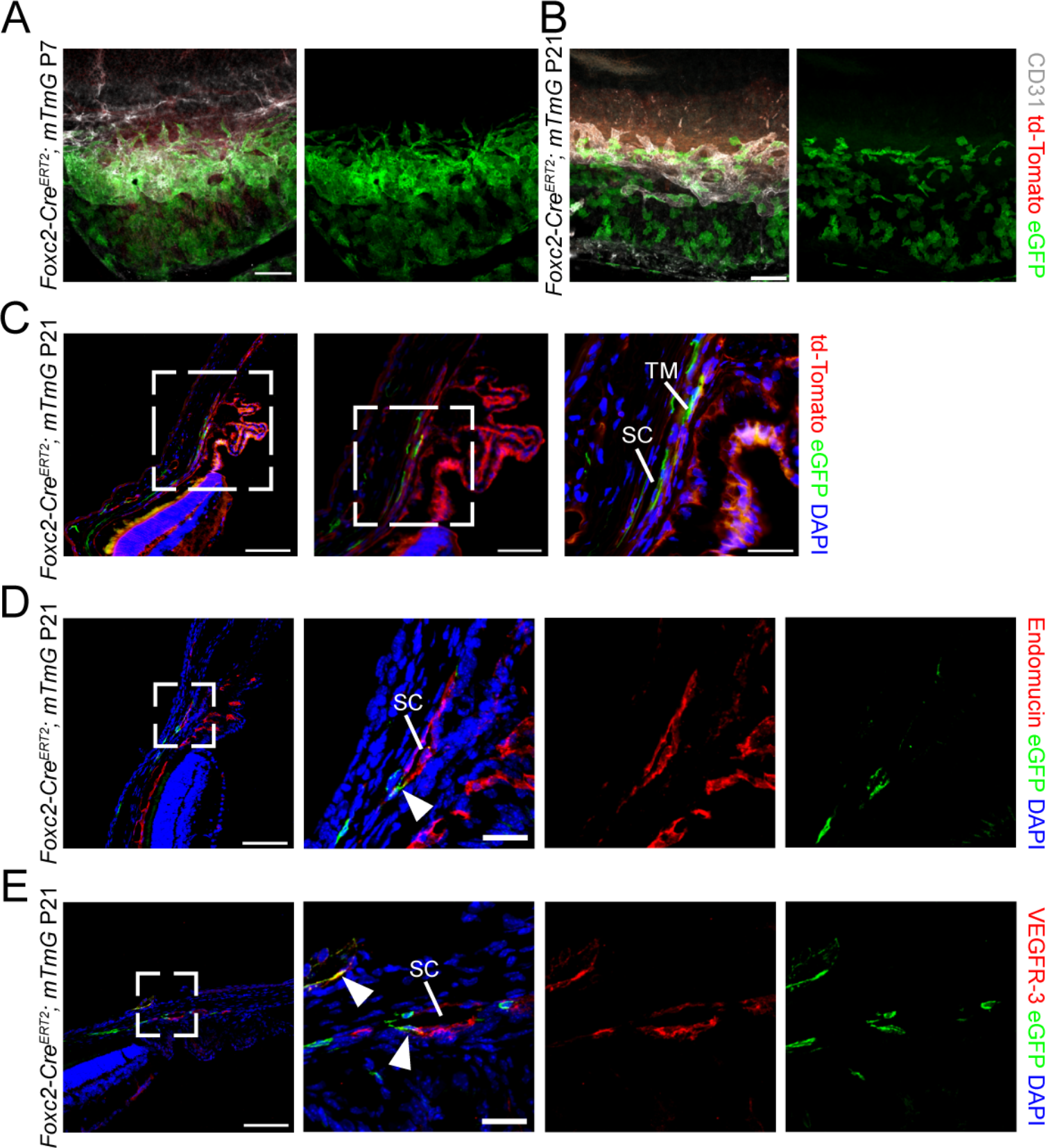
Fate-mapping of Foxc2 expression in cells of the trabecular meshwork and SC vasculature. (**A, B**) Representative image of the SC vasculature from P7 (**A**) and P21 (**B**) *Foxc2-Cre^ERT2^; mTmG* mice treated with Tamoxifen from postnatal day (P1) to P5 demonstrating Foxc2-GFP expression in the cells of the TM proximal to the SC as well as the SC vasculature denoted by CD31 immunostaining. Scale bar is 100 μm. (**C**) Representative cross-section images of eGFP-expressing, *Foxc2-Cre* mediated recombined cells in the SC and TM of a P21 *Foxc2-Cre^ERT2^; mTmG* mouse. Boxed areas denote magnified regions in sequential panels. TM –trabecular meshwork, SC – Schlemm’s Canal. Pink, dashed line denotes the SC vessel. Scale bars are 100, 50, and 25 μm respectively. (**D, E**) Representative cross-section images of eGFP- expressing, *Foxc2-Cre* mediated recombined cells in the SC of a P21 *Foxc2-Cre^ERT2^; mTmG* mouse immunostained with Endomucin (**D**) or VEGFR-3 (**E**). Boxed areas denote magnified regions in the next panels. SC –Schlemm’s Canal. Arrowheads denote GFP-positive and Endomucin- or VEGFR-3- positive cells. Scale bars are 100 and 25 μm respectively.

### Neural crest-specific deletion of *Foxc2* impairs anterior segment development and indirectly impairs Schlemm’s Canal morphology

Our group previously reported that NC-*Foxc2^-/-^* mice (*Wnt1-Cre; Foxc2^fl/fl^*) were characterized by an abnormal thickness of the peripheral corneas and conjunctiva, hypoplasia of the TM, defective, closed iridocorneal angles, and underdeveloped ciliary processes (15). To assess the direct contribution of NC-derived cell populations more carefully to the abnormal development of the anterior segment and conventional outflow pathway in NC-*Foxc2^-/-^* mice, we performed lineage tracing analysis by generating NC- *Foxc2^-/-^; mTmG* mice (*Wnt1-Cre; Foxc2^fl/fl^; ROSA^mT/mG^*). Cryosection immunostaining analysis of 3-week old NC-*Foxc2^-/-^; mTmG* mice showed similar abnormal defects in anterior segment tissues as to what our group reported (**Figure 1-figure supplement 1**) with eGFP-positive expression observed in the corneal stroma, TM, ciliary processes, and scleral tissue of NC-*Foxc2^-/-^; mTmG* mice compared to Cre-negative *Foxc2^fl/fl^; mTmG* controls (**Figure 1-figure supplement 1A**). Notably, analysis of sections showed regions of the iridocorneal angle where the SC appeared to be nearly absent or characterized by abnormal morphology with reduced area and a nearly closed vessel in NC-*Foxc2^-/-^; mTmG* mice compared to *Foxc2^fl/fl^; mTmG* controls (**Figure 1-figure supplement 1B – D**). As characterized in previously published reports (19), eGFP-positive expression was not observed in the SC endothelium following recombination mediated by the *Wnt1-Cre* driver whereas eGFP expression was observed in hypoplastic, PDGFRβ-positive TM (20) (**Figure 1-figure supplement 1, C and D**). Thus, loss of *Foxc2* expression and transcriptional activity in NC-derived cells during embryonic ocular development indirectly impairs SC morphogenesis during early postnatal development and maturation.

### NC-*Foxc2^-/-^* mice exhibit corneal neovascularization, reduced SC size, and normal relationship between SC and IOP

Our group has previously used both corneal flat-mount immunostaining and OCT imaging to show that NC-*Foxc2^-/-^* mice exhibit corneal neovascularization (15), however the characteristics of SC morphology in these individuals are unknown. To carefully assess possible phenotypes in SC morphology in these individuals, we utilized a more recently designed and established vis-OCT system to acquire compound circumlimbal scans to visualize the SC in live mice (33). We used this system to acquire vis-OCT angiography (vis-OCTA) volumes of the limbal region of both 6-8-week old Foxc2*^fl/fl^* control and NC-*Foxc2^-/-^* mice with milder ocular phenotypes, as mice with severe ocular phenotypes could not be accurately measured with vis-OCTA. Representative *en face* projections of the volumes show modest corneal neovascularization in NC-*Foxc2^-/-^* mice as previously reported (15) (**Figure 2, A and B, E and F**). Representative B-scan images from each location marked in red on the *en face* vis-OCTA images in **Figure 2, A and B** are shown in **Figure 2, C and D,** respectively. Here, the SC is clearly visible in the cross-sectional B-scan images showing the reduced area in NC-*Foxc2^-/-^* mice with milder ocular phenotypes.

**Figure 2.**
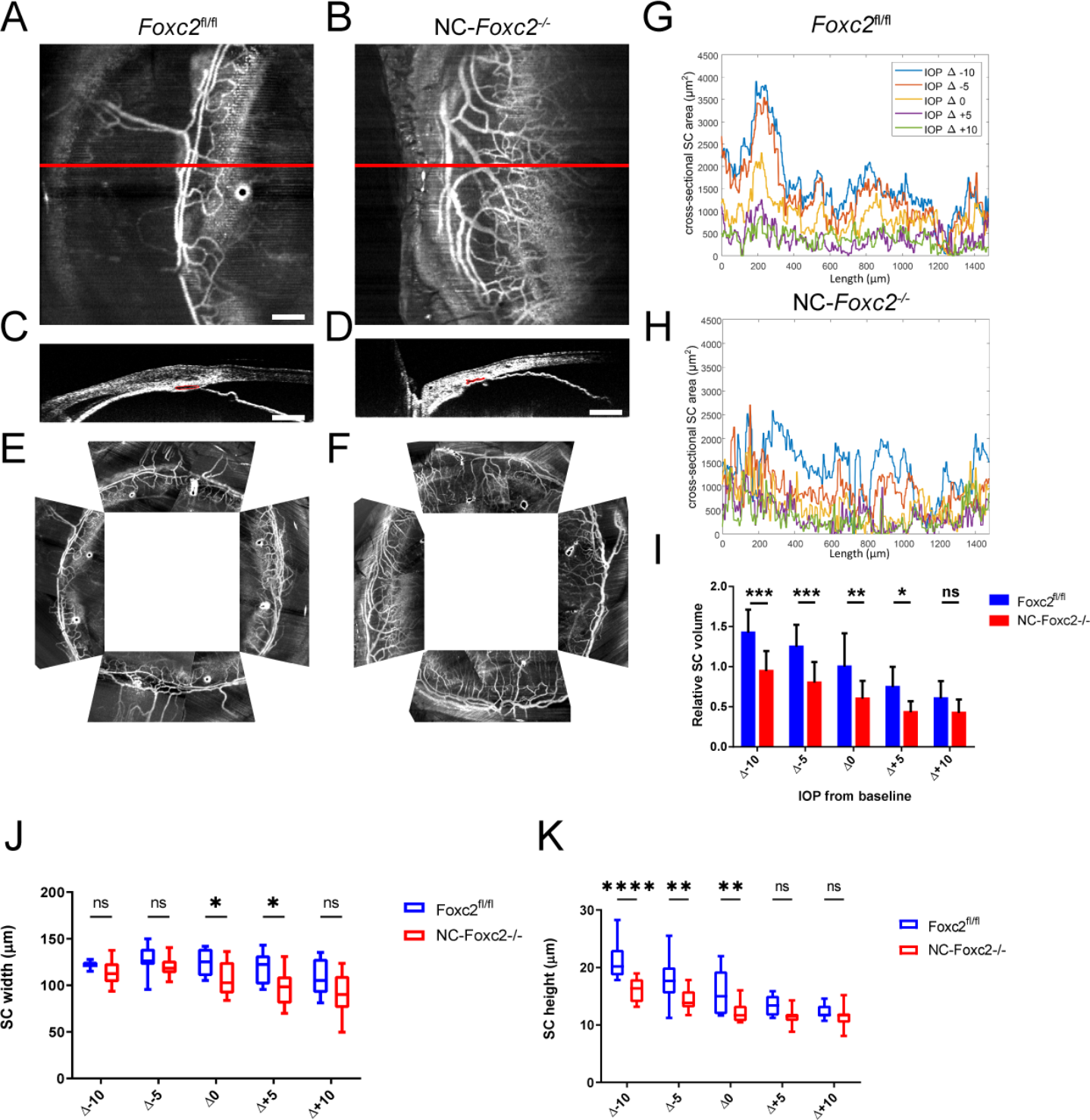
Vis-OCTA and vis-OCT identify corneal neovascularization and reduced SC area and volume in NC-*Foxc2*^-/-^ mice. (**A, B**) Representative angiograms of an individual vis-OCTA raster scan from adult, 7-month-old *Foxc2^fl/fl^* control (**A**) and NC- *Foxc2^-/-^* (**B**) mice. (**C, D**) Representative cross-sectional vis-OCT B-scan images from *Foxc2^fl/fl^* control (**C**) and NC-*Foxc2^-/-^* (**D**) mice. Red outline denotes SC area. (**E, F**) Representative compound circumlimbal scans composed of 8 separate raster angiograms each from *Foxc2^fl/fl^* control **(E)** and NC-*Foxc2^-/-^* (**F**) mice showing. Scale bar: 200 μm. (**G**) Representative plot of SC area versus length from one adult *Foxc2^fl/fl^* control mouse at IOP levels ranging from 10 mmHg below baseline IOP to 10 mmHg above baseline IOP. SC area is measured from segmented vis-OCT B-scan images. (**H**) Representative plots of SC area versus length from one adult NC-*Foxc2^-/-^* mice at IOP levels ranging from 10 mmHg below baseline IOP to 10 mmHg above baseline IOP. (**I**) Relative SC volume plotted against change in IOP, where the cycle of IOP changes was repeated 3 times in the same mouse to give a mean and standard error for N = 3 Control and N = 4 NC-*Foxc2^-/-^* mice. SC volume for each IOP and group was normalized by the mean *Foxc2^fl/fl^* control SC volume at baseline IOP. (**J, K**) Box-and-whisker plots of SC width and height plotted against change in IOP for N = 3 Control and N = 4 NC- *Foxc2^-/-^* mice where each IOP level was repeated 3 times. Statistical analysis: Two-way ANOVA with Šídák’s multiple comparisons test. * P < .05, ** P < .01, ***, P < .001, **** P < .0001.

To investigate the *in vivo* functional behavior of the SC in NC-*Foxc2^-/-^* mice, we assessed changes in SC morphology in response to changes from baseline IOP (33). Cross-sectional area versus length for 5 IOP levels relative to the measured baseline IOP, at - 10 mmHg, -5 mmHg, 0 mmHg, +5 mmHg, and +10 mmHg, was quantified for both Foxc2*^fl/fl^* control and NC-*Foxc2^-/-^* mice (**Figure 2, G and H**). In both, there is an inverse relationship between IOP and SC cross-sectional area. Additionally, in both, conserved regions of higher cross-sectional area across all IOP levels can be observed. Overall, the SC volume is smaller in NC-*Foxc2^-/-^* mice at IOP levels of Δ-10 mmHg, Δ-5 mmHg, Δ0 mmHg, and Δ+5 mmHg (**Figure 2I**). However, relative changes between IOP levels match the same pattern as in control mice, with an inverse relationship between IOP and SC volume (34). Quantification of SC width demonstrated that it is smaller in NC-*Foxc2^-/-^* when compared to Foxc2*^fl/fl^* control at Δ0 mmHg and Δ+5 mmHg (**Figure 2J**). Additionally, SC height is smaller in NC-*Foxc2^-/-^* when compared to Foxc2*^fl/fl^* control at Δ-10 mmHg, Δ- 5 mmHg, and Δ0 mmHg. (**Figure 2K**). Collectively, these data demonstrate that NC-specific loss of *Foxc2* results in abnormal SC morphology and impaired function in response to changes in IOP.

### The SC of NC-*Foxc2^-/-^* mice is characterized by abnormal morphology and failure to properly establish SC endothelial identity

We additionally characterized morphological and phenotypic changes in the SC associated with NC-specific loss of *Foxc2* in adult mice. Flatmount immunohistochemistry analysis of CD31 expression within the limbal region of 6 – 8-week-old *Foxc2^fl/fl^*, NC-*Foxc2^-/+^* (*Wnt1-Cre; Foxc2^fl/+^*), and NC-*Foxc2^-/-^* mice demonstrated that retainment of one allelic copy of *Foxc2* is sufficient to maintain proper SC morphogenesis, however loss of NC-*Foxc2* results in a hypoplastic SC vasculature, typically displaced from the limbal blood and lymphatic vascular plexus (**Figure 3A**).

**Figure 3.**
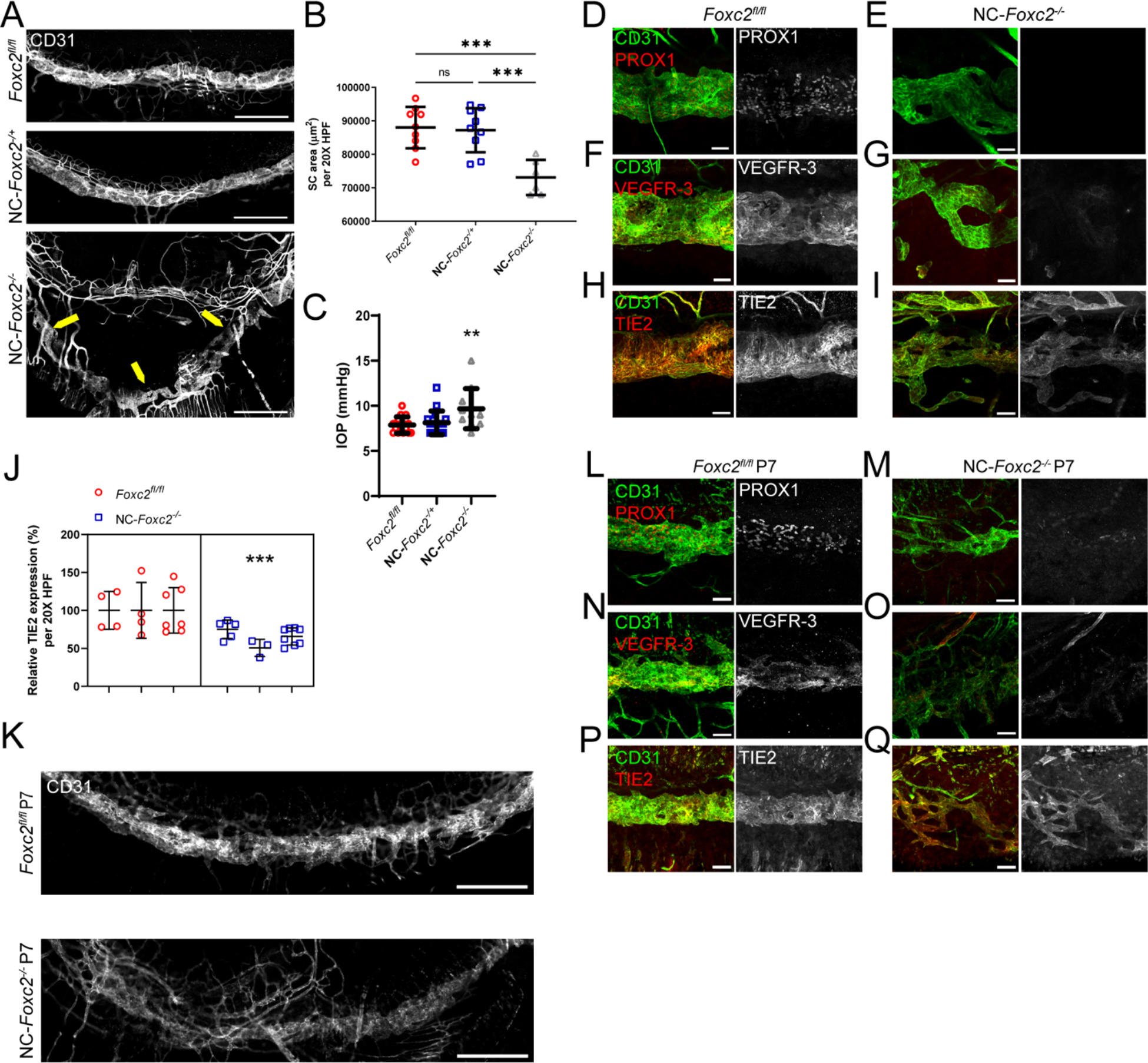
Neural crest derived *Foxc2* is required for proper morphogenesis of the SC and establishment of SC identity. (**A**) Representative images of SC vasculature immunostained with CD31 antibody in adult, *Foxc2^fl/fl^* control, NC-*Foxc2^-/+^*, and NC- *Foxc2^-/-^* mice. Yellow arrows highlight abnormal SC morphology and displacement in a NC-*Foxc2^-/-^* individual. Scale bars are 500 μm. (**B**) Quantification of SC area per 20X high-power field (HPF). N = 9 for *Foxc2^fl/fl^* controls. N = 9 for NC-*Foxc2^-/+^*. N = 6 for NC- *Foxc2^-/-^*. Data are mean ± SD. (**C**) Quantification of IOP in 7–8-week-old mice anesthetized with ketamine/doxylamine measured by a rebound tonometer. N = 10 for *Foxc2^fl/fl^* controls. N = 9 for NC-*Foxc2^-/+^*. N = 8 for NC-*Foxc2^-/-^*. Data are mean ± SD. Statistical analysis: One-way ANOVA with Tukey’s multiple comparisons test. ** P < .01,*** P < .001. (**D–I**) Representative images of CD31 and PROX1 (**D, E**), VEGFR-3 (**F, G**), or Tie2 (**H, I**) expression in the SC of adult *Foxc2^fl/fl^* Control (**D, F, H**) and NC-*Foxc2^- /-^* mice (**E, G, I**). Scale bars are 50 μm. (**J**) Quantification of relative expression of Tie2 in the SC of *Foxc2^fl/fl^* Control and NC-*Foxc2^-/-^* mice per 20X high-power field. N = 3 for *Foxc2^fl/fl^* controls and N = 3 for NC-*Foxc2^-/-^*. Symbols depict technical replicates per individual. Data are mean ± SD. Statistical analysis: Nested unpaired t-test. *** P < .001. (**K**) Representative images of SC vasculature immunostained with CD31 antibody show abnormal morphology and reduced area in P7 NC-*Foxc2^-/-^* mice compared to P7 *Foxc2^fl/fl^* Control mice. Scale bars are 500 μm. (**L–Q**) Representative images of CD31 and PROX1 (**L, M**), VEGFR-3 (**N, O**), or Tie2 (**P, Q**) expression in the SC of P7 *Foxc2^fl/fl^* Control (**L, N, P**) and NC-*Foxc2^-/-^* mice (**M, O, Q**). Scale bars are 50 μm.

Quantification of CD31 immunostained SC area was significantly reduced in NC-*Foxc2^-/-^* mice compared to *Foxc2^fl/fl^* controls whereas no differences were observed in NC- *Foxc2^-/+^* mice (**Figure 3B**). Of note, the penetrance of the severity of the SC phenotype was variable in NC-*Foxc2^-/-^* mice with a subset of individuals characterized by the absence of SC and severe corneal neovascularization (10/26, 38.5%). Rebound tonometry has been previously utilized as a non-invasive method for measuring IOP in mice under anesthesia (18) although evidence has shown that general anesthetics reduce IOP in normal mice compared to when they are alert (35). We measured IOP in anesthetized mice prior to their utilization in vis-OCT imaging analysis and observed increased IOP in NC-*Foxc2^-/-^* mice with milder ocular phenotypes and lacking severe corneal neovascularization (**Figure 3C**), indicative of increased outflow resistance.

As SC morphogenesis was impaired in NC-*Foxc2^-/-^* mice, we sought to characterize the expression of key molecular regulators of SC morphogenesis and maintenance (**Figure 3D – I**). PROX1 and VEGFR-3 expression was nearly absent in the hypoplastic SC vasculature of adult NC-*Foxc2^-/-^* mice (**Figure 3, E and G**), whereas TIE2 expression was modestly, but significantly, reduced (**Figure 3, I and J**) implying that the SC vasculature in NC-*Foxc2^-/-^* mice fails to properly establish or maintain SC endothelial identity. To address this discrepancy, we characterized SC vasculature in NC-*Foxc2^-/-^* mice and *Foxc2^fl/fl^* controls at P7 during the mid-stage of SC morphogenesis (**Figure 3, K – Q**). Like adult mice, the SC vasculature was absent in neonates with severe ocular defects, or hypoplastic in NC-*Foxc2^-/-^* mice with moderately abnormal ocular phenotypes compared to *Foxc2^fl/fl^* controls (**Figure 3K**), PROX1 and VEGFR-3 expression was nearly absent (**Figure 3, M and O**), and TIE2 expression was modestly reduced in neonatal NC-*Foxc2^-/-^* mice (**Figure 3Q**). Thus, early morphogenesis of the SC, as well as establishment and maintenance of SC endothelial identity, is impaired in NC-*Foxc2^-/-^* mice.

### Single-cell transcriptome analysis identifies molecular signaling pathway alterations in anterior-segment tissue populations of NC-*Foxc2^-/-^* mice

To understand the contribution of NC-*Foxc2* transcriptional regulation of development of the conventional outflow pathway at the molecular level, we performed single-cell RNA-seq (scRNA-seq) analysis of pooled anterior eye segments from 3-4-week old *Foxc2^fl/fl^* control mice and NC-*Foxc2^-/-^* mice with moderate ocular phenotypes. Young mice after weaning were utilized to optimize the total cell number utilized for single-cell sequencing, and NC-*Foxc2^-/-^* mice with moderate phenotypes were selected so that the anterior eye segment could be definitively dissected from the posterior segment. T-distributed stochastic neighbor embedding (t-SNE) visualization and clustering analysis of cells from both Foxc2*^fl/fl^* Control and NC-*Foxc2^-/-^* mice identified 22 transcriptionally distinct cell populations including uveal meshwork, in which aqueous humor first traverses through the TM, 3 transcriptionally distinct trabecular meshwork populations, and endothelial cells among other cell populations (**Figure 4, A – D, Figure 4-figure supplement 1**) characterized from previously reported scRNA-seq analysis studies of the murine conventional outflow pathway and anterior eye segment (36–40) . *Foxc2* expressing cells were detected in several cell clusters, but were primarily localized to trabecular meshwork, corneal stromal keratocyte, and pericyte populations (**Figure 4E**). Quantification of *Foxc2* expression in both *Foxc2^fl/fl^* and NC-*Foxc2^-/-^* cells validated reduction of *Foxc2* expression in our scRNA-seq dataset (**Figure 4F**).

**Figure 4.**
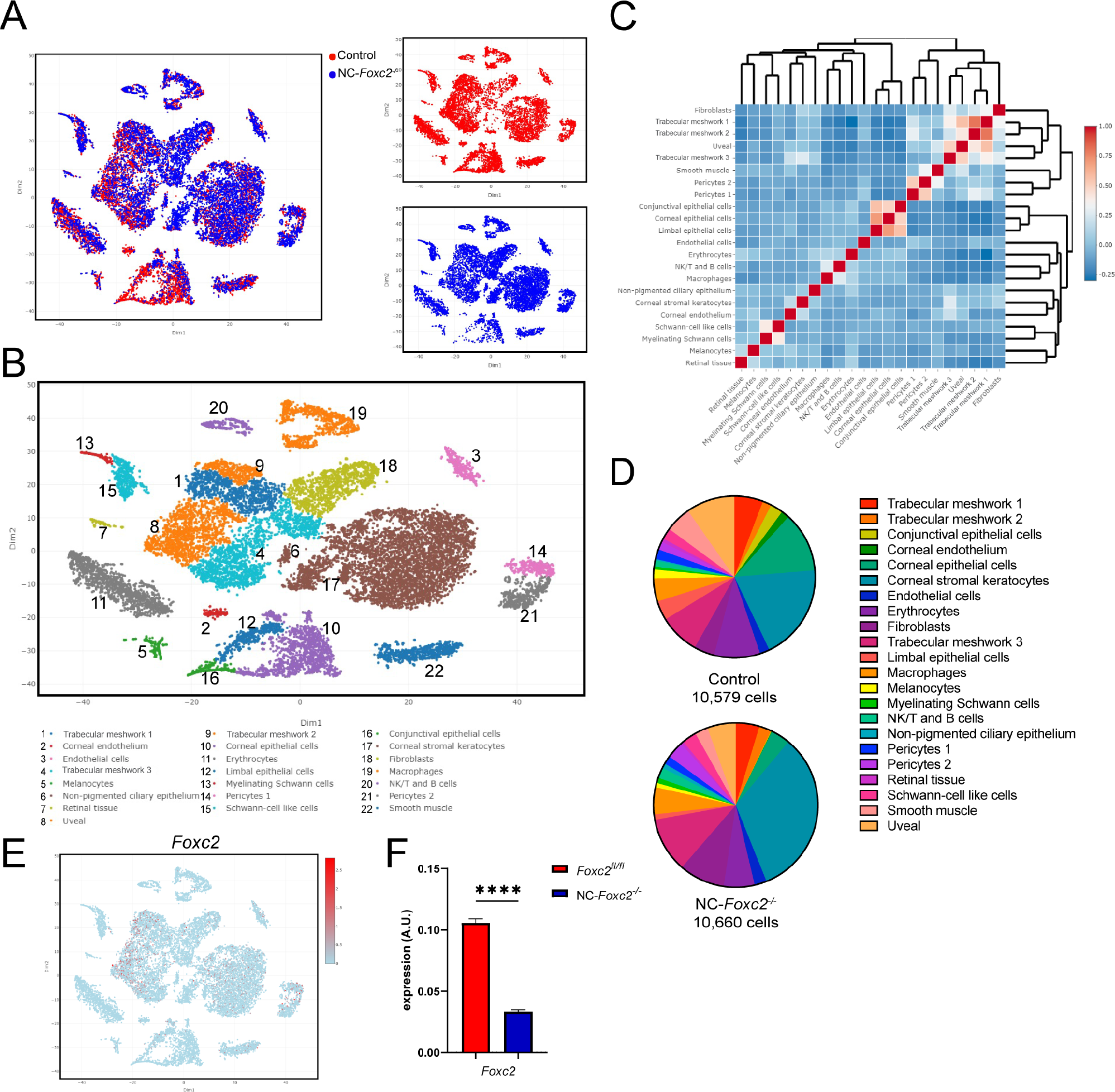
Single-cell transcriptome profiling of cell populations in the anterior eye segment of Control and NC-*Foxc2^-/-^* mice. (**A, B**) Visualization of individual cell contribution (**A**) and unsupervised clustering analysis of 22 transcriptionally distinct cell populations (**B**) by t-distributed stochastic neighbor embedding (t-SNE) in the anterior eye segment of 3–4-week-old *Foxc2^fl/fl^* Control and NC-*Foxc2^-/-^* mice. (**C**) Correlation heatmap and hierarchical clustering of gene-expression signatures of all cell populations. (**D**) Pie charts demonstrating the proportion of the total cell number analyzed contributing to each individual cell cluster for Control and NC-*Foxc2^-/-^* mice. (**E**) t-SNE plot colorized by the Seurat normalized expression *Foxc2*. (**F**) Quantitative analysis of *Foxc2* expression in the entirety of *Foxc2^fl/fl^* Control and NC-*Foxc2^-/-^* cells. Data are mean ± SEM. Statistical analysis: Student’s unpaired t-test. **** denotes p < .0001.

As NC-derived cells do not contribute to the development of the SC vasculature (19), we focused our analyses on assessment of uvea meshwork and TM cell populations provided their anatomical proximity to SC and positive *Foxc2* expression detected in these clusters. Previously reported single cell analyses of the outflow pathway have identified uniquely expressed markers within different cell populations comprising the TM, such as fibroblast-like trabecular “beam” cells comprising the inner part of the posterior filtering region of the TM, the juxtacanalicular tissue (JCT) comprising the outer part of the posterior filtering region located adjacent to the SC, and the possible presence of *Cd34*-positive corneal stromal cells/fibroblasts clustering within TM cell populations as recently characterized by Thomson et al. (37, 38, 40). Our initial analysis showed that known marker genes of TM, such as *Myoc* and *Chil1,* were predominately expressed in the Trabecular meshwork 3 (TM-3) cluster, with less prominent expression in the Trabecular meshwork 1 and 2 (TM-1 and TM-2) clusters (**Figure 4-figure supplement 1C, Figure 4-figure supplement 2A and D**). However, TM-1 and TM-2 both highly expressed *Dcn* and *Pdgfra* (**Figure 4-figure supplement 2A**), consistent with TM populations characterized in previously reported single cell analyses (37, 38).

TM-1 and TM-2 also contained high levels of *Edn3*, which may suggest that they are more closely related to the “Beam” cell populations previously characterized by van Zyl et al (**Figure 4-figure supplement 2B**). In contrast, TM-3 showed higher expression of *Chad*, *Chil1*, *Nell2*, and *Tnmd* (**Figure 4-figure supplement 2D**), which has previously been reported to be characteristic of the JCT. The uveal meshwork cluster also exhibited higher expression of *Col23a1* and *Lypd1* (**Figure 4-figure supplement 2D**), consistent with previous reports (38, 40). Our analysis also showed that our putative TM clusters expressed several fibroblast markers, such as *Cd34*, *Clec3b*, *Mfap5*, *Pi16*, and *Tnxb*, although their expression was particularly enriched in a separate cluster more characteristic of corneal stromal fibroblasts (**Figure 4-figure supplement 2B**).

To investigate transcriptional changes in signaling pathways potentially contributing to impaired SC morphogenesis, we performed analysis of differentially expressed genes (DEGs) in uveal and TM cell populations and identified significant transcriptional changes within these populations (**Figure 5, Supplementary Table 1**) and, in particular, TM-3 (**Figure 5D**). Analysis of gene ontology (GO) biological processes from DEGs within these populations elucidated several meaningful biological changes associated with NC-specific loss of *Foxc2*. Downregulated processes that were common among TM populations included blood vessel and vasculature development (**Figure 5, A, B and D**) and regulation of cell adhesion (**Figure 5, A and D**). Similarly, upregulated processes that were common among TM populations included blood vessel development (**Figure 5D)** and morphogenesis (**Figure 5B**) as well as regulation of cell adhesion **(Figure 5, A and B**). However, GO terms associated with ECM function were most upregulated among TM populations, including ECM organization, collagen chain trimerization, extracellular structure organization, and ECM-receptor interaction (**Figure 5, A, B, and D**). The JCT is located adjacent to the inner wall endothelium of the SC (41). Its cells can extend processes that communicate with both the inner wall endothelium and TM cells in the corneoscleral meshwork (42). Although GO analysis identified that blood vessel development was both associated with downregulated and upregulated genes in TM-3 (characteristic of a putative JCT population), pro-angiogenic growth factors and secreted proteins such as *Adm*, *Amot*, *Angptl4*, *Ecm1*, *F3*, *Hbegf*, *Jag1*, *Lgals3*, *Lox*, *Serpine1*, *Tgfb2*, and *Vegfa* were significantly downregulated (**Figure 5-figure supplement 1A, Supplementary Table 1**) whereas several anti-angiogenic signaling factors including *Adamts1, Sema3C,* and *Tgfbi* were significantly upregulated (**Figure 5-figure supplement 1B**), possibly contributing to abnormal endothelial cell signaling and impaired SC morphogenesis.

**Figure 5.**
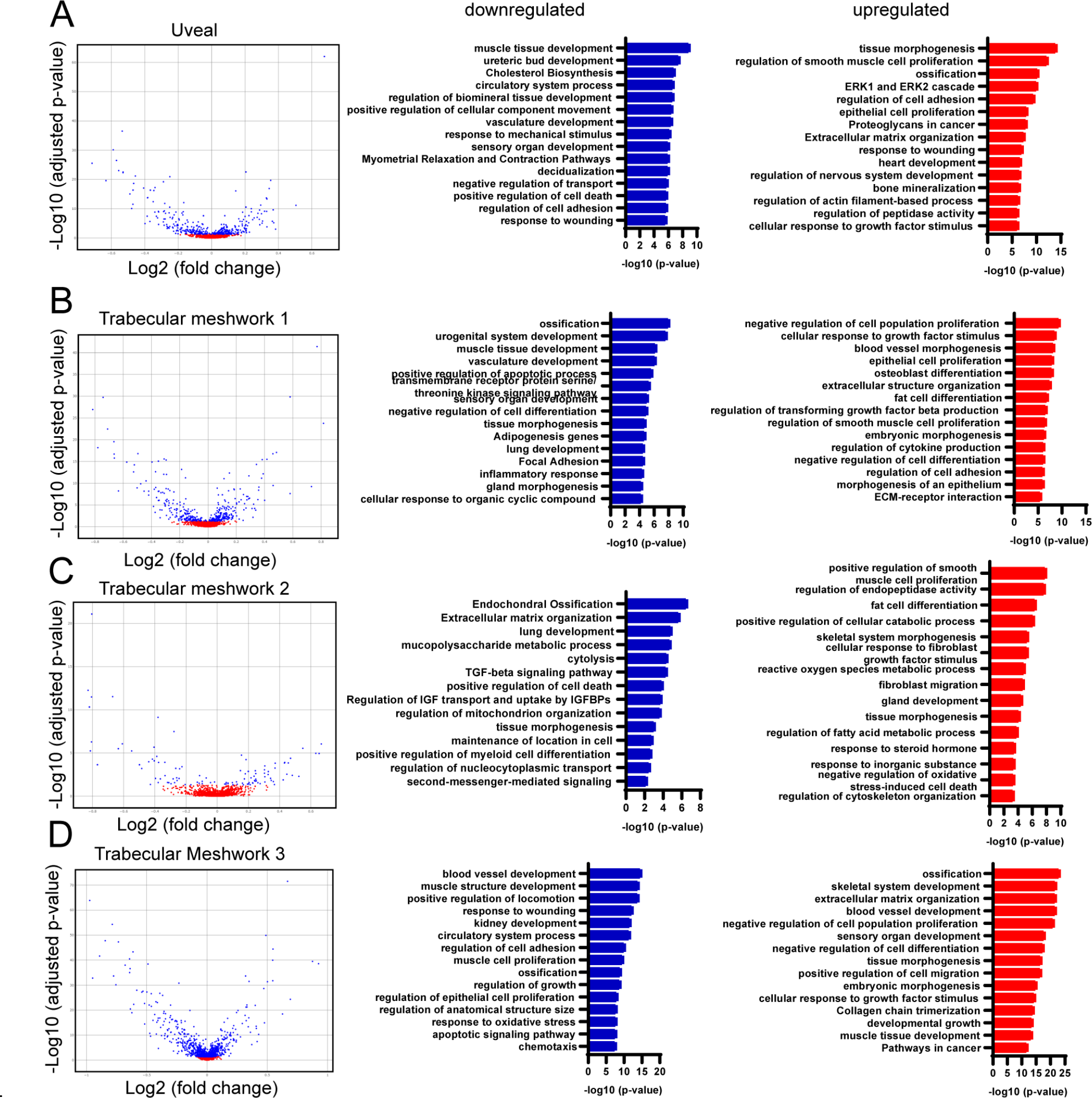
Differentially expressed genes in trabecular meshwork cell populations of NC-*Foxc2^-/-^* mice compared to *Foxc2^fl/fl^* Controls and GO over-representation analysis. (A-D) Volcano plots showing DEGs between NC-*Foxc2^-/-^* and *Foxc2^fl/fl^* Control mice in the Trabecular meshwork 1 (A), Trabecular meshwork 2 (B), Trabecular meshwork 3 (C), and uveal meshwork clusters (D), left panels. Blue points denote DEGs with adjusted p-value < .05. Bar plots of subsets of GO gene sets that were over- represented among the genes downregulated (blue bars) or upregulated (red bars) in NC-*Foxc2^-/-^* mice compared to *Foxc2^fl/fl^* Control mice, right panels. Values on the x-axis are represented as the –log10(p-value) of each associated GO gene set.

The TM primarily functions to regulate bulk aqueous humor outflow from the anterior chamber, which is accomplished by generating resistance to outflow (43). Previously reported work has demonstrated that forces generated by increased IOP likely act on the ECM and attached cells of the JCT and SC inner wall endothelium by stretching and distorting them, which in turn stimulates ECM processing and turnover initiated by MMP signaling in the TM (44, 45). Our scRNA-seq analysis revealed that ECM related genes are significantly upregulated in TM populations of NC-*Foxc2^-/-^* mice, including several collagen genes and MMPs such as *Adam19, Adamts1, Adamts5, Mmp2,* and *Mmp3* in TM-3 (**Figure 5-figure supplement 1B and C**), among other upregulated ECM related components (**Supplementary Table 1**). In partial validation of our scRNA-seq observations, immunostaining of MMP-2 demonstrated its expression was increased throughout the anterior segment of NC-*Foxc2^-/-^* mice compared to *Foxc2^fl/fl^* controls (**Figure 6**). Co-localization of increased MMP-2 expression with PDGFRβ+ TM cells and increased expression in the cornea was observed in NC-*Foxc2^-/-^* mice with particularly severe anterior segment developmental phenotypes, which lacked a mature SC as assessed by VEGFR-3 expression and anatomical features (**Figure 6B**), but it was weakly detected in PDGFRβ+ TM of *Foxc2^fl/fl^* control mice (**Figure 6A**). Although our scRNA-seq analysis determined that increased *Mmp2* expression was limited to TM-3, it is possible that dosage-dependent effects of *Mmp2* expression among other genes may be attributable to the differences we observe in penetrance and severity of the phenotype observed in NC-*Foxc2^-/-^* mice as individuals with more moderate ocular phenotypes were utilized for scRNA-seq analysis.

**Figure 6.**
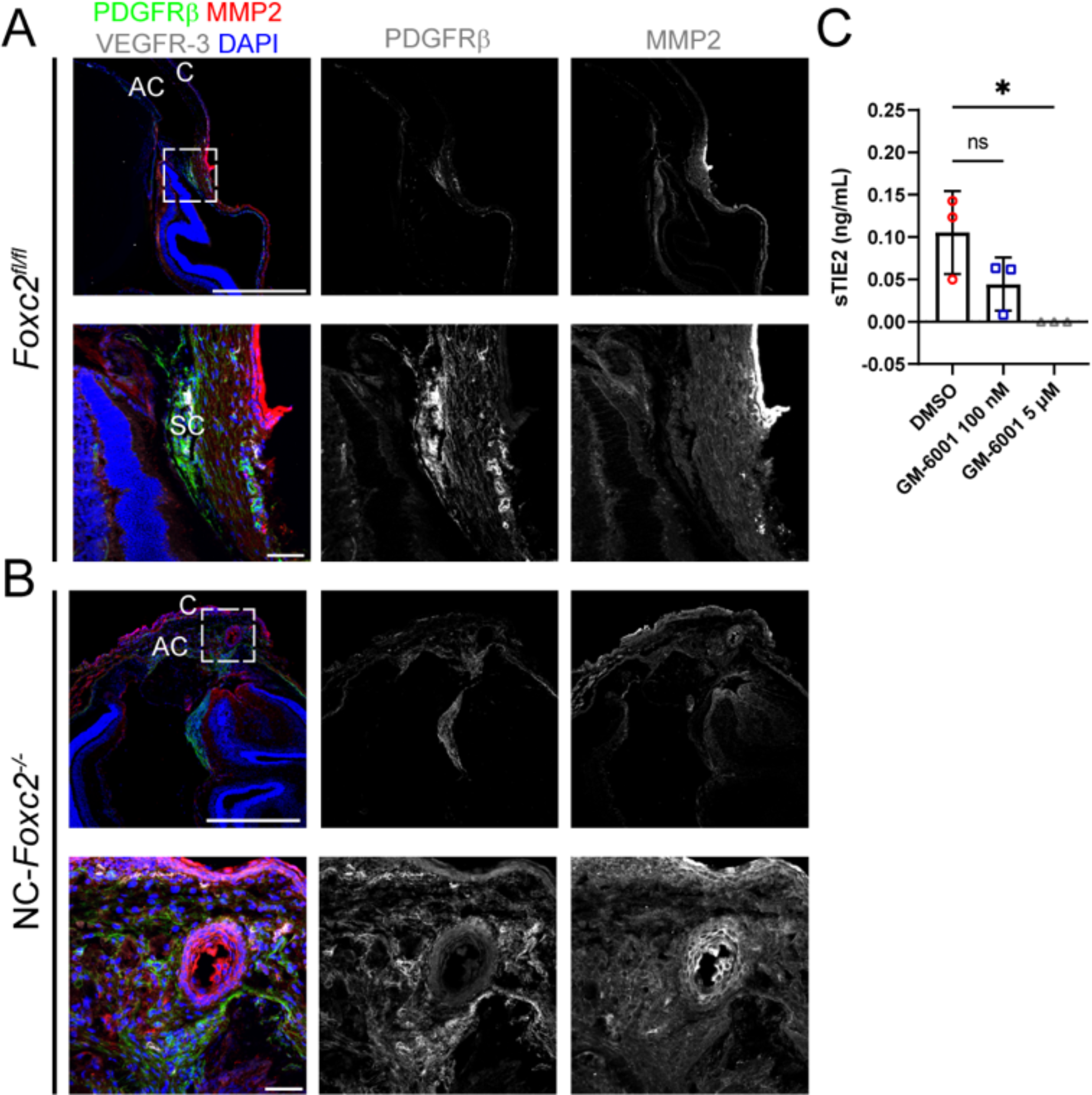
MMP-2 expression is abnormally increased in anterior segment tissues of NC-*Foxc2^-/-^* mice and MMP activity mediates TIE2 shedding in cultured LECs. (A, B) Representative cross-section images of the iridocorneal angle and anterior chamber of a *Foxc2^fl/fl^* control (A) and NC-*Foxc2^-/-^* individual immunostained with PDGFRβ, MMP2, VEGFR-3, and DAPI. Boxed regions in top panels denote magnified regions in lower panels. Scale bars are 500 μm and 50 μm respectively. C – cornea, AC – anterior chamber, SC – Schlemm’s Canal. (C) Quantification of sTIE2 in conditioned media from cultured HDLECs treated with DMSO vehicle or GM-6001 at 100 nM or 5 μm by ELISA. Data are mean ± SD from three biological replicates. Statistical analysis: Kruskal-Wallis test with Dunn’s multiple comparisons. * denotes p < .05.

Of particular note, previously reported work has demonstrated that MMP activity mediates cleavage of the TIE2 ectodomain to produce sTIE2 in cultured human umbilical vein endothelial cells (HUVECs) (29, 46). Moreover, sTIE2 is capable of binding angiopoietins to prevent them from activating TIE2 signaling (29). Provided the increase expression of MMPs in NC-*Foxc2^-/-^* mice and the crucial role of ANGPT/TIE2 signaling in SC development (25–27, 40), we hypothesized that increased MMP activity in the TM may mediate cleavage of TIE2 in SC endothelium, resulting in higher production of sTIE2, impairment of ANGPT/TIE2 signaling, and impaired SC morphogenesis. To investigate this mechanism, we collected conditioned media from cultured human dermal LECs (HDLECs) treated with either DMSO vehicle or the MMP inhibitor GM-6001 at dosages shown to inhibit sTIE2 shedding in cultured HUVECs (29) or impair tubulogenesis in three-dimensional culture assays (47). Quantification of sTIE2 concentration by ELISA demonstrated there was a trend in reduction of sTIE2 at a lower dosage of GM-6001 treatment, but that the higher dosage resulted in no detection of sTIE2 in conditioned media (**Figure 6C**). Thus, like blood Ecs, MMP activity also mediates TIE2 shedding in LECs, and the role of MMP signaling and potential cleavage of the TIE2 ectodomain warrants further investigation within the SC.

To directly investigate potential transcriptional changes in the SC endothelium secondarily associated with the loss of NC-*Foxc2* transcriptional activity, we performed subclustering analysis of the endothelial cell cluster (**Figure 4B**) identified in our dataset to identify the population of SC endothelium. Uniform manifold approximation and projection (UMAP) visualization and clustering analysis identified 8 transcriptionally unique clusters consisting of 6 blood endothelial cell (BEC) clusters, a cluster specific to NC-*Foxc2^-/-^* mice, and a cluster comprised of both SC and limbal lymphatic Ecs (**Figure 7, A and B**). The BEC-1 and BEC-4 clusters exhibited higher expression of venous endothelial markers such as *Ackr1*, *Mgp*, *Sele*, *Selp*, and *VWF* (**Figure 7-figure supplement 1A**). The BEC-1 cluster exhibited the strongest expression of these genes, potentially indicating that this cluster may be characteristic of collector channel vessels that exhibited a similarly high expression of these genes in a human dataset (38). The other BEC clusters showed stronger expression for markers recently reported to be associated with arterial limbal endothelium (40), with the exception of BEC-3, which exhibited higher expression of *Ihh*, a recently identified marker of choriocapillaris (40, 48) that may have been incorporated during dissection and tissue dissociation (**Figure 7-figure supplement 1B and C**). The SC and lymphatic EC cluster showed high expression of several markers identified in SC Ecs such as *Ccl21a*, *Flt4*, and *Prox1*. However, only a few cells exhibited positive expression of the classic lymphatic markers *Lyve1* and *Pdpn* (**Figure 7C**), which are absent in the SC, implying that this cluster predominately consists of SC Ecs with few limbal lymphatic Ecs that were not independently clustered during analysis. In support of this observation, the SC and lymphatic EC cluster exhibited high expression of *Npnt*, *Nts*, *Pgf*, and *Postn* and modest expression of *Itga9* and *Nts* (**Figure 7C and D)** and other endothelial marker genes including *Flt1*, *Kdr*, *Plvap*, *Ptprb*, and *Tek* (**Figure 7-figure supplement 1D**), which were recently shown to be characteristic of the SC endothelial transcriptional profile by Thomson et al (40).

**Figure 7.**
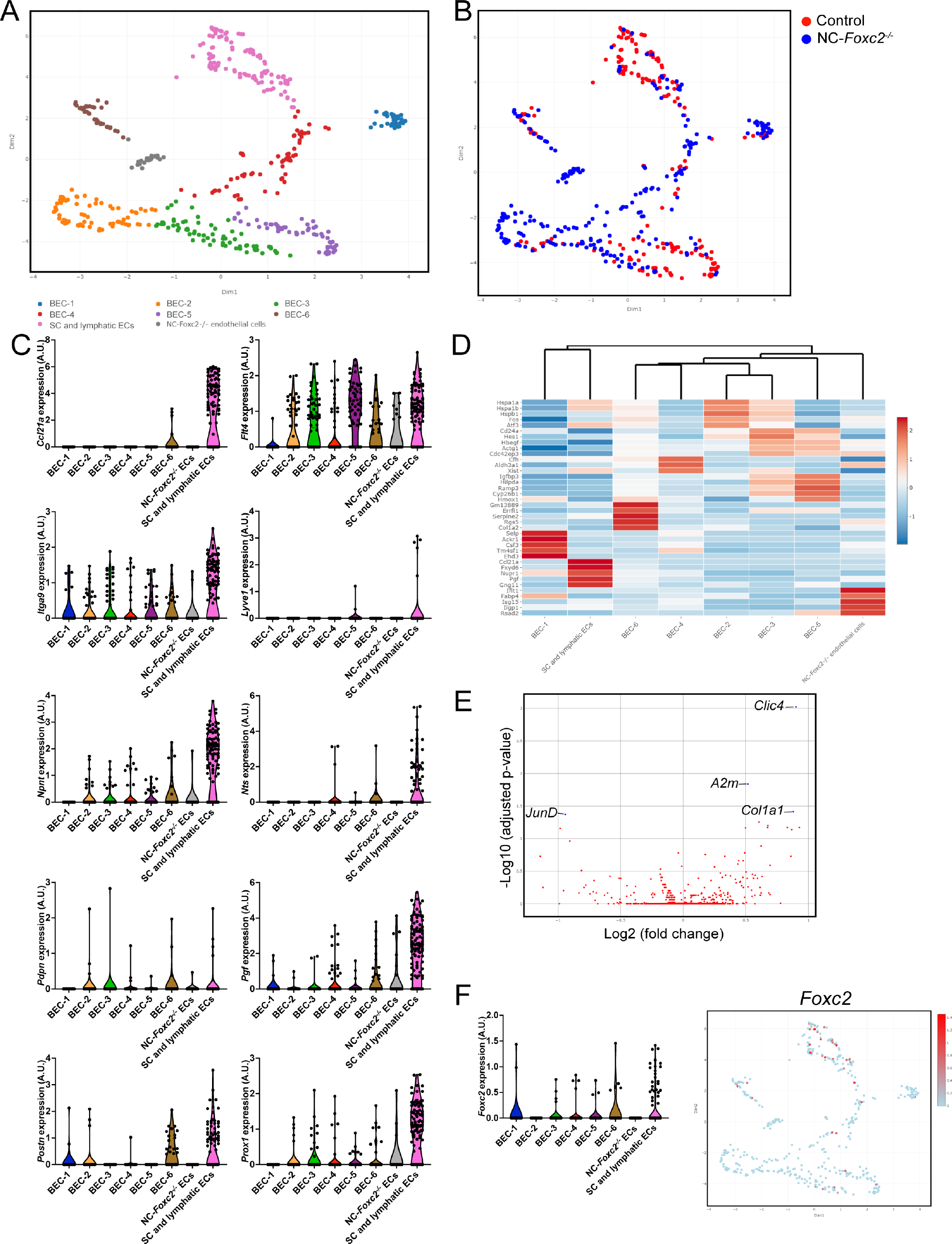
Single-cell transcriptome profiling of endothelium in the anterior eye segment of Control and NC-*Foxc2^-/-^* mice. (**A, B**) Visualization of unsupervised subclustering analysis of 8 transcriptionally distinct endothelial cell populations (**A**) and individual cell contribution (**B**) by uniform manifold approximation and projection (UMAP) of the endothelial cell cluster identified in Fig. 4B. (**C**) Violin plots showing the expression of gene markers from the endothelial subclusters. (**D**) Feature heatmap and hierarchical clustering of gene-expression levels of the top 5 marker genes for each endothelial subcluster. Colors represent row-wise scaled gene expression with a mean of 0 and SD of 1 (Z scores). (**E**) Volcano plot showing DEGs between NC-*Foxc2^-/-^* and *Foxc2^fl/fl^* Control mice in the Lymphatic-like endothelium subcluster. Blue points denote DEGs with adjusted p-value < .05. (**F**) Violin plot (left panel) and UMAP projection (right panel) showing the expression of *Foxc2* in each endothelial subcluster.

Analysis of DEGs in the SC and lymphatic EC cluster identified several transcriptional changes including significantly decreased expression of the AP-1 transcription factor component *JunD*, which has been previously implicated in the regulation of vascular injury response (49) and protection against aging-induced oxidative stress and endothelial dysfunction (50). In contrast, expression of *Col1a1*, which was shown to be increased in both normal and glaucomatous SC cells in response to substrate stiffness (51), was significantly upregulated in the SC and lymphatic EC cluster **(Figure 7E)**.

Collectively, these data demonstrate that loss of NC-*Foxc2* expression results in several transcriptional changes altering TM composition and the ECM environment that may be associated with increased matrix stiffening and reduction in TIE2 signaling, which have been associated with the progression of glaucoma.

### Endothelial cell-derived *Foxc2* transcriptional activity is required for SC morphogenesis via regulation of TIE2 expression

*Foxc2* is a critical regulator of early lymphatic development as well as lymphatic maturation, maintenance, and function (10, 11, 13, 14). Our linage tracing analysis demonstrated that *Foxc2*+ cell descendants contribute to the formation of the SC vasculature (**Figure 1**) and our single cell RNA-seq analysis shows that *Foxc2* was more highly expressed in the SC and lymphatic EC cluster compared to the other BEC clusters (**Figure 7F**). Moreover, Foxc2+/Prox1+ cell expression was previously observed in SC endothelium by immunostaining analysis as early as P7 continuing to 2 months of age (20). As the SC vasculature shares characteristics of lymphatic endothelium, we sought to investigate the direct role of endothelial-*Foxc2* signaling in SC morphogenesis as its role is unknown (**Figure 8**). Because early, inducible postnatal blood and/or lymphatic endothelial-specific deletion of *Foxc2* results in lymphatic dysfunction and eventual mortality (10, 14), we performed the analysis at the approximate midpoint of morphogenesis (P7) following the administration of tamoxifen from P1-P5 to delete *Foxc2* during SC morphogenesis initiation. Compared to *Foxc2^fl/fl^* controls, inducible, endothelial-specific deletion of *Foxc2* (*Cdh5-Cre^ERT2^; Foxc2^fl/fl^*, EC- *Foxc2*-KO) resulted in a hypoplastic SC vasculature (**Figure 8, A – F**) with significantly reduced SC area (**Figure 8G**), in addition to markedly impaired lymphatic valve development and maturation which we previously reported (47). While EC-*Foxc2*-KO mice maintained expression of PROX1 and VEGFR-3 (**Figure 8, B and D**), reduced TIE2 expression was detected in the SC vasculature compared to *Foxc2^fl/fl^* controls (**Figure 8F**). As *Foxc1* and *Foxc2* share cooperative roles in lymphatic development and maintenance (13, 14), we sought to also characterize possible roles for endothelial- *Foxc1* transcriptional signaling during SC morphogenesis (**Figure 8-figure supplement 1**). Compared to Cre-negative *Foxc1^fl/fl^* controls, *Cdh5-Cre^ERT2^; Foxc1^fl/fl^* (EC-*Foxc1*-KO) mice trended toward a reduction in SC area, but the difference was not statistically significant (**Figure 8-figure supplement 1G**). Additionally, EC*-*Foxc1-KO mice maintained expression of PROX1, VEGFR-3, and TIE2 (**Figure 8-figure supplement 1, A – F**). Thus, similar to their individual roles in the mesenteric lymphatic vasculature (14), SC endothelial morphogenesis is predominately regulated by *Foxc2* compared to its closely related family member *Foxc1*.

**Figure 8.**
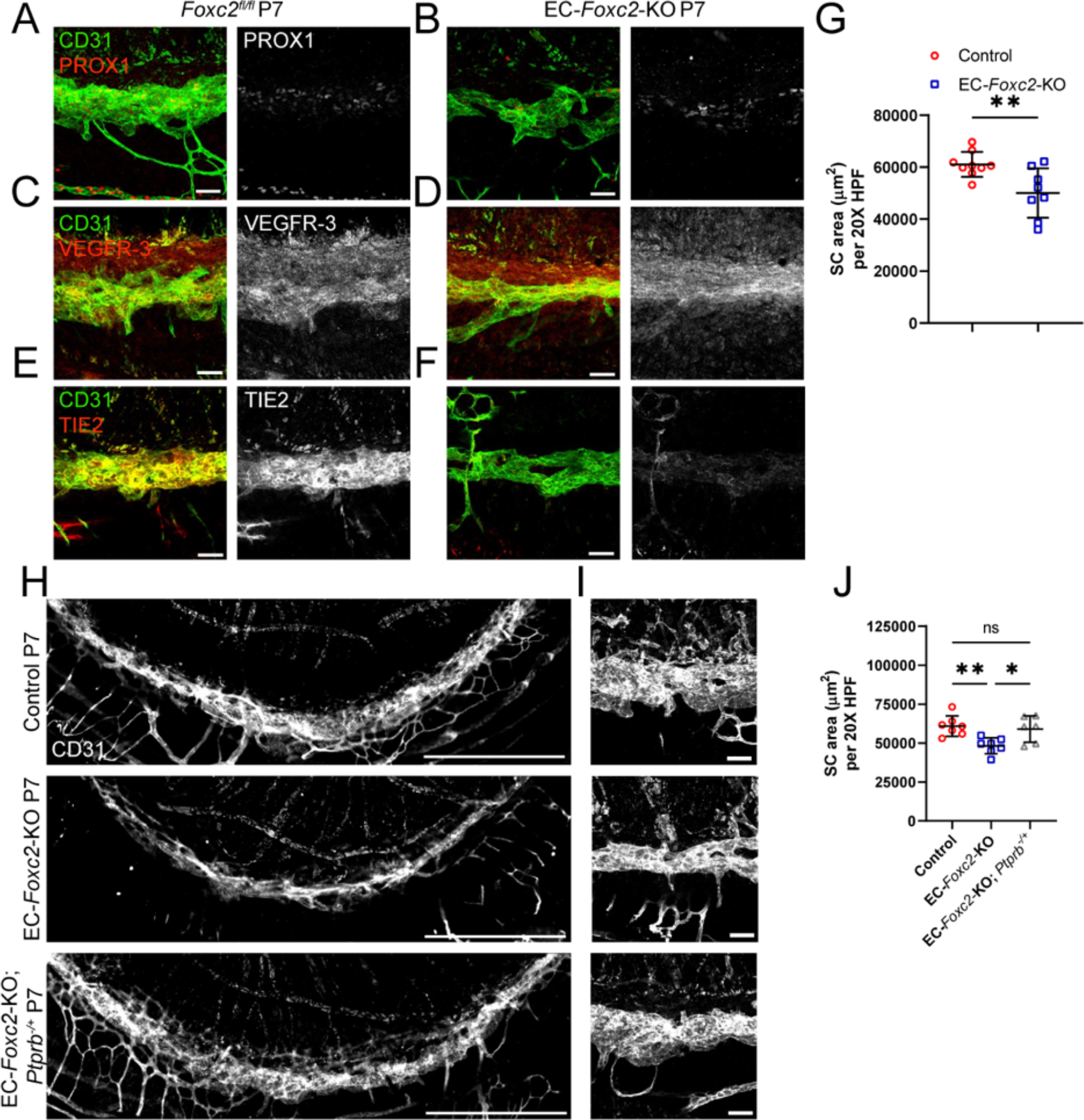
SC morphogenesis is impaired by early postnatal, endothelial-specific deletion of *Foxc2*, which is accompanied by reduced TIE2 expression and rescued by reduction of endothelial *Ptprb* expression. (A–F) Representative images of CD31 and PROX1 (A, B), VEGFR-3 (C, D), or Tie2 (E, F) expression in the SC of P7 *Foxc2^fl/fl^* control (A, C, E) and EC-*Foxc2*-KO mice (B, D, F). Scale bars are 50 μm. (G) Quantification of SC area per 20X high-power field in P7 *Foxc2^fl/fl^* control and EC-*Foxc2*- KO mice. N = 9 for Control and N = 8 for EC-*Foxc2*-KO mice. Data are mean ± SD. Statistical analysis: Student’s unpaired t-test. ** P < .01. (H, I) Representative images of SC vasculature immunostained with CD31 antibody in P7 Control, EC-*Foxc2*-KO, and EC-*Foxc2*-KO; *Ptprb^-/+^* mice. Scale bars are 500 μm (H) and 50 μm (I). (J) Quantification of SC area per 20X high-power field in P7 Control, EC-*Foxc2*-KO, and EC-*Foxc2*-KO; *Ptprb^-/+^* mice. N = 7 for Control, N = 7 for EC-*Foxc2*-KO, and N = 6 for EC-*Foxc2*-KO; *Ptprb^-/+^* mice. Statistical analysis: One-way ANOVA with Tukey’s multiple comparison’s test. * P < .05, ** P < .01.

*Ptprb* encodes receptor-type tyrosine-protein phosphatase beta, which is also known as VE-PTP. VE-PTP functions to negatively regulate ANGPT-TIE2 signaling by dephosphorylating the TIE2 receptor (52, 53). Notably, deletion of one *Ptprb* allele was able to rescue impaired SC morphogenesis and retinal ganglion cell (RGC) loss associated with *Tie2/Tek* haploinsufficiency (27) and a small molecule inhibitor of VE-PTP, AKB-9778, was shown to activate TIE2 signaling in the SC, increase outflow facility, and reduce IOP (28). Previous studies have also demonstrated that *FOXC2* directly binds to the *TIE2/TEK* locus (54) and that siRNA-mediated knockdown of *FOXC2* reduces TIE2 expression in cultured human dermal lymphatic endothelial cells (21). Therefore, to more directly assess the role of *Foxc2* in regulation of *Tie2/Tek* expression in the SC vasculature, we generated *Cdh5-Cre^ERT2^; Foxc2^fl/fl^; Ptprb^fl/+^* (EC- *Foxc2*-KO; *Ptprb^-/+^*) mice and performed conditional deletion postnatally from P1-P5. Compared to the development of a hypoplastic SC in postnatal EC-*Foxc2*-KO mice, EC- *Foxc2*-KO; *Ptprb^-/+^* individuals appeared to have normal SC vasculature (**Figure 8, H and I**) and SC area returned to levels similar to Cre-negative *Foxc2^fl/fl^* controls (**Figure 8J**). Thus, endothelial-*Foxc2* transcriptional activity regulates TIE2 expression to promote activation of ANGPT-TIE2 signaling during SC morphogenesis.

### *Foxc2* can functionally substitute for *Foxc1* during ocular development of the anterior segment

FOXC1 and FOXC2 share nearly identical forkhead DNA binding domains and function cooperatively during early cardiovascular (12) and ocular (15) development as well as during embryonic lymphangiogenensis (13) and postnatal lymphatic valve development and maintenance (14). However, mutations in *FOXC1* are predominately associated with the ocular autosomal dominant disorder Axenfeld-Rieger syndrome and progression of secondary glaucoma (55) and NC-*Foxc1^-/-^* mice are perinatal lethal (56, 57) compared to NC-*Foxc2^-/-^* mice, thus underscoring the critical role for NC-*Foxc1* function. We previously reported the generation of mice that carry a *Foxc2* knock-in allele (*Foxc1^c2^*) in which the *Foxc1* coding region has been replaced with the cDNA coding for *Foxc2* and that these mice appear to develop normally and do not exhibit abnormal development of the mesenteric lymphatic vasculature (14). To investigate whether the development of the anterior eye segment and SC morphogenesis is similarly conserved, we assessed ocular phenotypes in homozygous *Foxc1^c2/c2^* mice and WT (*Foxc1^+/+^)* controls by immunohistochemical analysis (**Figure 9**). At embryonic day ©15.5, development of the anterior chamber and cornea appeared morphologically similar to *Foxc1^+/+^* controls (**Figure 9A**). Analysis of 6-8-week old adult individuals showed that there were no obvious morphological differences in the SC between *Foxc1^+/+^*and *Foxc1^c2/c2^* and expression of key SC markers such as PROX1 and TIE2 were maintained (**Figure 9, B – E)**. Quantitative analysis of relative SC volume by vis-OCT and quantification of CD31 immunostained SC area did not identify significant differences between either group (**Figure 9, F and G**). Thus, these data demonstrate that like our group’s previous observations regarding lymphatic development and maturation, FOXC2 can functionally substitute for FOXC1 during anterior segment development.

**Figure 9.**
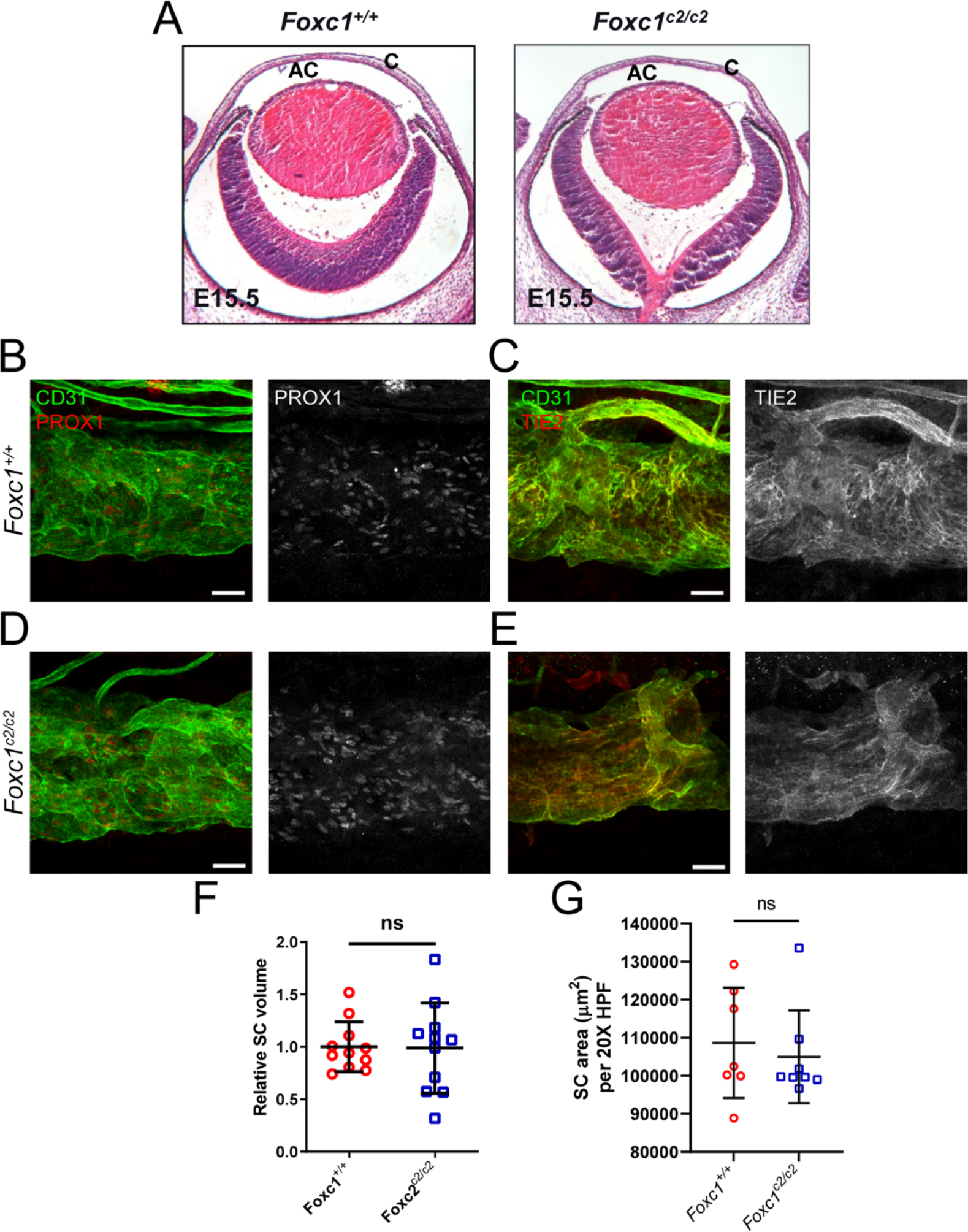
Substitution of *Foxc2* into the *Foxc1* locus does not impair anterior eye segment development nor SC morphogenesis. (**A**) Representative images of hematoxylin and eosin-stained transverse eye sections from embryonic day (E)15.5 *Foxc1^+/+^* and *Foxc1^c2/c2^* mice show no difference in the normal development of the anterior chamber. AC, anterior chamber, C, cornea. (**B-E**) Representative images of CD31 and PROX1 (**B, D**) or Tie2 (**C, E**) expression in the SC of adult *Foxc1^+/+^* (**B, C**) and *Foxc1^c2/c2^* mice (**D, E**). Scale bars are 50 μm. (**F**) Relative SC volumes of *Foxc1^c2/c2^* and *Foxc1^+/+^* mice in a 1.5 mm x 1.5 mm field of view. SC volume for both groups were normalized by mean *Foxc1^+/+^* SC volume. N = 11 volumes from 11 individuals for *Foxc1^+/+^* and N = 11 volumes from 11 individuals for *Foxc1^c2/c2^* mice. (**G**) Quantification of SC area per 20X high-power field (HPF). N = 7 for *Foxc1^+/+^* and N = 8 for *Foxc1^c2/c2^* mice. Data are mean ± SD. Statistical analysis: Student’s unpaired t-test.

## Discussion

Critical to the development of glaucoma is ocular hypertension resulting from abnormally increased IOP, which is tightly regulated by control of outflow facility in part through the conventional outflow pathway consisting of the TM and SC that drain into the ocular veinous circulation (6). While several recent studies have implicated the direct roles of key vascular signaling events contributing to the morphogenesis and functional maintenance of the SC during early postnatal development and adulthood (20, 21, 23–25), the role of paracrine signaling events from the TM and environmental cues contributing to proper morphogenesis of SC during early development is not well understood. In this study, we identify a critical role for NC-*Foxc2* in maintenance of SC endothelial identity and its proper morphogenesis. In contrast, we also demonstrate that endothelial-*Foxc2* is required for TIE2 expression in the SC, thus highlighting key differences of the functional role of FOXC2 transcriptional regulation in different tissues during ocular development (**Figure 10**).

**Figure 10.**
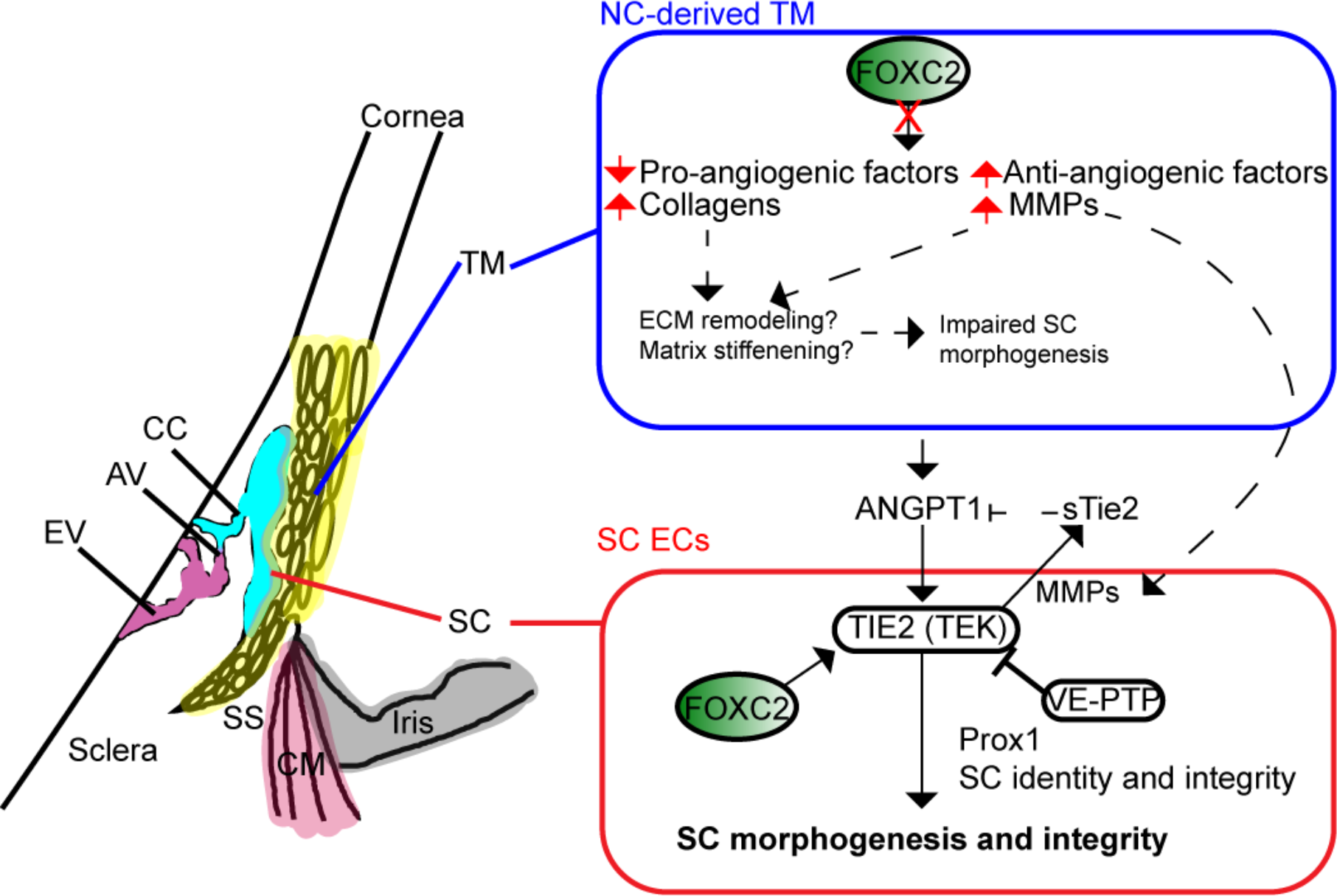
Neural crest- and endothelial- derived Foxc2 differentially regulate proper morphogenesis of the SC. Neural crest-specific deletion of *Foxc2* results in several transcriptional changes in the trabecular meshwork including the decrease in expression of pro-angiogenic factors and in contrast an increase in expression of anti-angiogenic factors, collagens, and MMPs that likely result in abnormal ECM remodeling, matrix stiffening, and induction of TIE2 shedding that potentially contribute to impaired SC morphogenesis. In contrast, endothelial derived Foxc2 regulates the expression of TIE2 to promote ANGPT1-mediated activation of the TIE2 receptor and in turn proper SC morphogenesis and maintenance of SC integrity. TM, trabecular meshwork. SC, Schlemm’s Canal. SS, scleral spur. CM, ciliary muscle. CC, collector channel. AV, aqueous veins. EV, episcleral veins.

Our group had previously reported that NC-specific deletion of *Foxc2* is associated with several anterior segment abnormalities, including corneal conjunctivalization, ectopic corneal neovascularization, and disrupted ocular epithelial cell identity (15), however the outcome on development and changes within the conventional outflow pathway, and in particular the SC, was not well elucidated. By applying an *in vivo* vis-OCT imaging technique (33) for the first time to characterize SC phenotypes in a transgenic knockout model of abnormal anterior segment development, we identified significant SC volume differences between *Foxc2^fl/fl^* control and NC-*Foxc2^-/--^* mice at every IOP level from Δ- 10 mmHg to Δ+5 mmHg **(Figure 2 I)**. Such observations indicate that there exists increased outflow resistance in SC in NC-*Foxc2^-/-^* mice at physiological IOPs. Further analysis quantifying the SC width and height revealed patterns suggesting increased TM stiffness in NC-*Foxc2^-/-^* mice compared with the *Foxc2^fl/fl^* control mice **(Figure 2, J and K)**. Indeed, it was noted that the height of the SC was smaller in NC-*Foxc2^-/-^* mice (**Figure 2, C and D, Figure 1-figure supplement 1C and D**). The height of the SC is primarily influenced by the pressure difference across the TM, with higher IOPs leading to reductions in height. At typical IOPs, the tension in the TM keeps the SC open, although at higher IOPs septae within the SC prevent the complete collapse of the SC (58). In our analysis, as IOP increased, both the measured height and width of the SC decreased in NC-*Foxc2^-/-^* mice, while decreases in only height were seen in *Foxc2^fl/fl^* mice (**Figure 2, J and K**). Notably, decreases in SC width are caused by compression of parts of the SC to heights smaller than the resolution of our vis-OCT system, making it impossible to visualize these portions of the SC.

In support of a change in ECM composition potentially leading to increased TM stiffness, our scRNA-Seq analysis between *Foxc2^fl/fl^* and NC-*Foxc2^-/-^* mice identified several differentially expressed genes which provide additional evidence of differences in ECM matrix composition in the TM. For example, many of the differentially expressed genes upregulated in NC-*Foxc2^-/-^* are involved in extracellular matrix organization and collagen chain trimerization (**Figure 5)**. While the scRNA-Seq data does not directly prove increased TM stiffness in NC-*Foxc2^-/-^* mice, there is substantial evidence of differences in the extracellular matrix organization leading to altered TM elasticity in NC-

*Foxc2^-/-^* mice as is illustrated by the trends in SC height and width. While changes in SC area, width, and height with IOP changes provide clues related to TM stiffness (59, 60), further work is required to numerically quantify the stiffness. Given the relevance of TM stiffness in glaucoma development and the observation of elevated IOP in NC-*Foxc2^-/-^* mice, further studies focused on the role of Foxc2 on TM stiffness are warranted.

Additionally, it is important to consider whether the elevations in IOP we observe in NC- *Foxc2^-/-^* mice are associated with loss of RGCs and the progression of glaucomatous neuropathy as previously demonstrated (61, 62). Characterization of the retina nerve fiber layer (RNFL) of NC-*Foxc2^-/-^* mice demonstrated that it was thinner compared to *Foxc2^fl/fl^* controls (**Figure 10-figure supplement 1**), consistent with other models of impaired SC morphogenesis leading to glaucomatous neuropathy (24). However, it is important to consider that this phenotype may be secondary to potential ocular development defects present in the posterior segment of NC-*Foxc2^-/-^* individuals, thus warranting further investigation.

Additional changes in ECM matrix composition and signaling may be attributable to impaired SC morphogenesis observed in NC-*Foxc2^-/-^* mice as well. Changes in ECM composition and matrix stiffness have dynamic effects on vascular signaling (63) and increases in collagen density impaired neovessel length and interconnectivity in an *in vitro* model of sprouting angiogenesis (64). Similarly, recent evidence has shown that reduced matrix stiffness primes lymphatic endothelial cells to form cord-like structures and increase the expression of lymphatic markers such as LYVE-1 and PROX1 *in vitro* in response to VEGF-C (65) and promote increased GATA2 expression and GATA2- dependent upregulation of genes involved in cell migration and lymphangiogenesis (66). Notably, we show increased expression of MMP-2 in tissues of the anterior segment of NC-*Foxc2^-/-^* mice (**Figure 6**). Evidence has shown that the TM increases secretion of MMP-2, -3, and -14 in response to IOP elevation and mechanical stretching forces, although these changes were not accompanied by increases in their mRNA levels, but likely by selective translation mediated by mammalian target of rapamycin (mTOR) (67). However, it is possible that increased expression of genes associated with ECM remodeling may be in response to abnormal development of the anterior segment as opposed to other mechanisms generating outflow resistance. While increased expression of MMP-2 and MMP-14 is associated with stimulating angiogenesis via ECM turnover (68), the increased expression of MMPs may have a negative impact on SC morphogenesis. Several studies have demonstrated that MMPs mediate cleavage and shedding of the TIE2 ectodomain to produce sTIE2, which acts an endogenous inhibitor by trapping ANGPT1 (29, 46). Provided the critical role of ANGPT/TIE2 signaling in SC morphogenesis and maintenance and our observations that 1) TIE2 expression is reduced in the SC endothelium of NC-*Foxc2^-/-^* mice and 2) MMP activity mediates TIE2 shedding in cultured HDLECs, it is a reasonable to speculate that sTIE2 is increased in NC-*Foxc2^-/-^* mice. Thus, it is probable that activation of ANGPT/TIE2 signaling is reduced, resulting in loss of SC identity establishment (**Figure 10**). However, additional investigation is needed to validate this mechanism and whether changes in ECM remodeling or activation of TIE2 is directly associated with the changes we observe in the SC and lymphatic-like cells of the anterior segment such as reduction of *JunD* and increase of *Col1a1* (**Figure 7E**).

Our scRNA-seq analysis also shows a significant reduction of several pro-angiogenic factors in the TM (**Figure 5-figure supplement 1, Supplementary Table 1**), including *Vegfa*. We previously reported that loss of NC-*Foxc1* is associated with increased MMP signaling and VEGF-A bioavailability in the corneal stroma (56). Notably, we also observed a significant increase in *Vegfa* mRNA expression within the corneal stroma keratocyte population in our scRNA-seq analysis (**Supplementary Table 1**). As SC sprouting morphogenesis is not initiated from the limbal lymphatic vasculature and instead from the blood limbal vascular plexus and radial vessels, which requires VEGFR-2 function (19, 22), it is possible that the severe cornea neovascularization and subsequent impairment in SC morphogenesis may be attributable to alterations in VEGF-A bioavailability and alterations in VEGF gradient signaling in NC-*Foxc2^-/-^* mice.

In contrast to the role of NC-*Foxc2* expression in the development of anterior segment tissues, our results demonstrate that endothelial-*Foxc2* is necessary for regulation of TIE2 expression in SC endothelium (**Figure 8**) and support previous findings identifying a role for FOXC2 in the regulation of TIE2 expression in both blood and lymphatic endothelium (21, 54). While the significance of maintenance of ANGPT/TIE2 signaling activity in the SC is well understood, it is plausible that FOXC2 has other critical functional roles in the SC endothelium as well. Loss of *Foxc2* expression in lymphatic collecting vessels perturbs “zipper-like” cell-cell junctions and increases vessel leakiness (10, 14). Like lymphatic collecting vessels, the SC vasculature also contains “zipper-like” cell-cell junctions (19, 20). Whether *Foxc2* regulates cell-cell junctions in the SC endothelium similar to other lymphatic vascular beds remains to be well understood and will be the focus of further investigation.

Together, the data presented in this report present a unique bi-functional role for *Foxc2* in both neural crest- and endothelial-derived cells in regulating the development of the anterior eye segment to promote proper morphogenesis of the SC. Given that our evidence also demonstrates that *Foxc2* can functionally substitute for *Foxc1* during ocular morphogenesis (**Figure 9**), this work may also offer insight into pathological signaling mechanisms associated with *FOXC1* mutations in Axenfeld-Rieger Syndrome and the progression of secondary glaucoma to identify novel therapeutic targets.

## Materials and Methods

### Animal Generation and Husbandry

Mice were housed and kept under normal lighting conditions with 12-hour-on, 12-hour- off cycles in the Center for Comparative Medicine at Northwestern University. *Wnt1-Cre; Foxc2^fl/fl^* (NC-*Foxc2*-KO) mice were generated as reported previously (15). Endothelial specific *Foxc1* and *Foxc2* knockout mice were generated and induced with tamoxifen dissolved in corn oil as previously described (14). Briefly, neonates were orally administered 75 μg of tamoxifen dissolved in corn oil from postnatal day (P)1 – P5 to induce gene deletion and mice were euthanized at the indicated time points for analysis. For cell fate mapping, *Wnt1-Cre; Foxc2^fl/fl^* and *Foxc2-Cre^ERT2^* mice (30) (a gift from Kazushi Aoto, Hamamatsu University School of Medicine) were crossed with mTmG reporter mice (The Jackson Laboratory). *Cdh5-Cre^ERT2^; Foxc2^fl/fl^; Ptprb^fl/+^* (EC-*Foxc2*- KO; *Ptprb^+/-^*) mice were generated by crossing *Ptprb^fl/fl^* mice (69), acquired from Northwestern University’s NU GoKidney Preclinical Models Core, with *Cdh5-Cre^ERT2^; Foxc2^fl/fl^* mice through several generations. *Foxc2* knock-in mice (*Foxc1^c2/c2^*) were generated as described previously (14). Genotyping of mice for use in analysis was performed by Transnetyx Inc (Cordova, TN) using real-time PCR. All experimental protocols and procedures used in this study were approved by the Institutional Animal Care and Use Committee (IACUC) at Northwestern University.

### *In Vivo* imaging of Schlemm’s Canal using Visible-Light OCT

*In vivo* imaging of the mouse Schlemm’s canal (SC) was performed using a custom-built anterior segment vis-OCT microscopy system as previously described (33). Briefly, mice were anesthetized by intraperitoneal injection (10 mL/kg bodyweight) of ketamine xylazine cocktail (ketamine: 11.45 mg/L; xylazine: 1.7 mg/mL, in saline) prior to imaging procedures. IOP was measured before imaging and after deep anesthesia using the TonoLab rebound tonometer (Colonial Medical Supply). The probe tip was positioned about 2-3mm away from the central cornea in alignment with the optical axis of the eye. The measured IOP value was taken as the average of six consecutive measurements.

During imaging, body temperature was maintained by a heating lamp. The entire 360 degrees of the SC and surrounding vasculature was captured in 8 separate raster scans, with a rotating two-mirror assembly used to change the field of view between scans as previously described (33). Each raster scan had a 1.8mm x 1.8mm field of view. The resolutions of the system in tissue are 7 μm laterally and 1.3 µm axially. Vis-OCTA detecting motion contrast from flowing blood cells was used to visualize the surrounding vasculature, with each B-scan repeated 5 times and processed as previously described (70).

To assess changes in SC volume in response to alterations in intraocular pressure (IOP), the anterior chamber was cannulated with a 34-gauge needle, and the IOP level was manometrically set prior to acquiring a vis-OCT raster image as previously described (33). Only the nasal most raster scan was used for volume calculation. One vis-OCT dataset was acquired at each of 5 IOP levels, and each IOP level was repeated 3 times. A 1.5 mm length of the SC was segmented from each individual raster scan. All SC volumes are reported as volumes normalized by the average control mouse volume at baseline IOP. For calculation of SC width and height, an ellipse was fitted to the segmented SC in every segmented cross-sectional B-scan image using the *regionprops* function in MATLAB 2020b (MathWorks, Natick, MA, USA). In cases where the SC was composed of multiple parts, an ellipse was fitted to every part of the SC with an area at least 20% that of the largest SC area. SC width was calculated by summing the major axis length of the fitted ellipses within each cross-section and averaging the value across all cross-sections. SC height was calculated by taking the weighted average by area of the minor axis length of the fitted ellipses and averaging across all cross-sections.

### Tissue section processing, histologic and immunohistochemical analysis

Whole embryos were fixed in 4% paraformaldehyde (PFA) for 2 hours at 4°C, dehydrated with methanol, embedded in paraffin, and cut into 8 μm sections. Adult eyeballs were immersion fixed in 4% PFA overnight at 4°C, followed by immersion in 30% sucrose (wt/vol in PBS) overnight at 4°C. Tissues were then embedded in OCT Compound (Tissue-Tek) and frozen in an ethanol/dry ice bath. Frozen tissues were cut into 8 μm sections. Both paraffin sections from whole embryos and frozen sections from adult eyeballs were stained with hematoxylin and eosin (H&E) and subject to immunohistochemistry. H&E staining images were acquired using an Olympus Vanox AHBT3 Research Microscope using both a 4X or 40X objective. For immunohistochemical analysis of markers from frozen sections, sections were blocked with 10% normal donkey serum (Sigma-Aldrich Corp.) in PBS with 0.1% Triton X-100 (Sigma-Aldrich Corp.). Following blocking, sections were then incubated in blocking buffer with primary antibodies listed in **Table S2** overnight at 4°C. Sections were then washed with PBS/0.1% Tween 20 and incubated with secondary antibodies conjugated to AlexaFluor 488, Alexafluor 568, or AlexaFluor 647 listed in **Table S2** for 2 hours at room temperature. Wash steps were repeated, then the sections were counterstained with 4’6-diamidino-2-phenylindole (DAPI), and mounted with PermaFluor aqueous mounting media (Thermo Fisher). Images of the iridocorneal angle were captured on a Nikon A1R confocal microscope at the Northwestern University Center for Advanced Microscopy using a 20X objective with a numerical aperture of 0.75 and a pinhole of 1.2 Airy units to collect Z-stacks. Images were acquired using Nikon NIS-elements software and are shown as maximum intensity projections, which were generated using Fiji software and were post-processed using Adobe Photoshop. Large-field images of eyes were obtained by stitching images captured using a 10x objective with a numerical aperture of 0.3 and pinhole of 1.2 Airy units. Images were post-processed using Adobe Photoshop.

### Wholemount immunostaining analysis of SC morphology

Mice were euthanized at the indicated time points, eyes were enucleated and then immersion fixed in 2% PFA overnight at 4°C. Eyes were then bisected from the optic nerve to the center of the cornea and the lens and retina tissue were removed. Tissues were then blocked in buffer containing 5% donkey serum, 2.5% Bovine Serum Albumin (BSA, Sigma-Aldrich Corp.), 0.5% Triton X-100 in TBS pH 7.4 overnight at 4°C on a shaker. Following blocking, the tissues were incubated with primary antibodies listed in **Table S2** diluted in blocking buffer overnight at 4°C on a shaker. Tissues were then washed in 0.05% Tris-buffered Tween-20 (TBST) solution then incubated with secondary antibodies listed in **Table S2** diluted in blocking buffer overnight at 4°C on a shaker. Wash steps were repeated, then the tissues were further processed by making additional cuts in the cornea and scleral regions to flat-mount tissues on glass microscope slides with PermaFluor aqueous mounting medium to visualize the Schlemm’s Canal vasculature. Flatmounted tissues were imaged using a Nikon A1R confocal microscope at the Northwestern University Center for Advanced Microscopy. Images of SC vasculature were captured using a 20X objective with numerical aperture of 0.75 and a pinhole of 1.2 Airy units to collect Z-stacks. Images were acquired using Nikon NIS-elements software and are shown as maximum intensity projections, which were generated using Fiji software and were post-processed using Adobe Photoshop. Large-field images of SC morphology were obtained by stitching images captured using the same 20x objective and a fully opened 150 μm pinhole. Images were post-processed using Adobe Photoshop.

### Preparation of single-cell suspension from mouse anterior eye segment for single cell RNA-sequencing analysis

For single-cell RNA sequencing, eyes were pooled from 4 – 6 individuals of both 3 – 4 week old *Foxc2^fl/fl^* control and NC-*Foxc2^-/-^* mice. The anterior segment from each eye was dissected in ice-cold dye-free DMEM containing 10% FBS and the iris was gently removed using fine forceps. Pooled tissues from each group were then chopped with Vannas Scissors in ice-cold DMEM, then digested in DMEM containing 10% FBS, 1 mg/mL Collagenase A (Millipore Sigma), and 10 μM Y-27632 (R&D Systems) for 2 hours at 37°C. Tissues were then washed in 1X PBS solution, then further digested in 0.25% Trypsin solution containing 10 μM Y-27632 and 0.2 mg/mL DNAse I (Sigma) at 37°C for 25 minutes with shaking and gentle trituration using a P1000 pipettor and wide-bore pipette tips. Following dissociation, the tissues were centrifuged and the supernatant was removed. Pelleted cells were washed and re-suspended in warm DMEM containing 10% FBS, then passed through a Flowmi 40 μm cell strainer before repeating centrifugation. Washing and centrifugation was repeated once more to pellet cells, which were then resuspended in 100 μL of ice-cold Hank’s balanced salt solution (HBSS) containing 1% BSA. Cell viability was assessed using the Cellometer Auto 2000 Cell Viability Counter. Cell viability of 70% was used as a minimum threshold.

### Single-cell 3’ gene expression library construction and sequencing

Single cell 3’ gene expression libraries were constructed from samples using the Chromium Next GEM Single Cell 3’ Reagent Kits v3.1 (10x Genomics, Pleasonton, CA, USA) according to the manufacturer’s instructions. Libraries were then assessed for quality (TapeStation 4200, Agilent, Santa Clara, CA, USA) and then processed for paired-end sequencing on an Illumina HiSeq 4000 platform (Illumina, San Diego, CA, USA). 10,000 cells were targeted for each sample with a sequencing depth of 20,000 read pairs per cell.

### Pre-processing of single-cell RNA data

Following library construction and sequencing, raw sequencing data were de- multiplexed and mapped to the mouse reference genome (mm10) using the CellRanger toolkit (10X Genomics, version 4.0.0). Gene expression matrices were then generated from *Foxc2^fl/fl^* control and NC-*Foxc2^-/-^* mice. The matrix files were then utilized for data processing and downstream analysis using the BIOMEX browser-based software platform and its incorporated packages developed in R (71). Quality control and data pretreatment was performed in BIOMEX with the following manually set parameters: i) genes with a row average of <.005 were excluded for downstream analysis and ii) cells in which over 8% of unique molecular identifiers (UMIs) were derived from the mitochondrial genome were considered as dead cells and removed from downstream analysis. The data were then normalized in BIOMEX using similar methodology to the *NormalizeData* function as implemented in the *Seurat* package (72).

### Variable gene identification, dimensionality reduction, clustering analysis, and differential gene expression analysis

Following data pretreatment, BIOMEX was utilized for downstream dimensionality reduction of data and clustering analysis using the incorporated R packages. First, highly variable genes (HVGs) were identified utilizing the following feature selections: mean lower threshold = 0.01, mean higher threshold = 8, dispersion threshold = 0.5. Data (using highly variable genes only) was then auto-scaled and summarized by principal component analysis (PCA), followed by visualization using t-distributed stochastic neighbor embedding (t-SNE; top 15 principal components (PCs)) to reduce the data into a two-dimensional space. Graph-based clustering was then performed in BIOMEX to cluster cells according to their respective gene expression profile using methodology similar to the *FindClusters* function in *Seurat* (clustering resolution = 0.8, k-nearest neighbors = 25). Clusters formed by doublets were then removed before further analysis. Clusters containing doublets could be identified by high expression levels of marker genes characteristic of several cell types as well as the lack of uniquely expressed genes.

For analysis of specific endothelial cell types, dimensionality reduction was repeated on the endothelial cell cluster, characterized by high expression levels of *Pecam1* and *Cdh5*, through utilization of PCA on identified HVGs (mean lower threshold = 0.01, mean higher threshold = 8, dispersion threshold = 0.5) followed by Uniform Manifold Approximation and Projection (UMAP). Graph-based clustering was then repeated in BIOMEX, using cluster resolution = 0.8 and k-nearest neighbors = 25.

Marker set analysis was performed in BIOMEX on HVGs to identify gene markers highly expressed in each initial cluster using similar methodology described previously (73).

Marker genes were then compared with previously reported single cell RNA-seq data characterizing the tissues of the anterior eye-segment and aqueous humor outflow pathway (36–39) to identify unique cell populations. Clusters with highly similar expression patterns indicative of the same cell phenotype were merged into the same cluster. Marker set analysis was then repeated on characterized cell clusters to identify top marker genes, which were utilized for generation of heatmap visualization.

Differential gene expression analysis between Control and NC-*Foxc2^-/-^* mice for individual cell clusters was performed in BIOMEX using the Model-based Analysis of Single-cell Transcriptomics (MAST) package.

### Single-cell RNA-seq data visualization

BIOMEX implementation of *Plotly* package was used for t-SNE and UMAP visualization. BIOMEX implementation of the *Heatmaply* package was utilized for heatmap visualization. Heatmaps were based on cluster-averaged gene expression and data was autoscaled for visualization.

### Cell Culture, GM-6001 administration, and ELISA

Primary HDLECs were isolated from neonatal human foreskins as described previously (74) and cultured from passages 5 – 7 on fibronectin-coated plates with EGM-2 MV growth media (Lonza) supplemented with human VEGF-C (R&D). Cells were cultured in 6-well plates until confluent, washed with cold PBS three times and then supplemented with serum-free DMEM in the presence of DMSO or the MMP inhibitor GM-6001 (Tocris Biosciences) at 100 nM or 5 μM concentrations. Cells were cultured overnight prior to collecting conditioned media that was filtered through a 0.22-μm pore membrane (Millipore) and treated with 1 mM sodium orthovanadate and Roche Complete Protease Inhibitor Cocktail tablets (according to the manufacturer’s instructions). Quantification of the concentration of sTIE2 in conditioned media was performed by ELISA using a human TIE-2 Quantikine ELISA kit (R&D) following instructions from the manufacturer.

### Quantification and Statistical analysis

Statistical analyses were performed using GraphPad Prism 9. For quantification of SC area, 4-8 20X high-power fields (HPF) were acquired per individual and the CD31+ area was measured and averaged. For comparisons of average measurements between two groups, Student’s two-tailed, unpaired t-test was used. For comparisons of measurements between more than two groups, one-way ANOVA was used with Tukey’s multiple comparisons test. For comparisons of mean values between two groups of mice at multiple IOP levels, two-way ANOVA was used with Šídák’s multiple comparisons test. For quantification of TIE2 expression, 3-8 HPF were acquired per individual and mean TIE2 levels were assessed using ImageJ. Statistical analysis was then performed using a Nested t-test. Statistical analysis for comparison of gene expression between *Foxc2^fl/fl^* and NC-*Foxc2^-/-^* in scRNA-seq violin plot datasets was completed using a Mann-Whitney test. For quantification of sTIE2 concentration, 3 biological replicates were measured by ELISA. Statistical analysis was then performed using a Kruskal-Wallis test with Dunn’s multiple comparisons. Pathway enrichment analysis on differentially expressed genes was performed using Metascape (75) on the KEGG, Canonical Pathways, GO, Reactome, and CORUM databases. P < 0.05 was determined to be statistically significant.

## Acknowledgments

We thank Drs. Mark Johnson (Northwestern University) and Benjamin R. Thomson (Northwestern University) for helpful advice. *Cdh5-Cre^ERT2^* mice were provided by Dr. Ralf Adams at the Max-Planck-Institute for Molecular Biomedicine, Germany. *Wnt1-Cre* mice were provided by Dr. Andrew McMahon and HDLECs were provided by Dr. Young-Kwon Hong, both at the Keck School of Medicine of the University of Southern California, USA. scRNA-seq experiments were performed at the NUSeq Core Facility at Northwestern University. This work was supported by NIH (R01HL144129 and R01EY028304 to TK, R01EY026078, R01EY029121, and U01EY033001 to HFZ and 5T32HL094293 to PRN). Confocal imaging work was performed at the Northwestern University Center for Advanced Microscopy supported by NCI CCSG P30 CA060553 awarded to the Robert H Lurie Comprehensive Cancer Center.

## Supplementary materials

**Figure 1-figure supplement 1.**
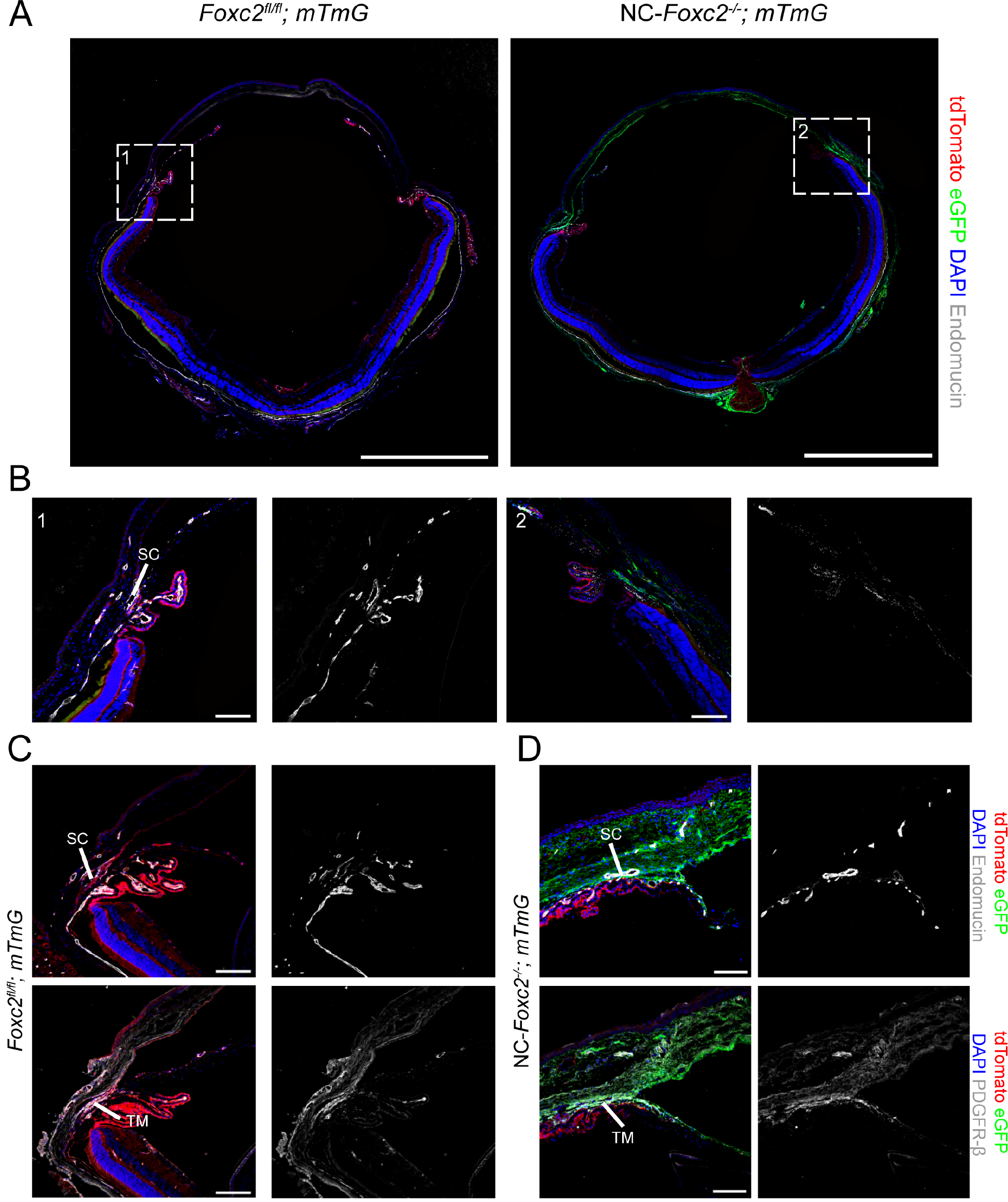
Fate-mapping of neural crest-derived cell populations in NC-*Foxc2^-/-^* mice. (**A**) Representative images of eye sections from 3- week-old control *Foxc2^fl/fl^; mTmG* and NC-*Foxc2^-/-^*; mTmG mice immunostained with antibody against endomucin where eGFP expression is regulated by *Wnt1-Cre*- mediated recombination. Scale bars are 1 mm. (**B**) Magnified images of boxed regions denoted in (**A**). Scale bars are 100 μm. (**C, D**) Representative images of sections of the iridocorneal angle from a 3-week-old NC-*Foxc2^-/-^; mTmG* individual with a severe phenotype (**D**) compared to a control *Foxc2^fl/fl^*; *mTmG* individual immunostained with antibody against endomucin or PDGFR-β. Scale bars are 100 μm.

**Figure 4-figure supplement 1.**
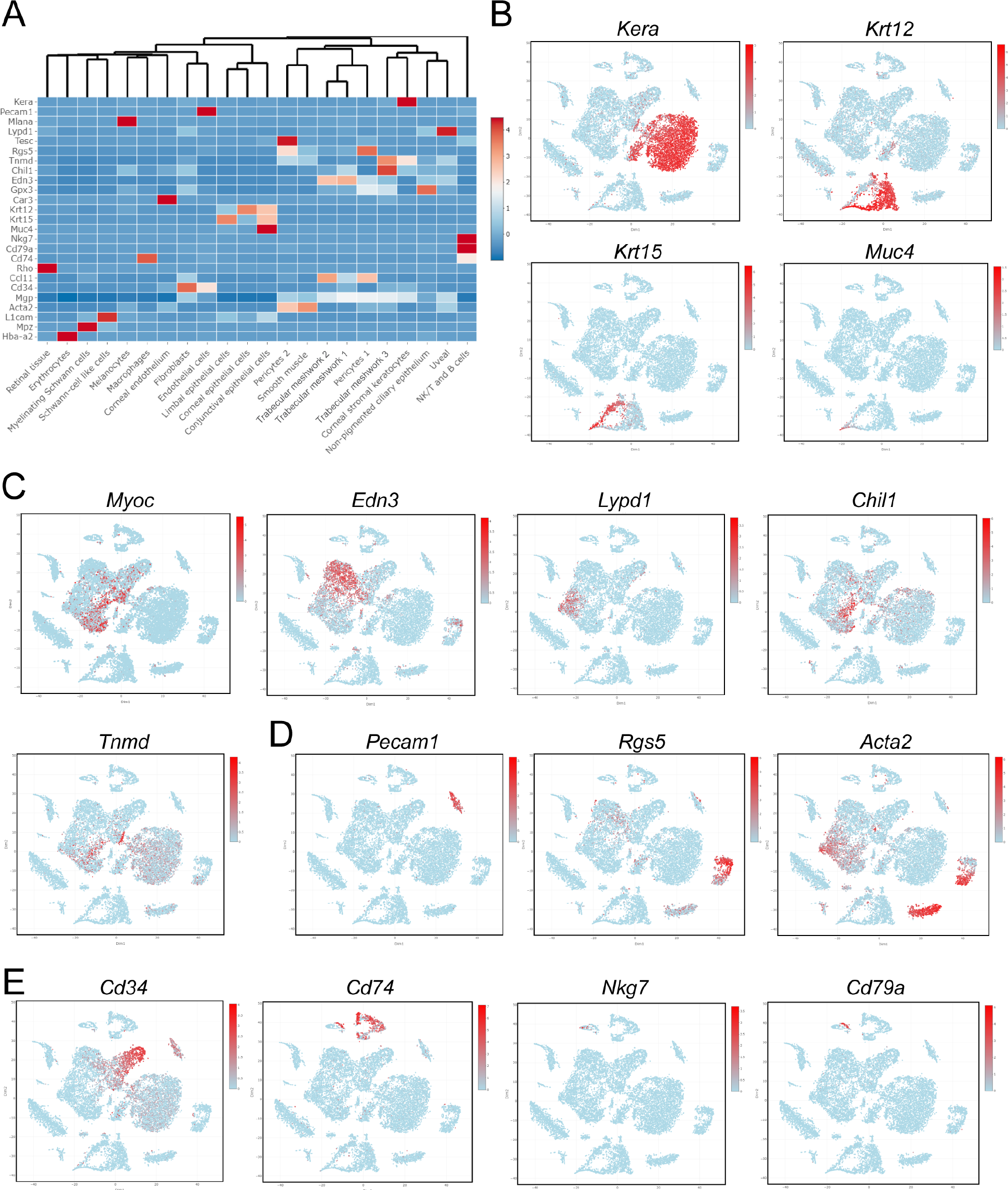
Characterization of cluster phenotypes. (**A**) Heatmap and hierarchical clustering of cell populations based on selective genes uniquely expressed in each cell population. (**B-E**) t-SNE plots colorized by the Seurat normalized expression of *Kera, Krt12, Krt15*, and *Muc4* identifying corneal stromal keratocytes and epithelial cell populations (**B**), *Myoc, Edn3, Lypd1, Chil1,* and *Tnmd* identifying trabecular meshwork cell populations (**C**), *Pecam1, Rgs5,* and *Acta2* identifying endothelial, pericyte, and smooth muscle cell populations (**D**), or *Cd34, Cd74, Nkg7*, and *Cd79a* identifying Fibroblast, macrophage, NK/T and B cell populations (**E**).

**Figure 4 – figure supplement 2.**
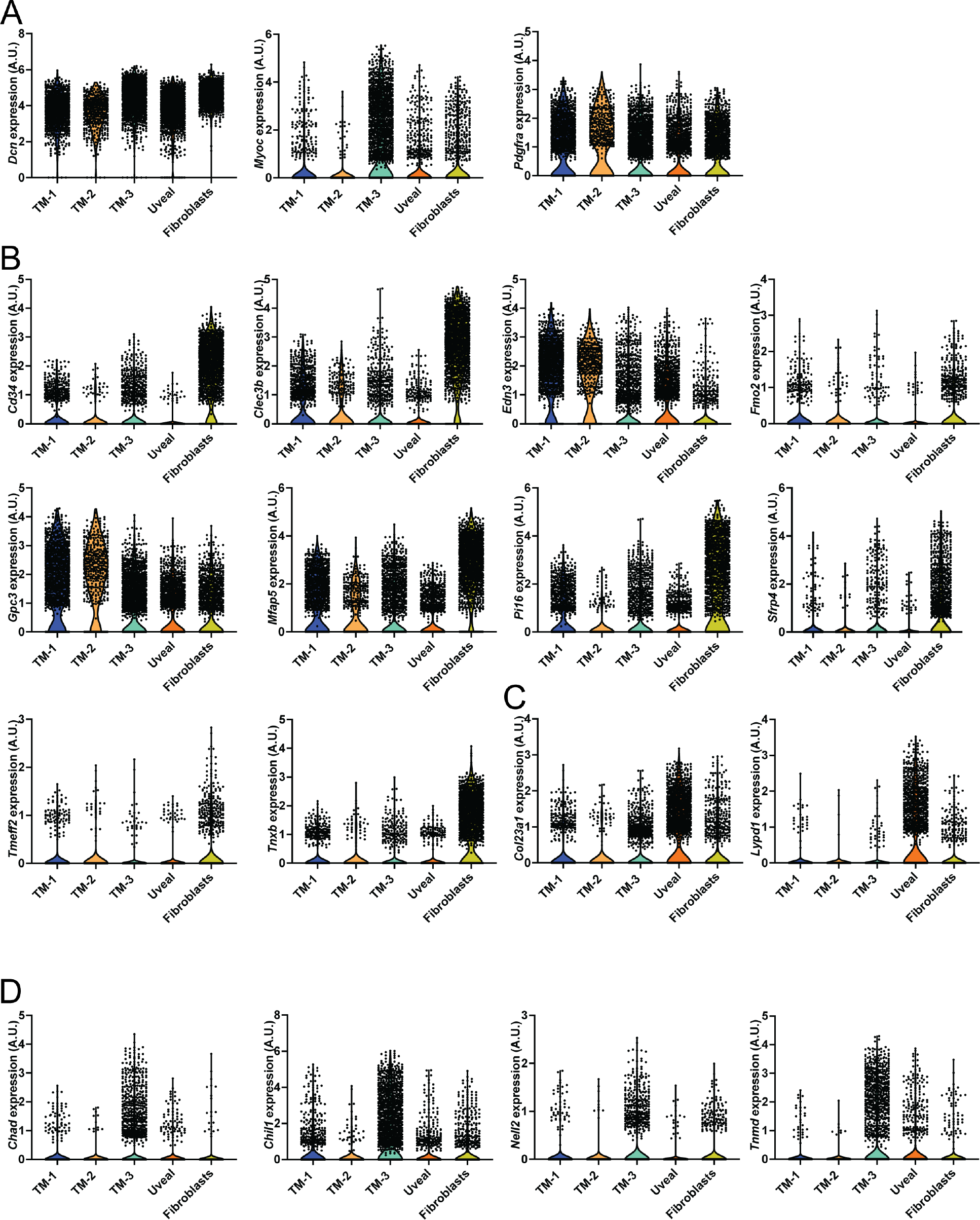
Characterization of trabecular meshwork clusters. (**A – D**) Violin plots showing the expression of *Dcn, Myoc,* and *Pdgfra* (**A**), *Cd34, Clec3b, Edn3, Fmo2, Gpc3, Mfap5, Pi16, Sfrp4, Tmeff2,* and *Tnxb* (**B**), *Col23a1* and*Lypd1* (**C**), *Chad, Chil1, Nell2*, and *Tnmd* (**D**) from Trabecular meshwork, uveal meshwork, and fibroblast clusters. Clusters are color coded as in Figure 4.

**Figure 5-figure supplement 1.**
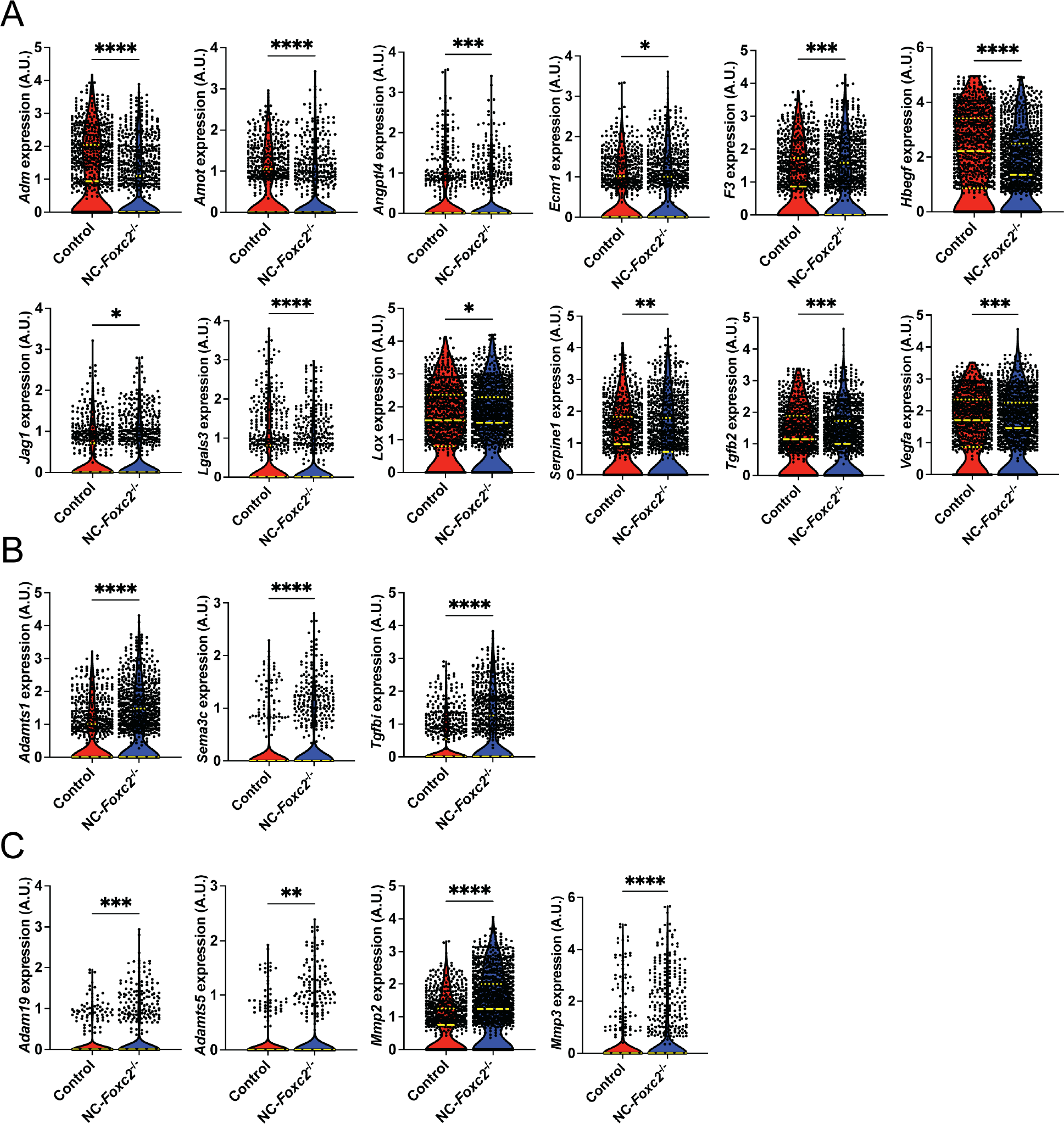
sc-RNA seq analysis identifies differentially expressed genes related to angiogenesis and ECM organization between Control and NC-*Foxc2^-/-^* mice in cells comprising the Trabecular meshwork 3 cluster. (A –**C**) Violin plots showing differential expression of *Adm, Amot, Angptl4, Ecm1, F3, Hbegf, Jag1, Lgals3, Lox, Serpine1, Tgfb2,* and *Vegfa* (**A**), *Adamts1, Sema3c,* and *Tgfbi* (**B**), *Adam19, Adamts5, Mmp2*, and *Mmp3* (**D**) between Control and NC-*Foxc2^-/-^* mice. Long, dashed lines denote median values. Statistical analysis: Mann-Whitney Test. * P < .05, ** P < .01, *** P < .001, **** P < .0001.

**Figure 7-figure supplement 1.**
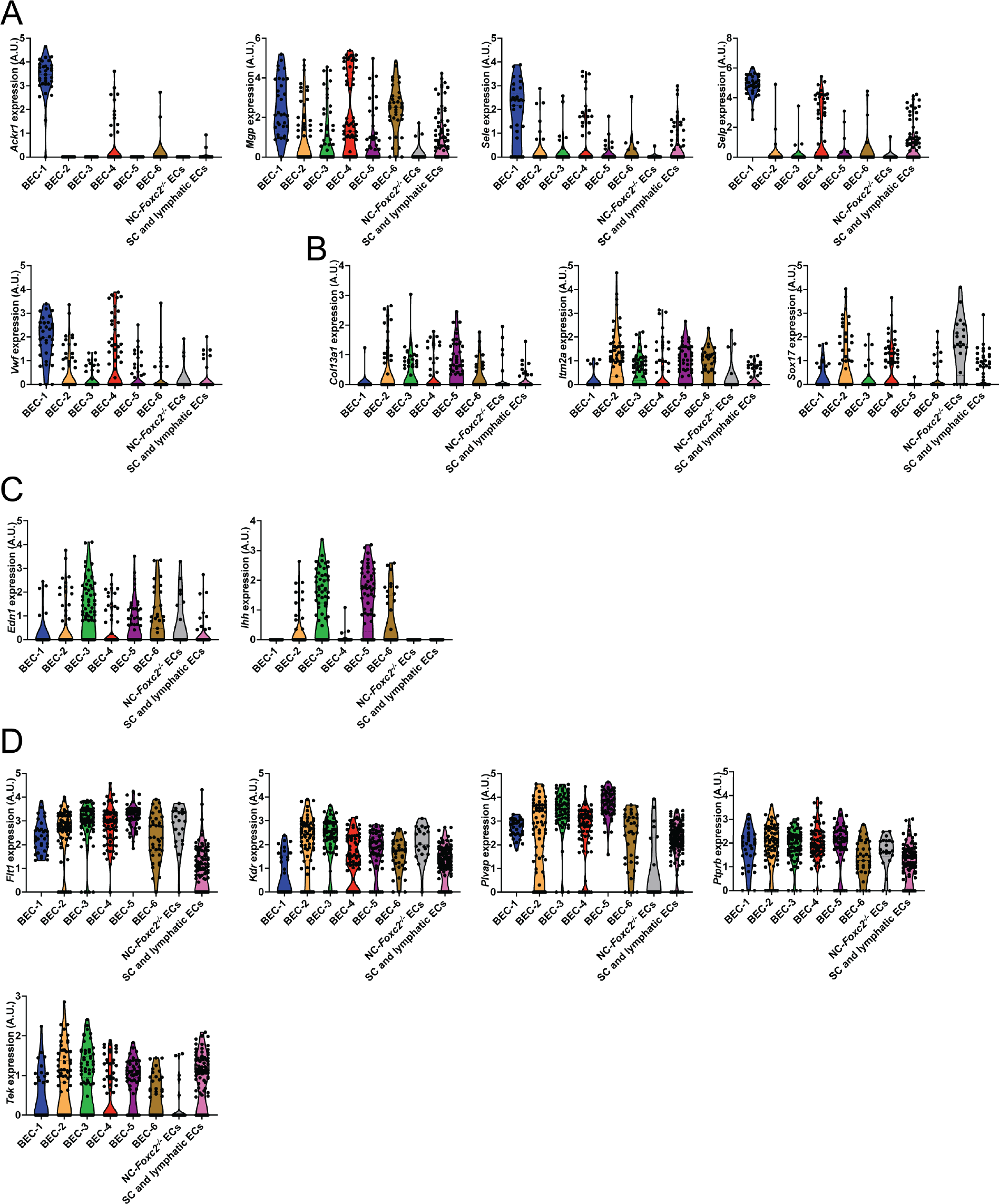
Characterization of endothelial subclusters. (**A – D**) Violin plots showing the expression of *Ackr1, Mgp, Sele*, *Selp,* and *Vwf* (**A**), *Col13a1, Itm2a,* and *Sox17* (**B**), *Edn1* and *Ihh* (**C**), *Flt1, Kdr, Plvap, Ptprb,* and *Tek* (**D**) from endothelial subclusters. Clusters are color coded as in Figure 7.

**Figure 8-figure supplement 1.**
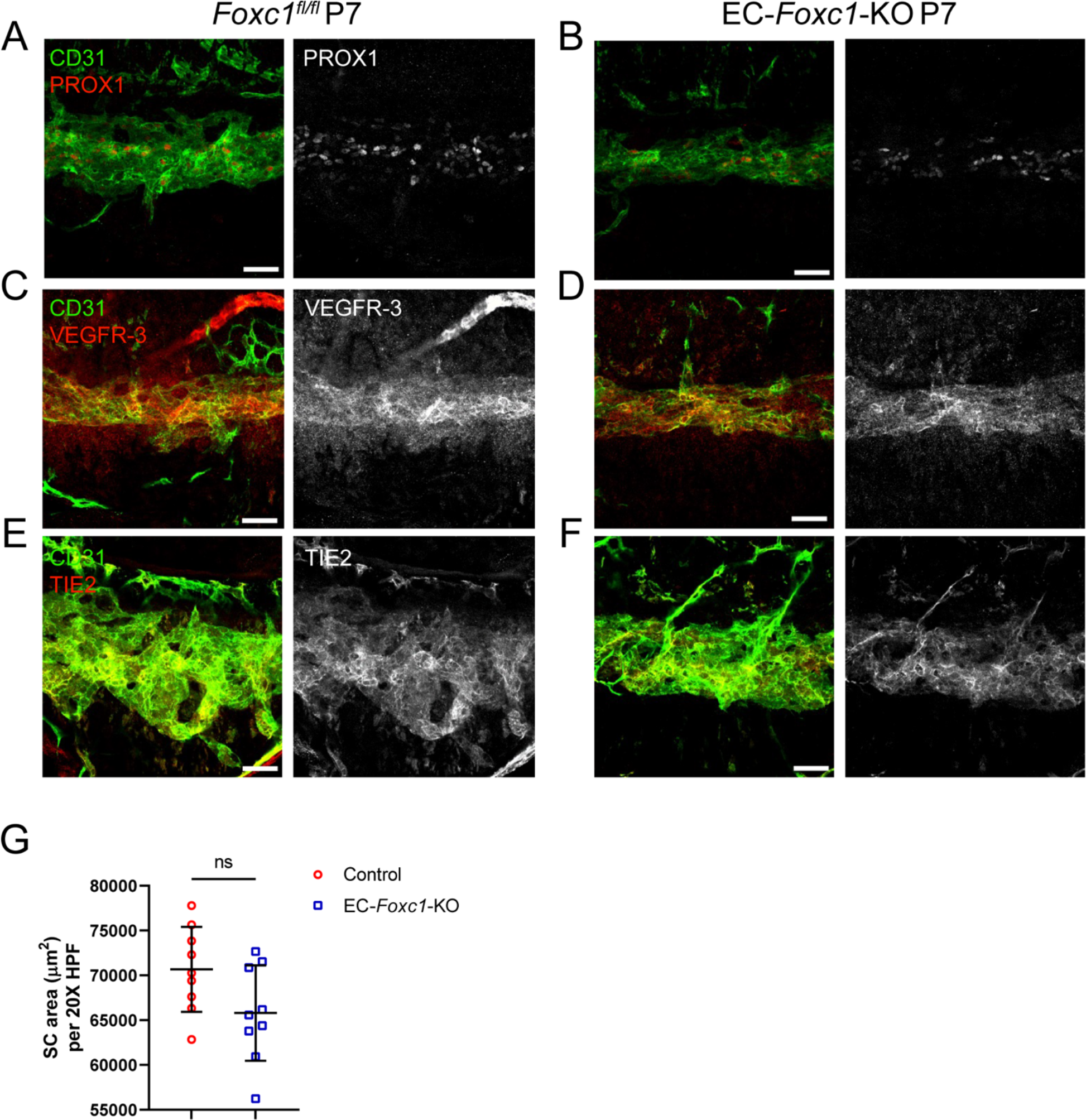
SC morphogenesis is not severely impacted by early postnatal, endothelial-specific deletion of *Foxc1*. (A–F) Representative images of CD31 and PROX1 (A, B), VEGFR-3 (C, D), or Tie2 (E, F) expression in the SC of P7 *Foxc1^fl/fl^* control (A, C, E) and EC-*Foxc1*-KO mice (B, D, F). Scale bars are 50 μm. (G) Quantification of SC area per 20X high-power field in P7 *Foxc1^fl/fl^* control and EC-*Foxc1*-KO mice. N = 9 for Control and N= 9 for EC-*Foxc1*-KO mice. Data are mean ± SD. Statistical analysis: Student’s unpaired t-test.

**Figure 10-figure supplement 1.**
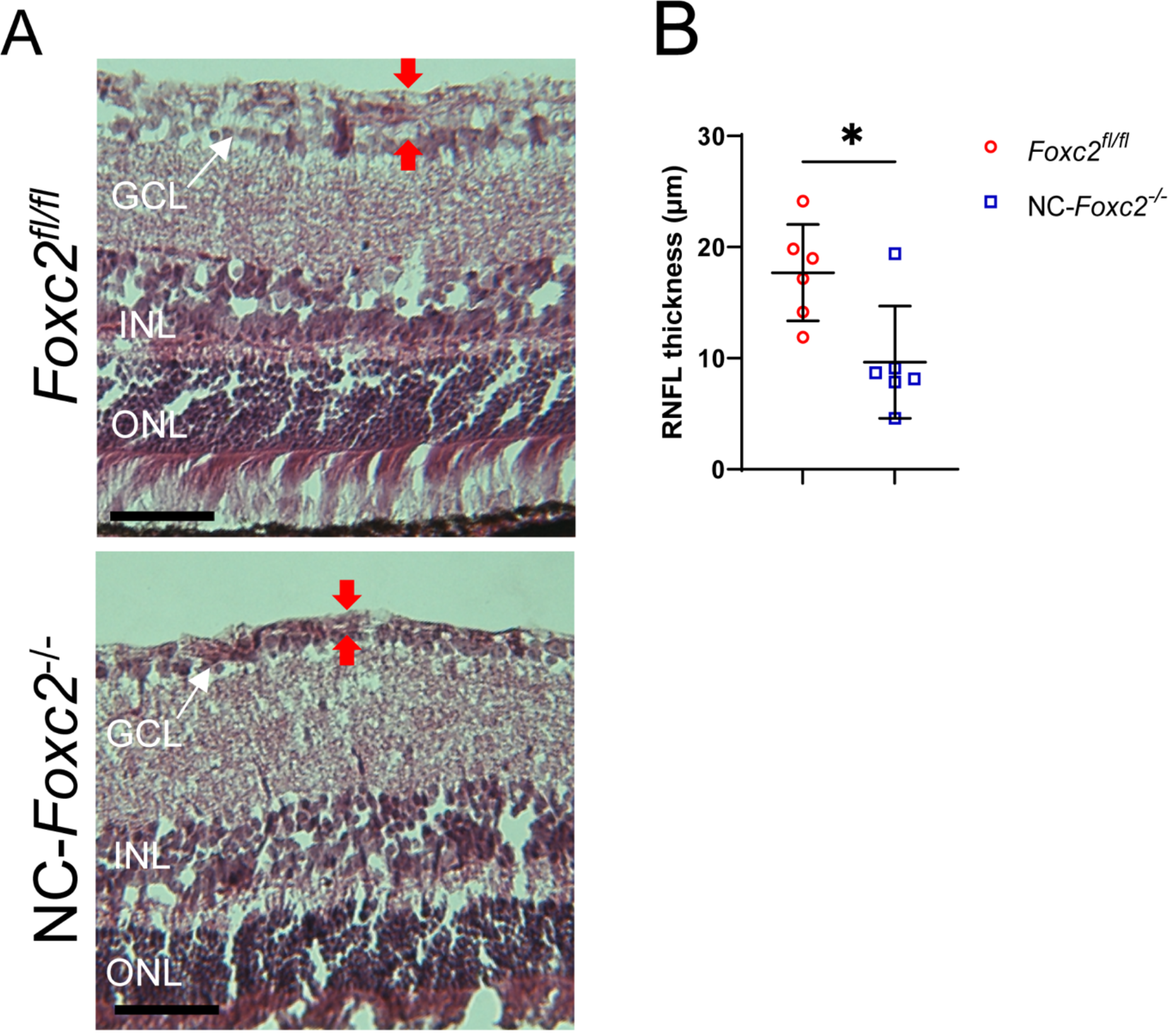
Characterization of retina nerve fiber layer thickness in NC-*Foxc2^-/-^* mice. (**A, B**) Representative images and quantification of retina nerve fiber layer (RNFL) thickness from hematoxylin and eosin-stained sections. N = 6 *Foxc2^fl/fl^* control and N = 6 for NC-*Foxc2^-/-^* mice. Data are mean ± SD. Statistical analysis: Student’s unpaired t-test. * P < .05. ONL = outer nuclear layer, INL = inner nuclear layer, GCL = ganglion cell layer.

**Supplementary Table 1.**
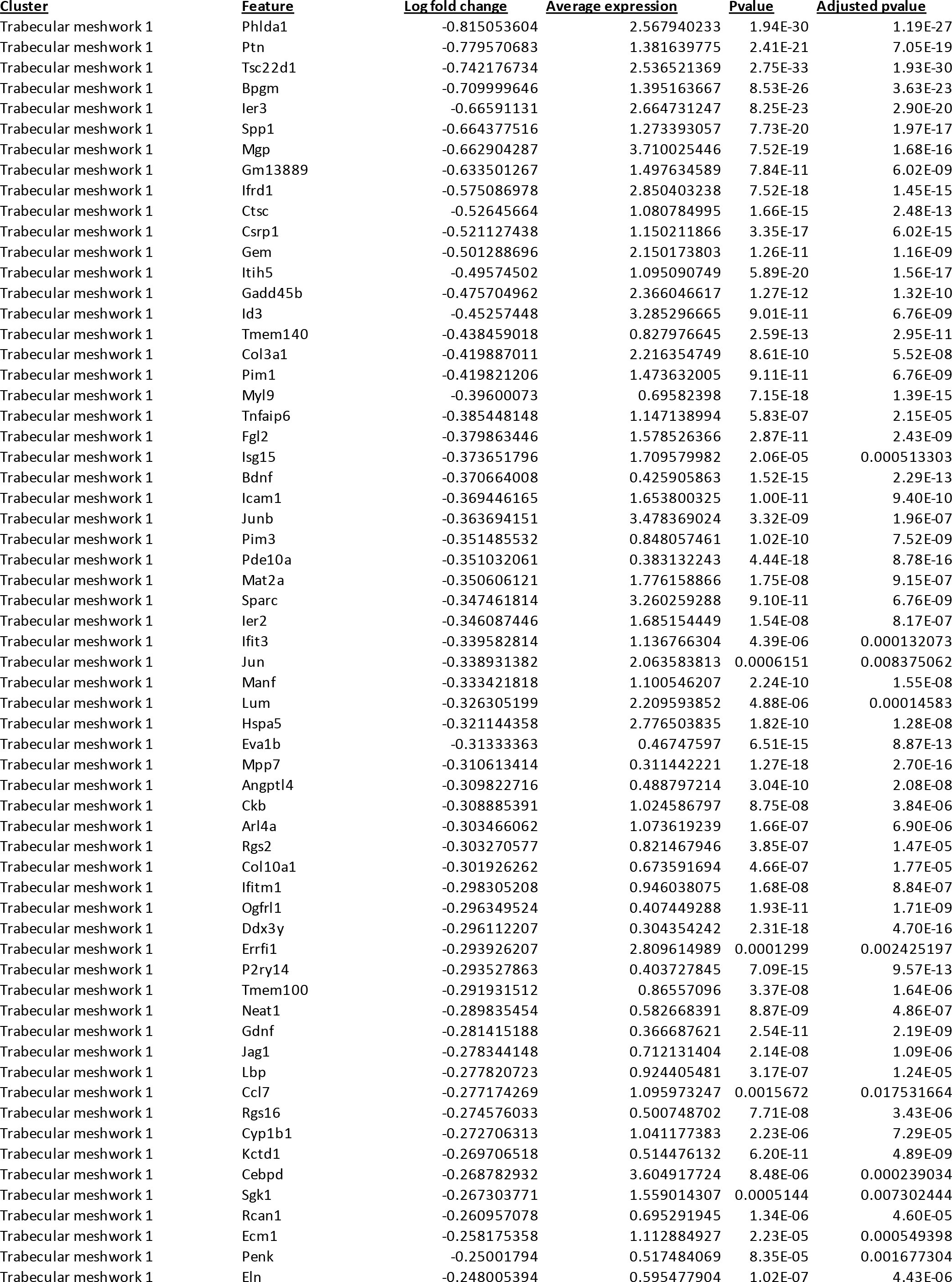
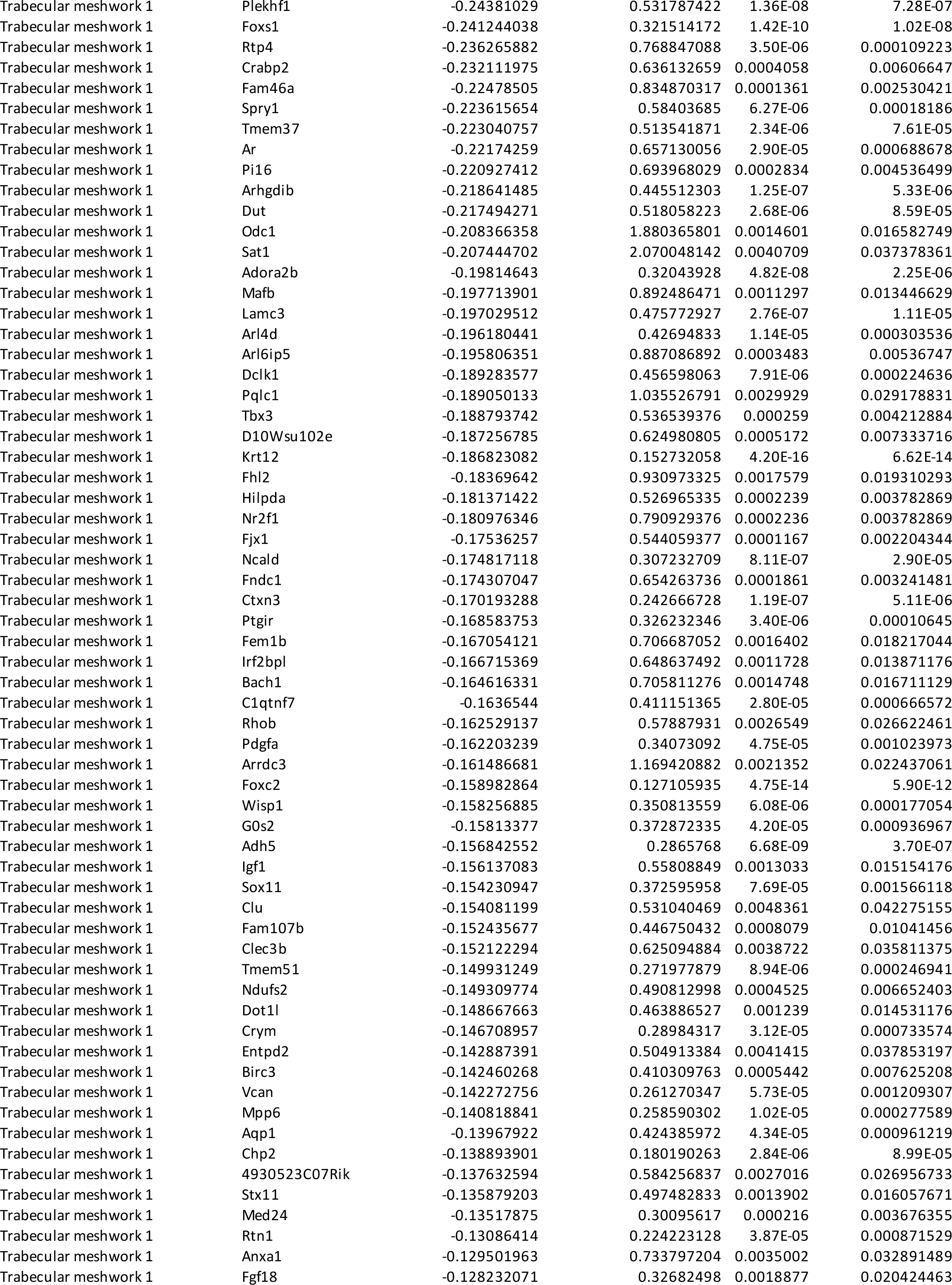
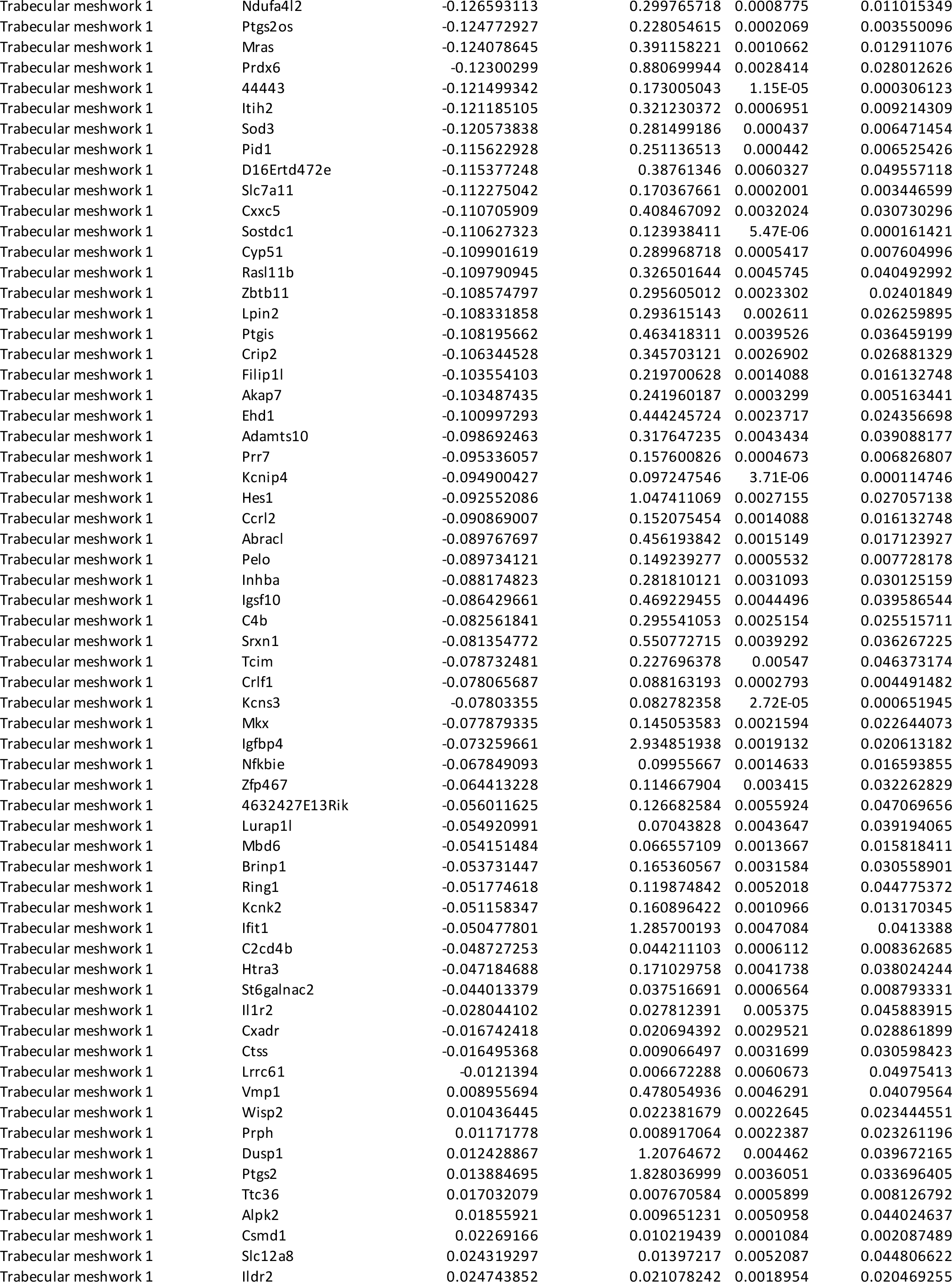
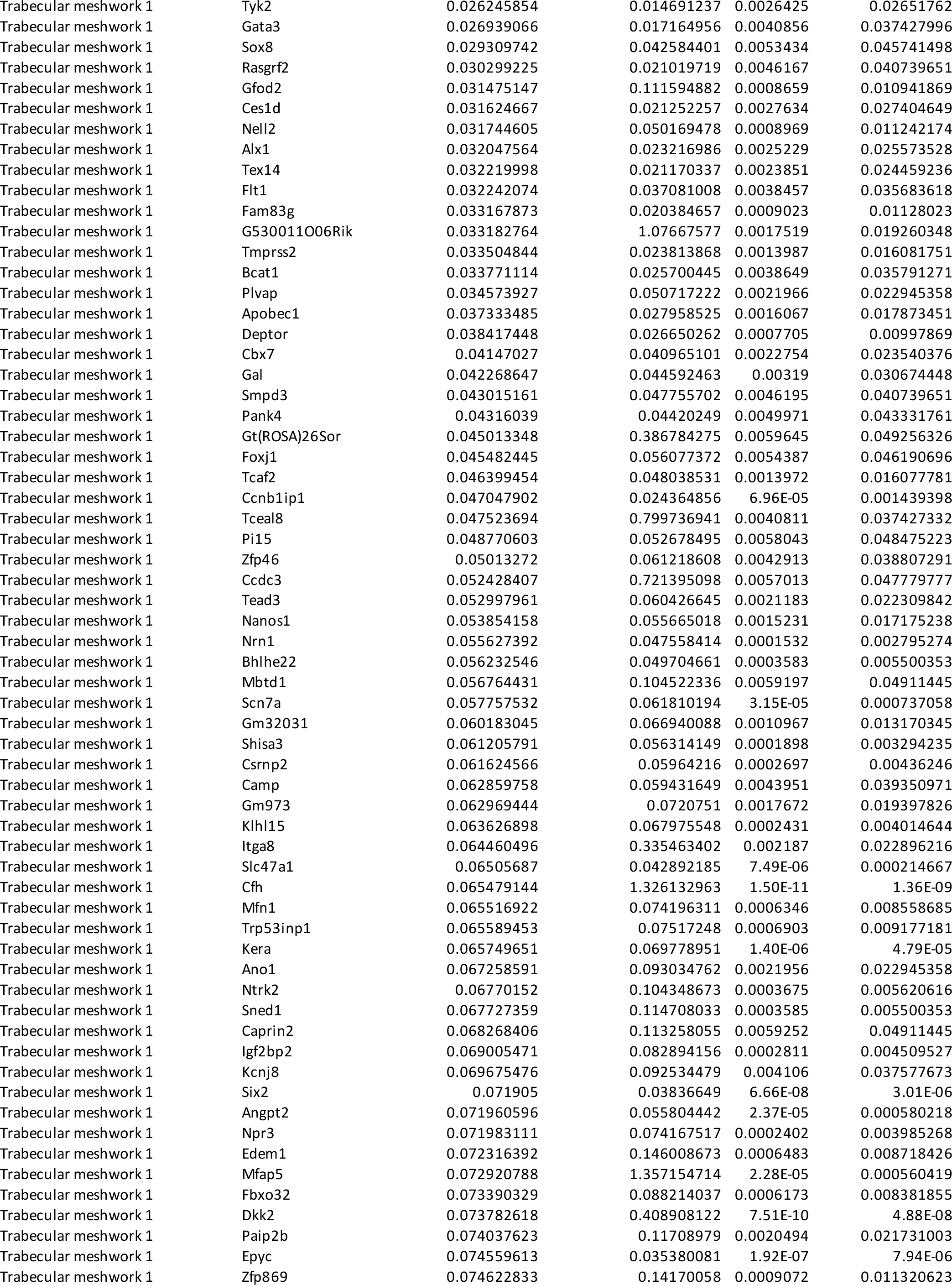
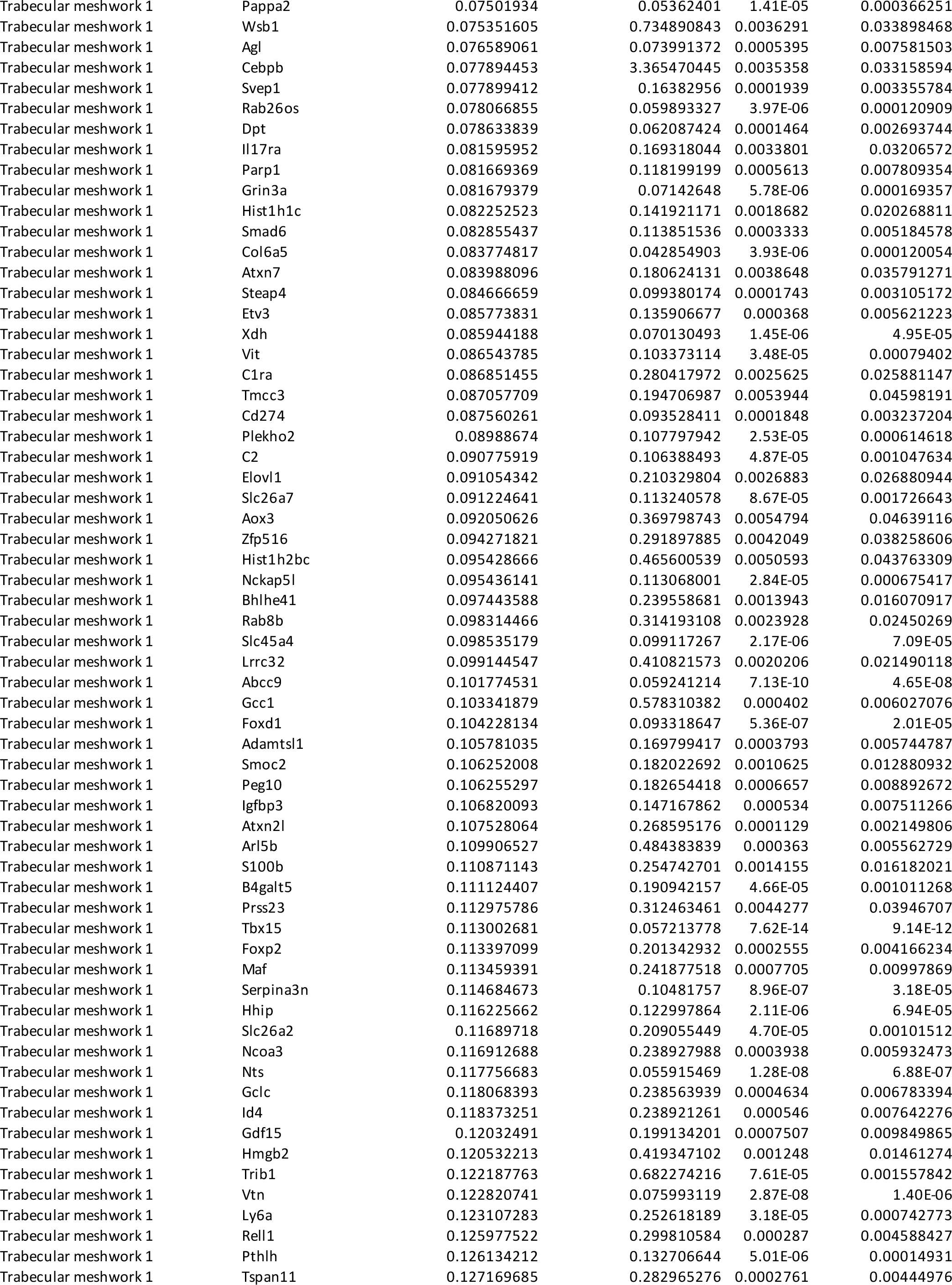
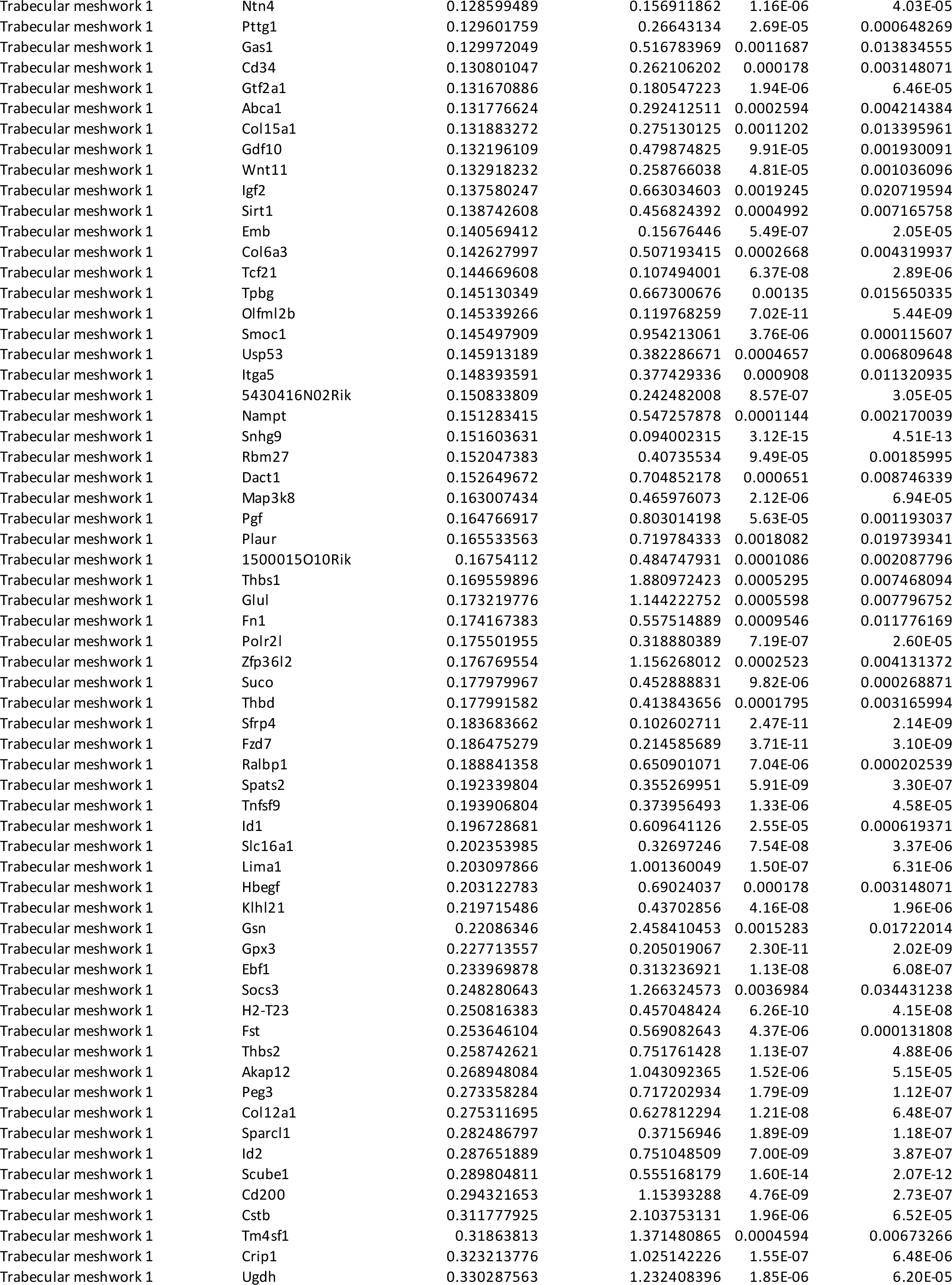
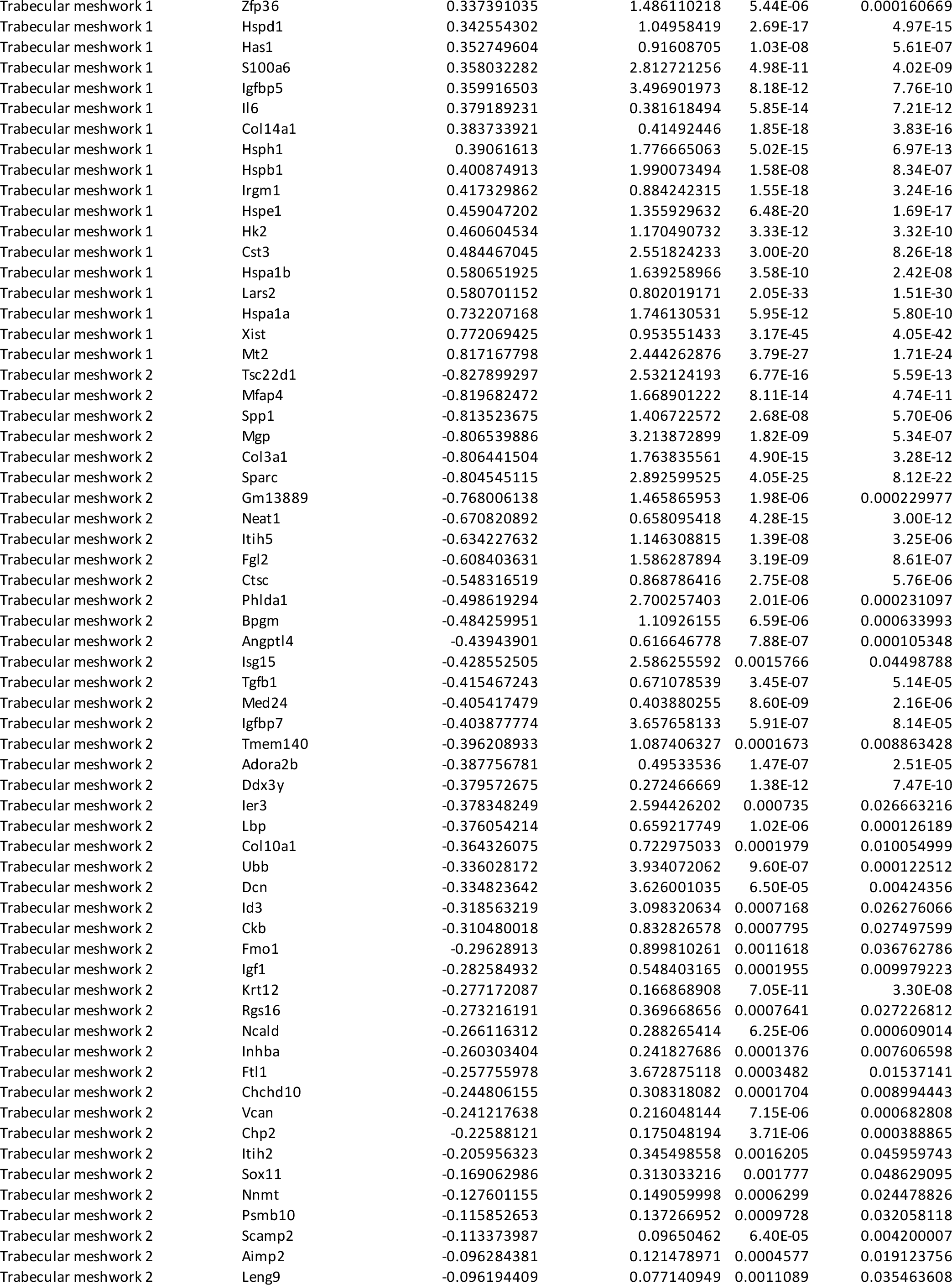
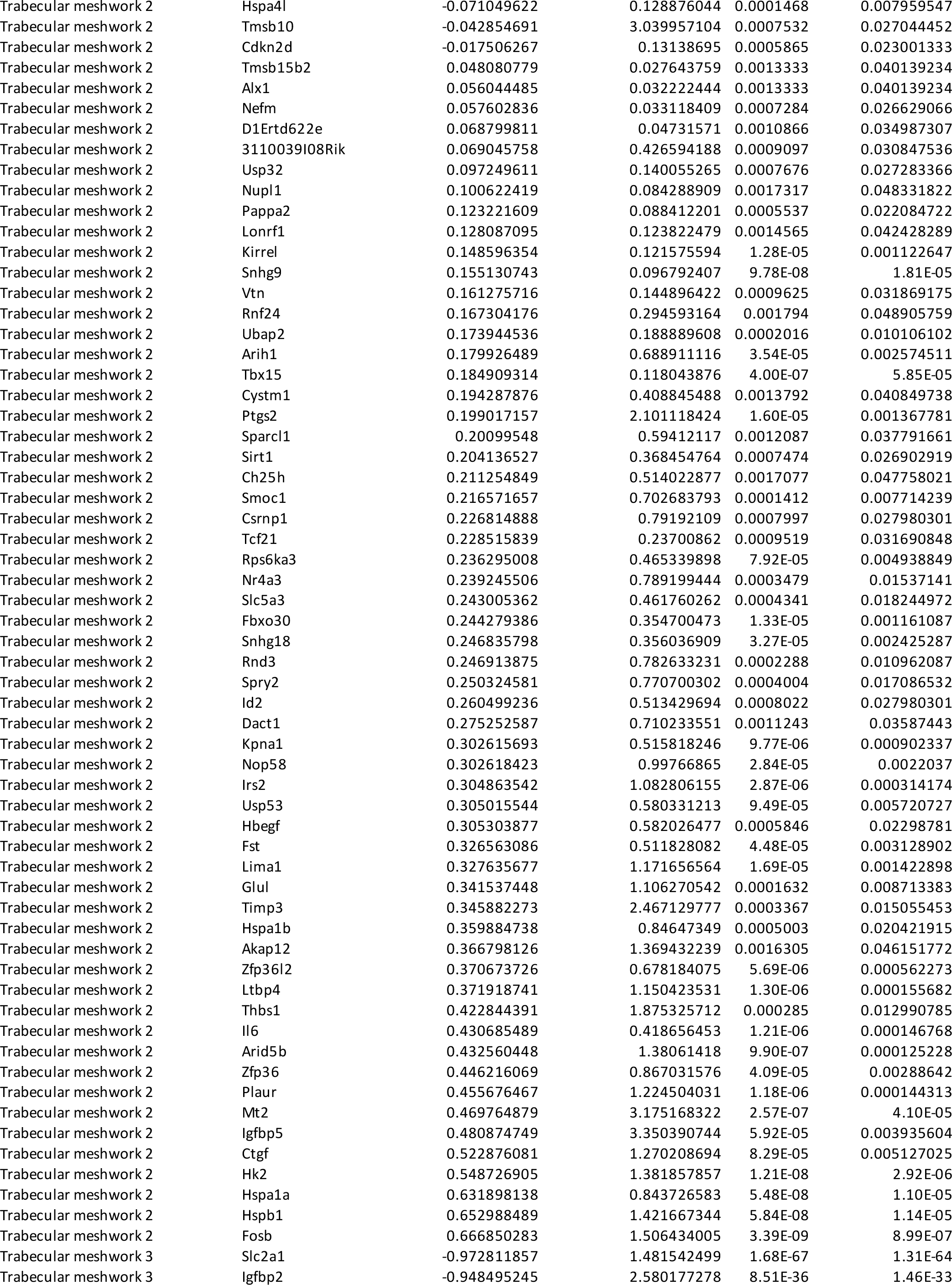
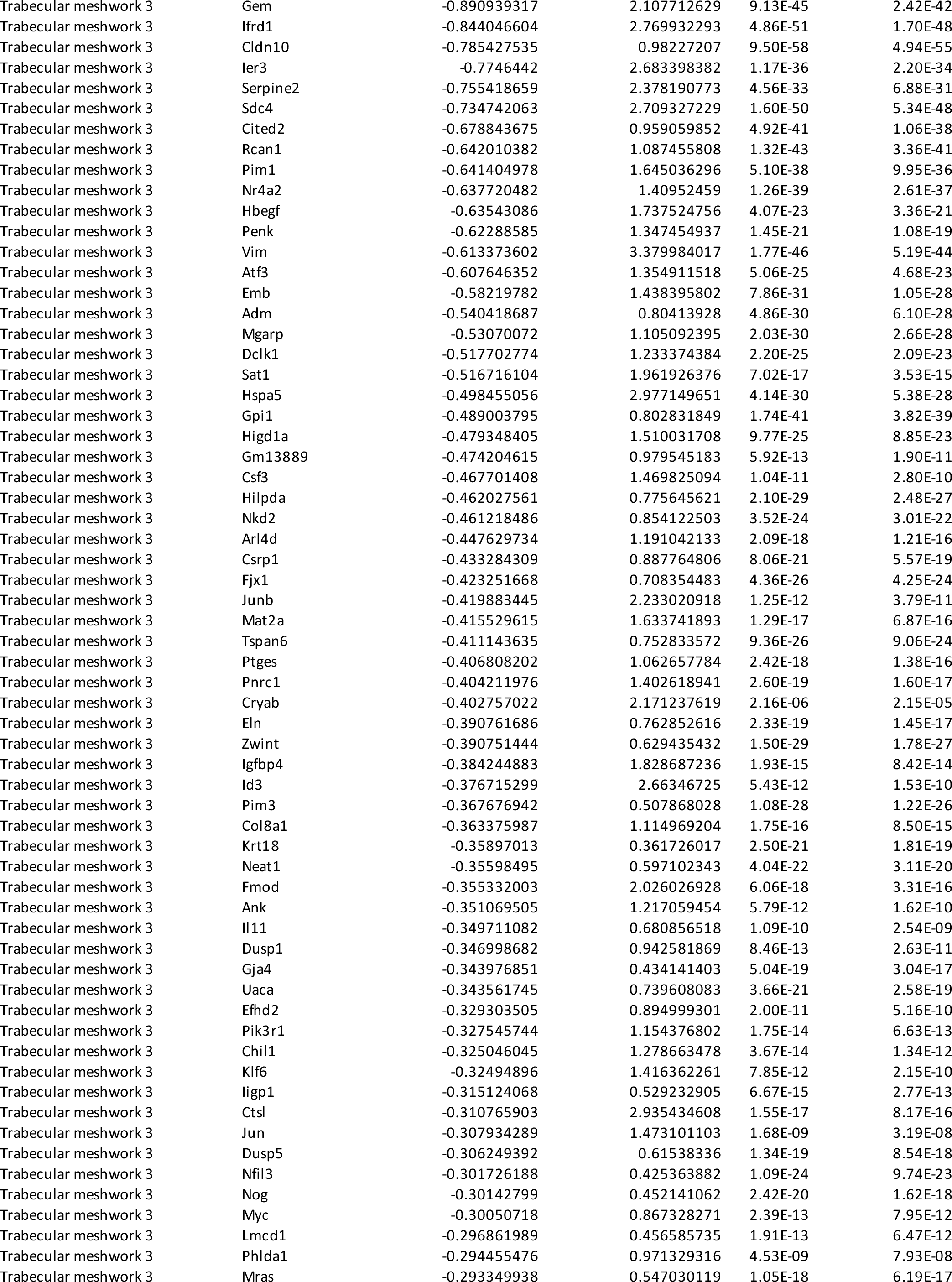
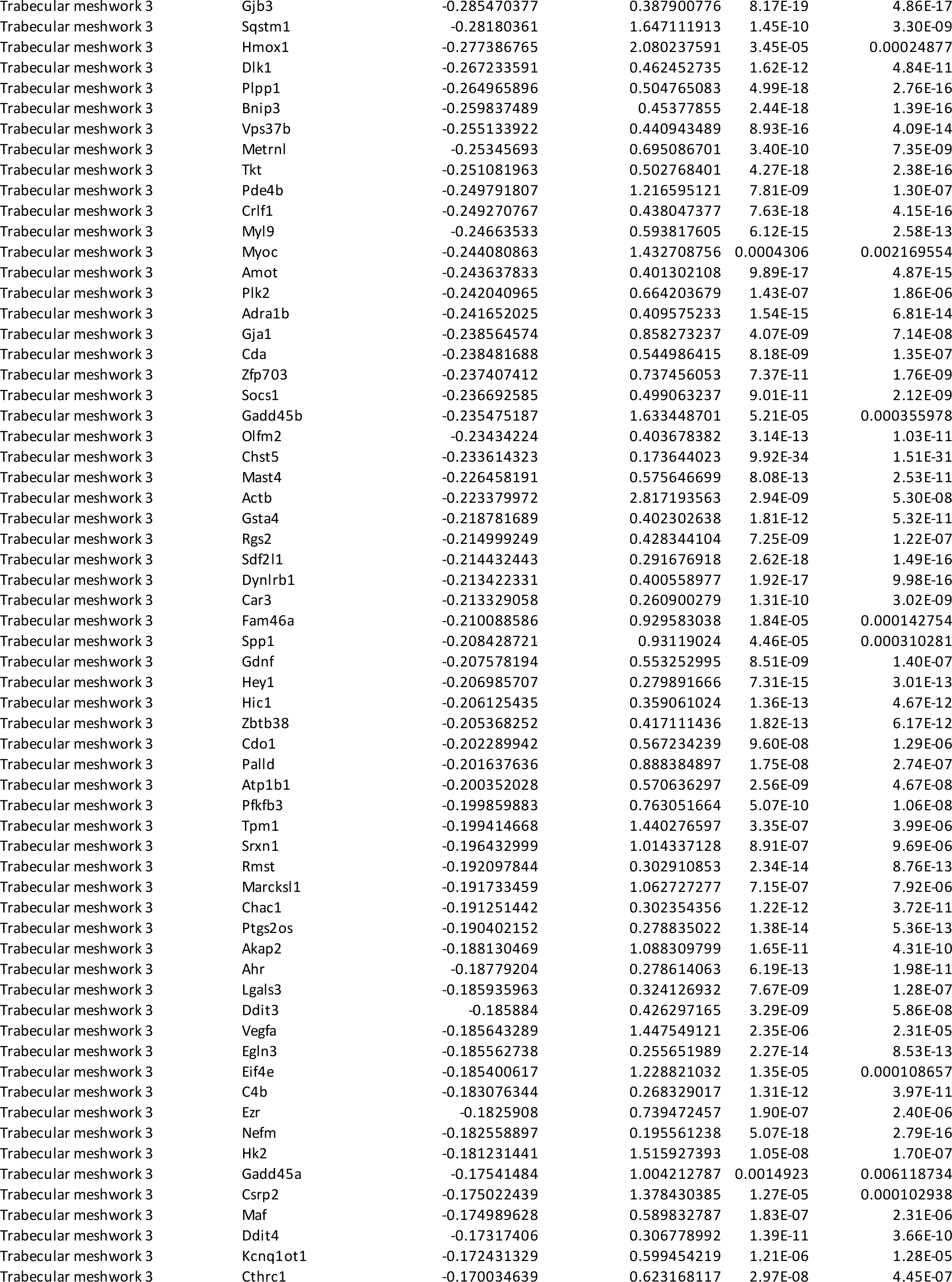
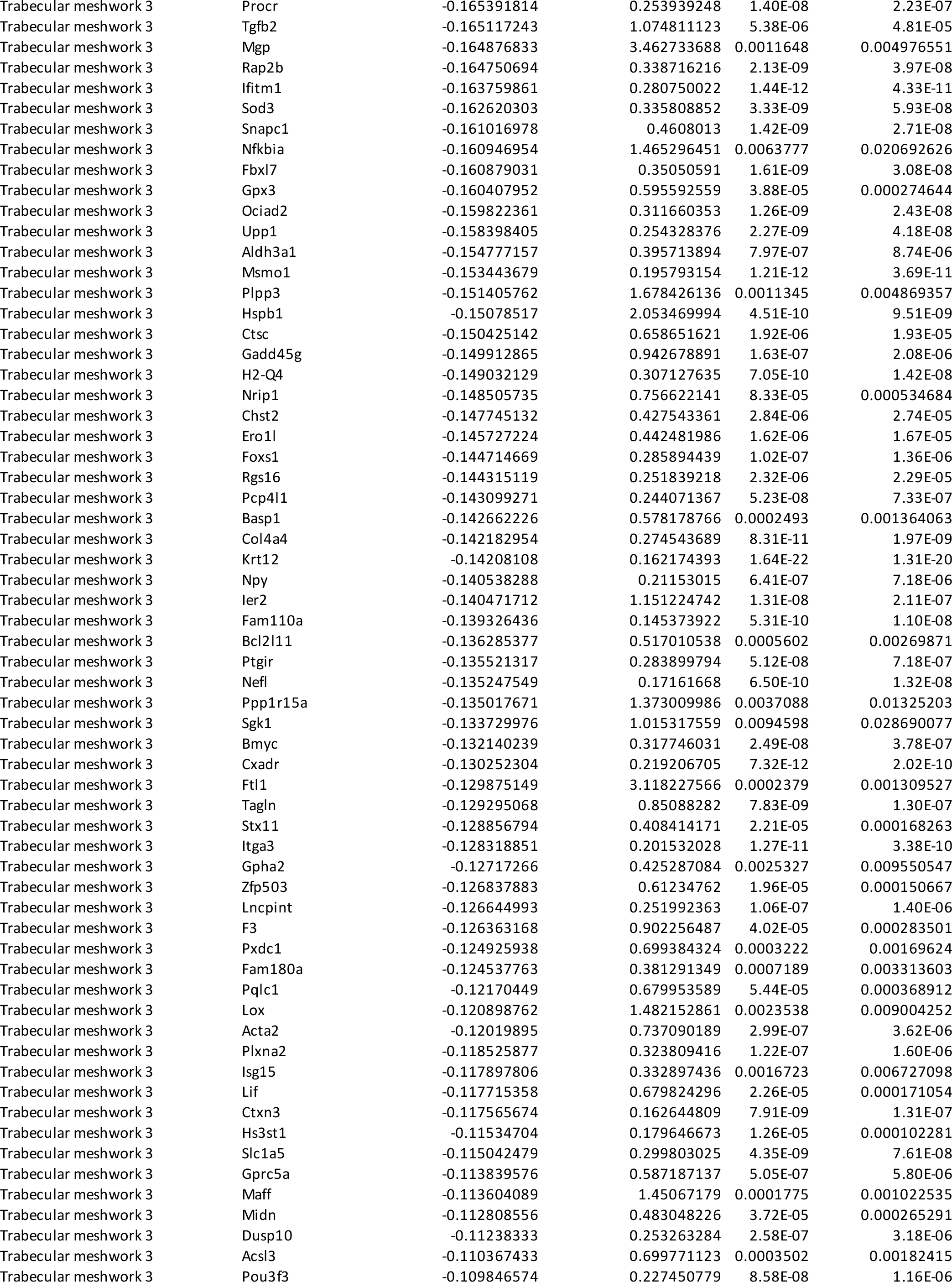
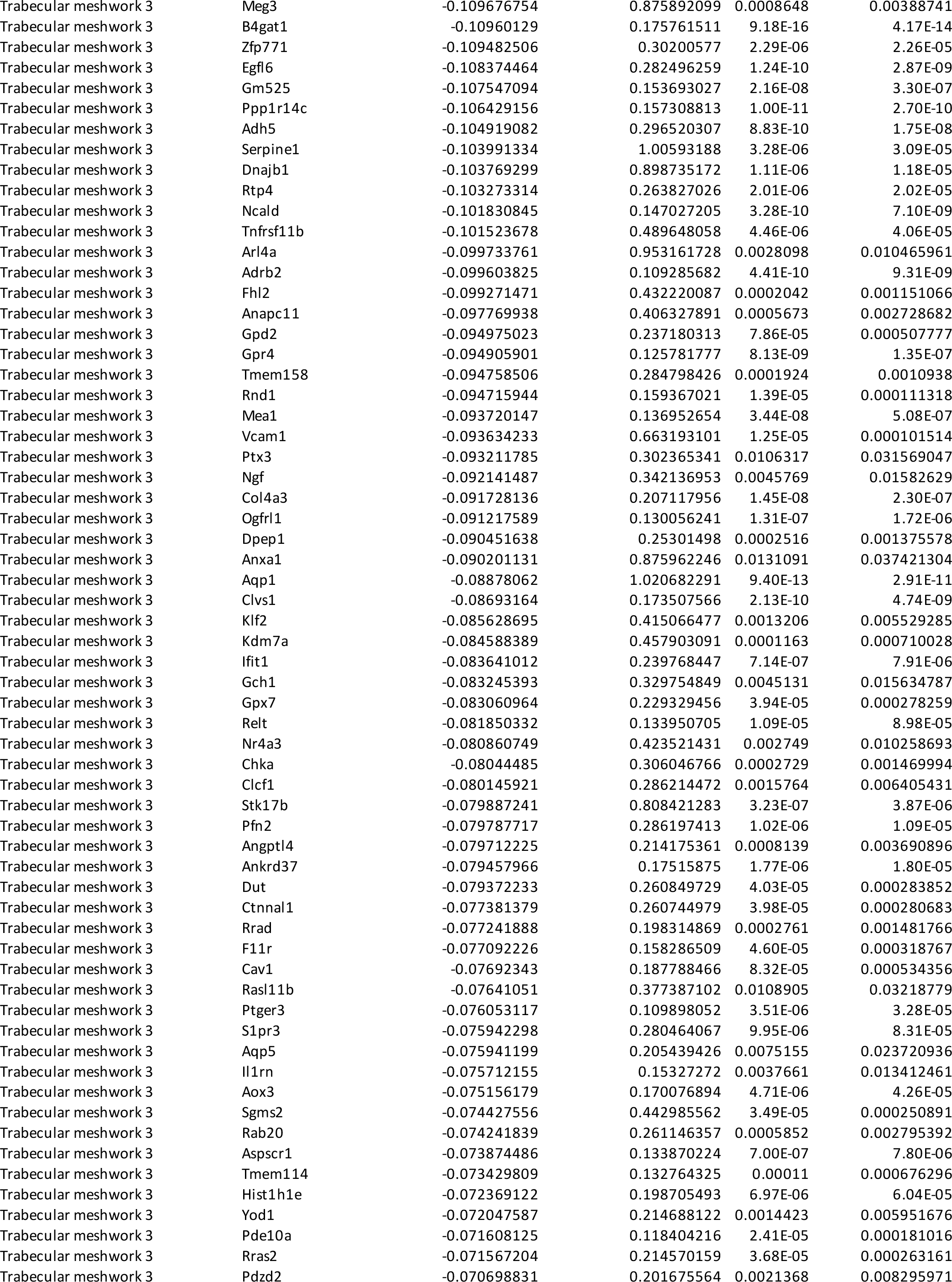
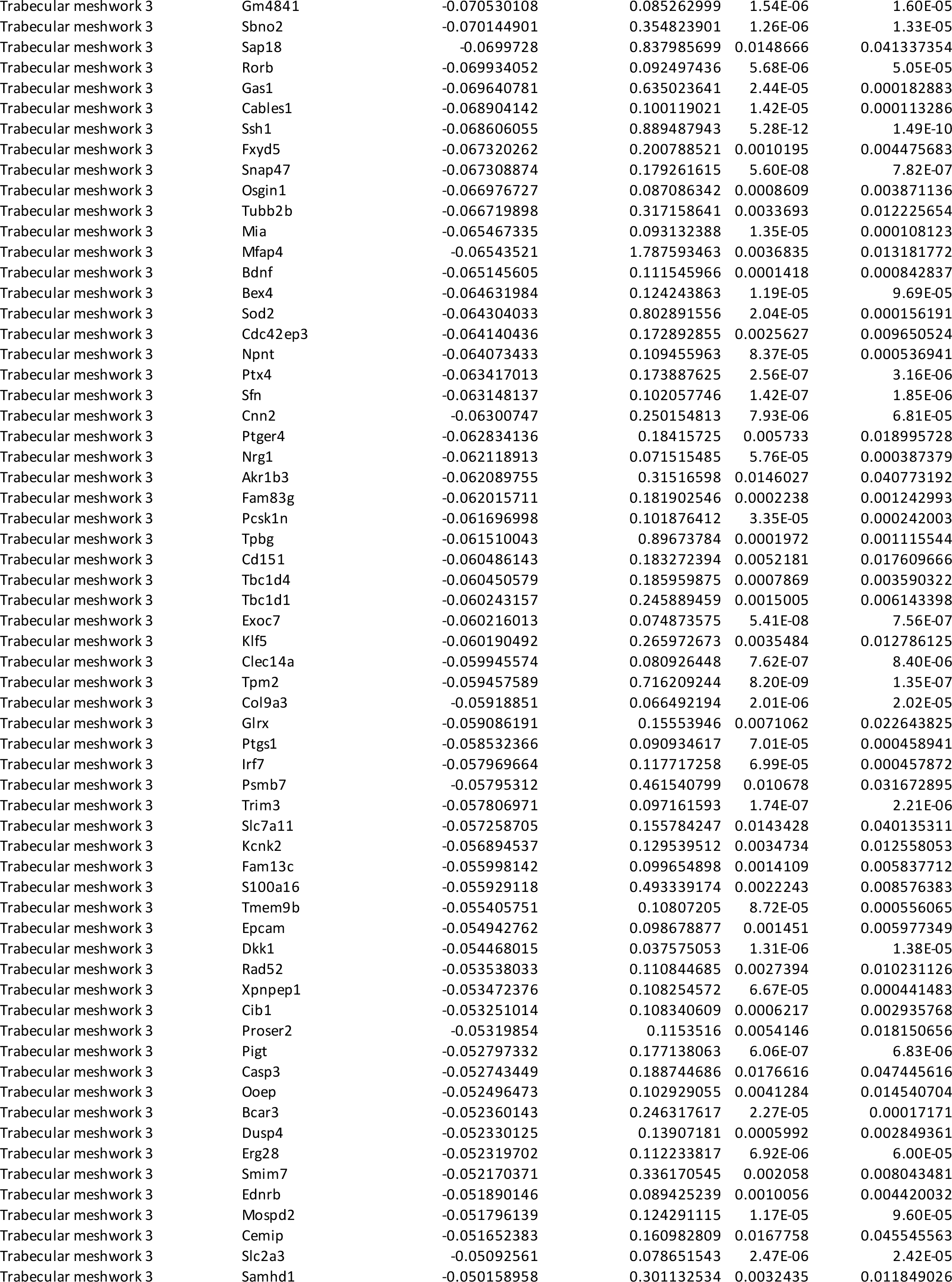
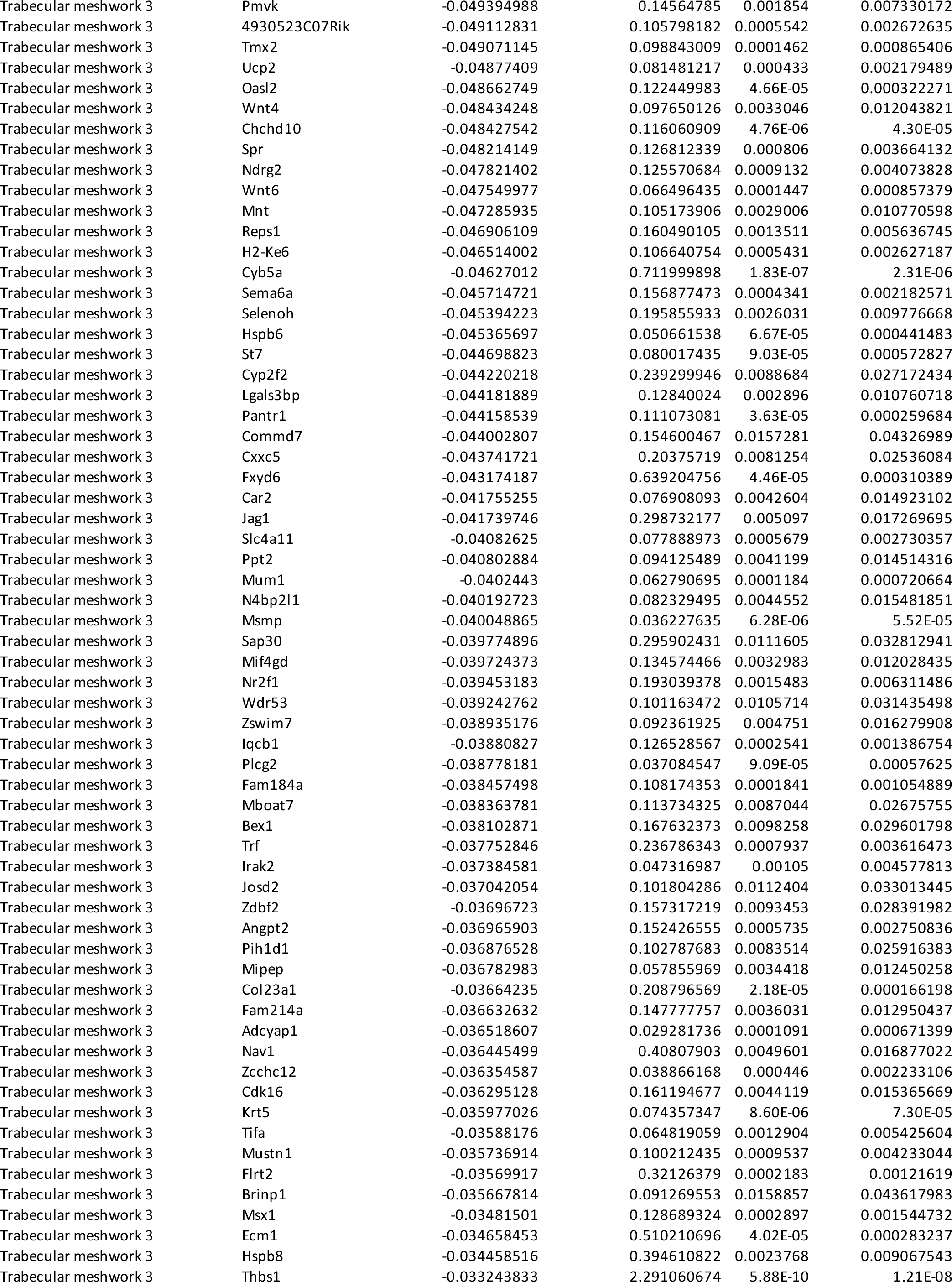
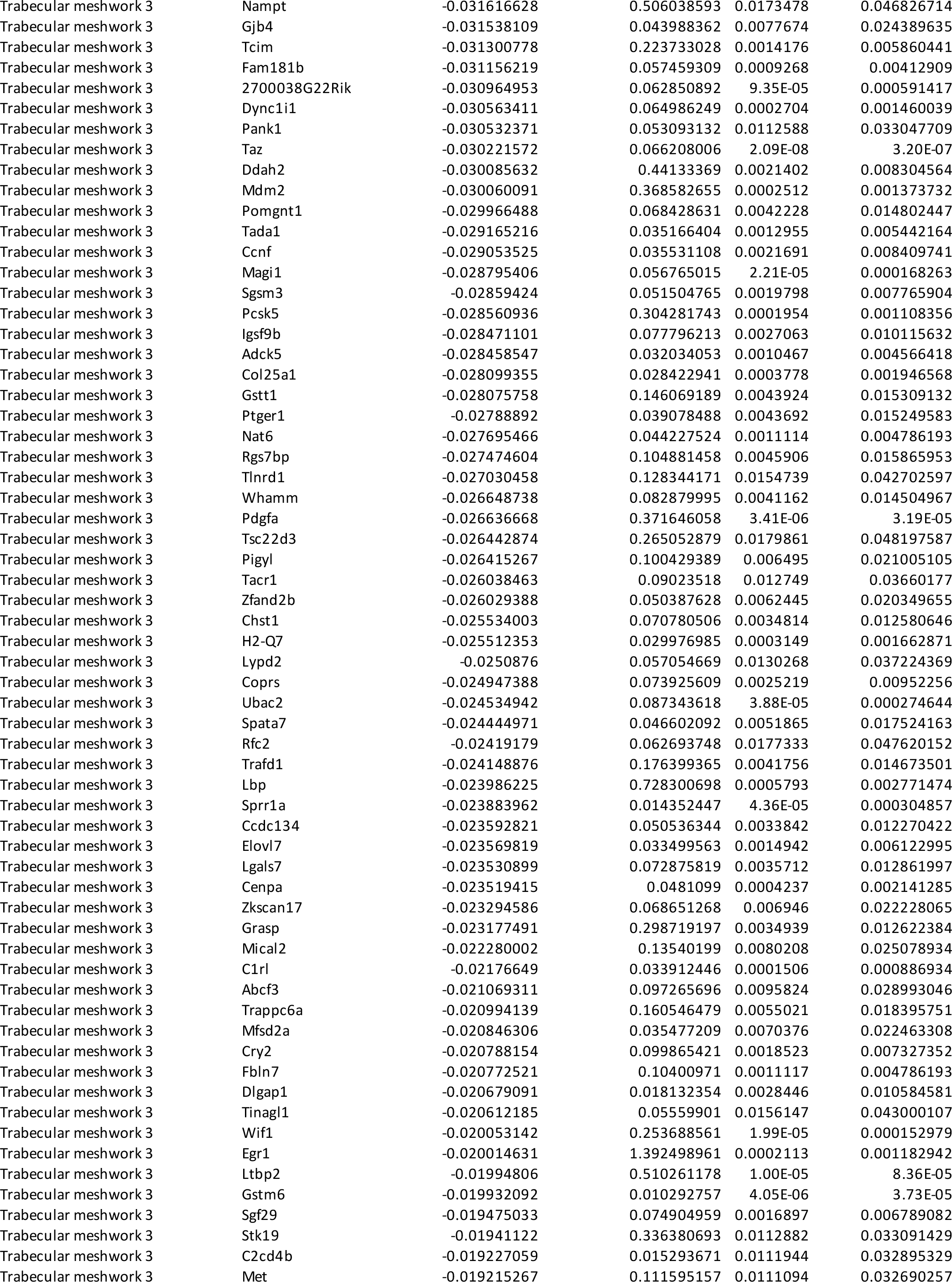
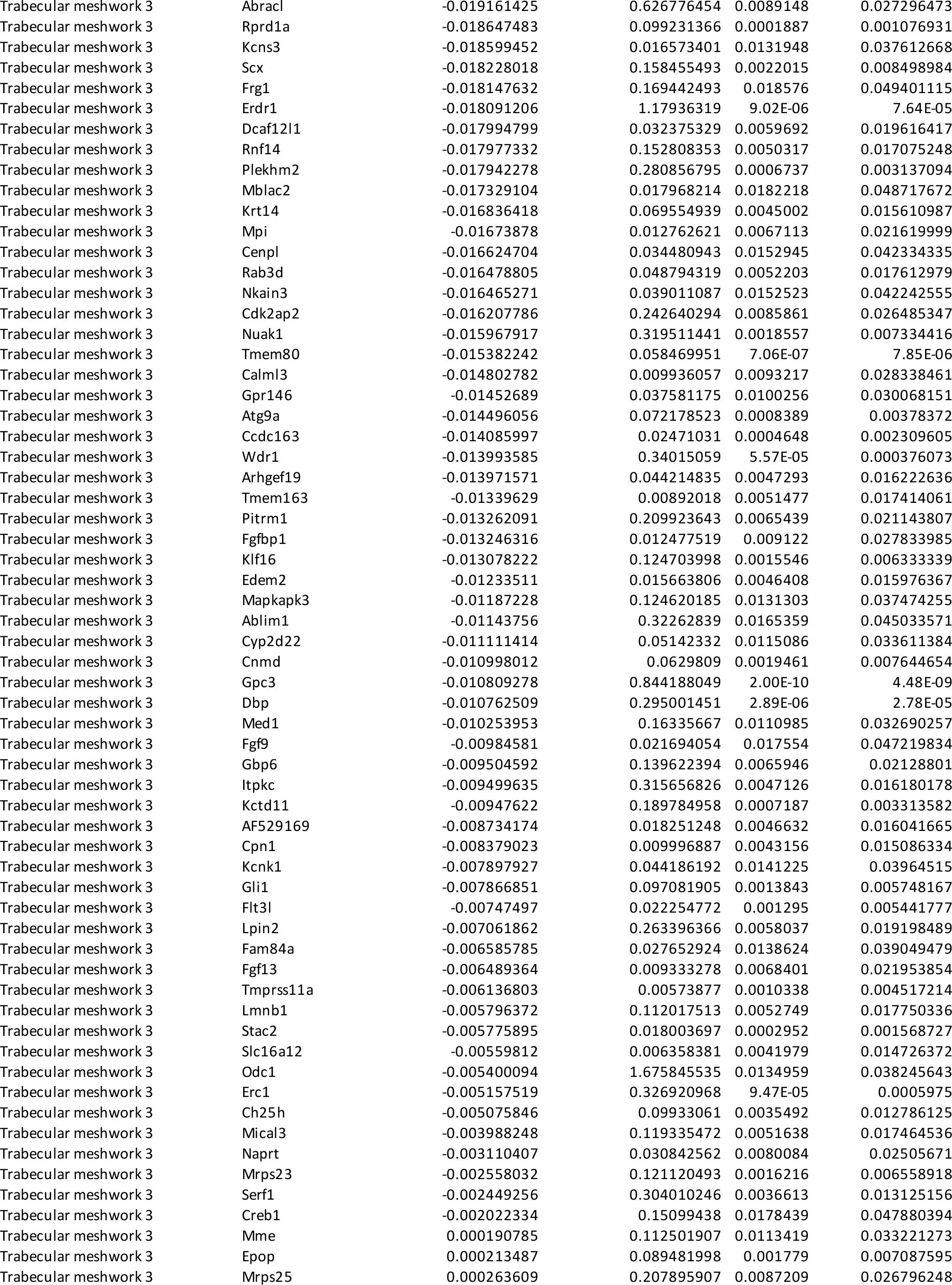
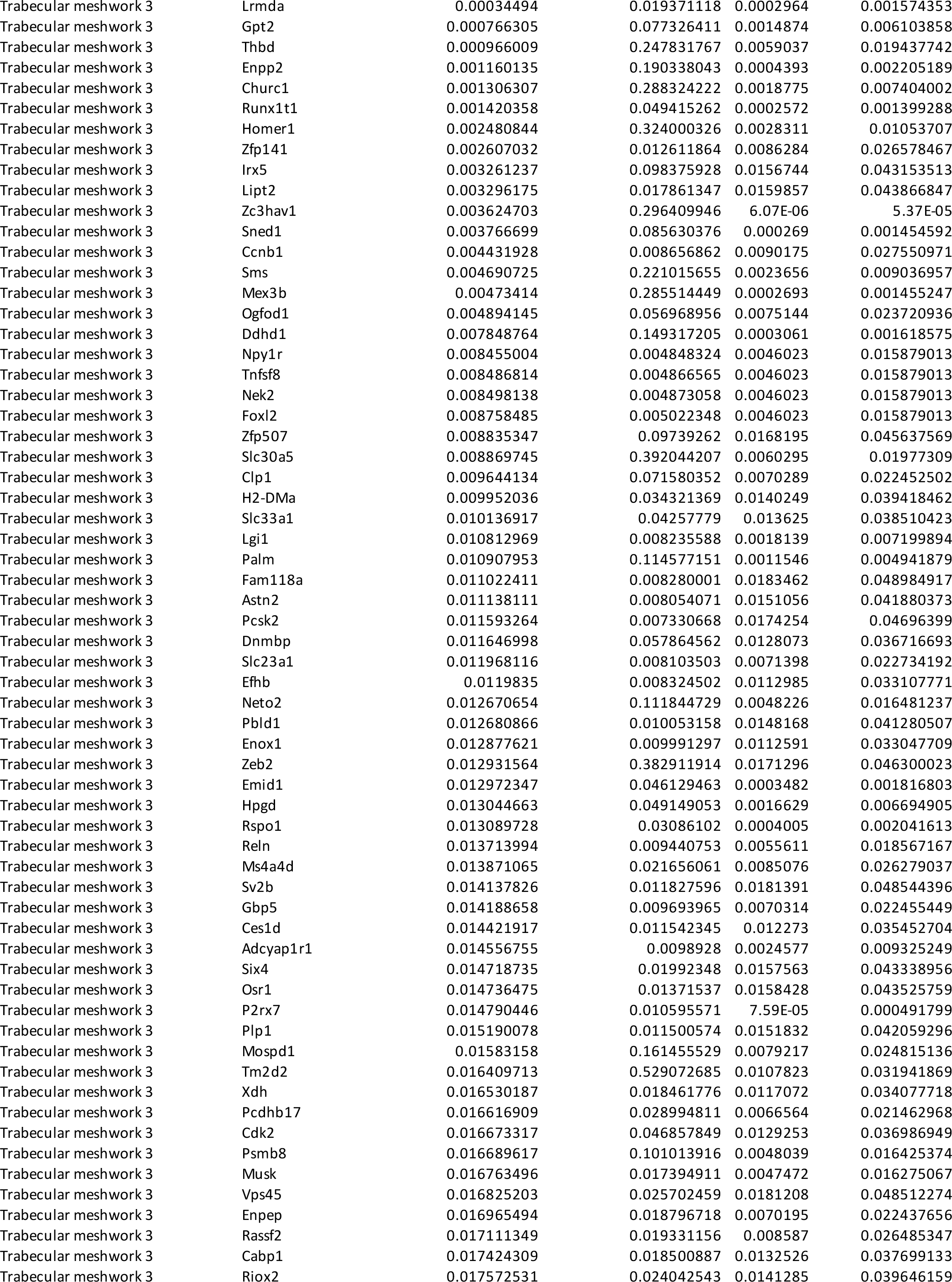
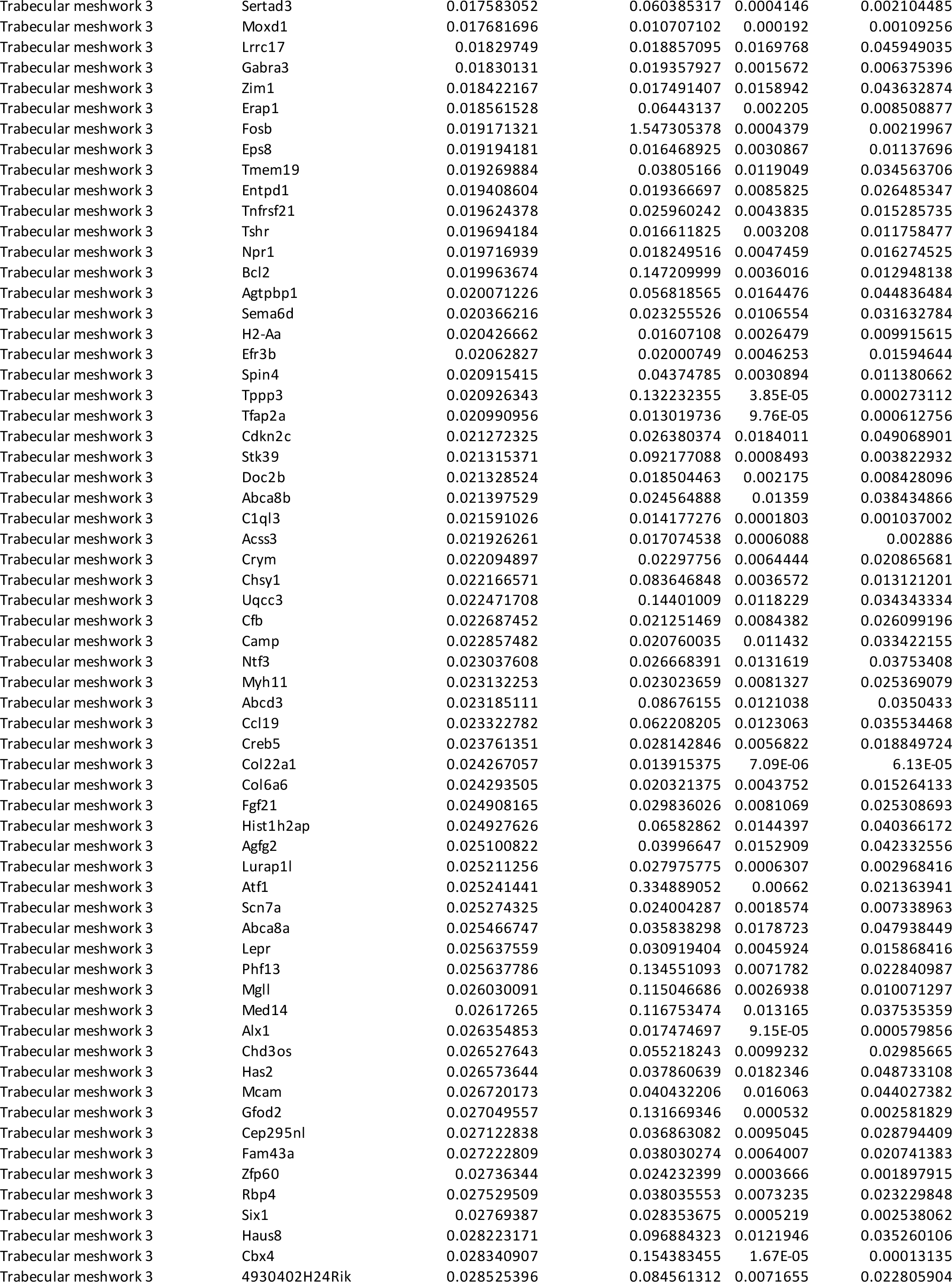
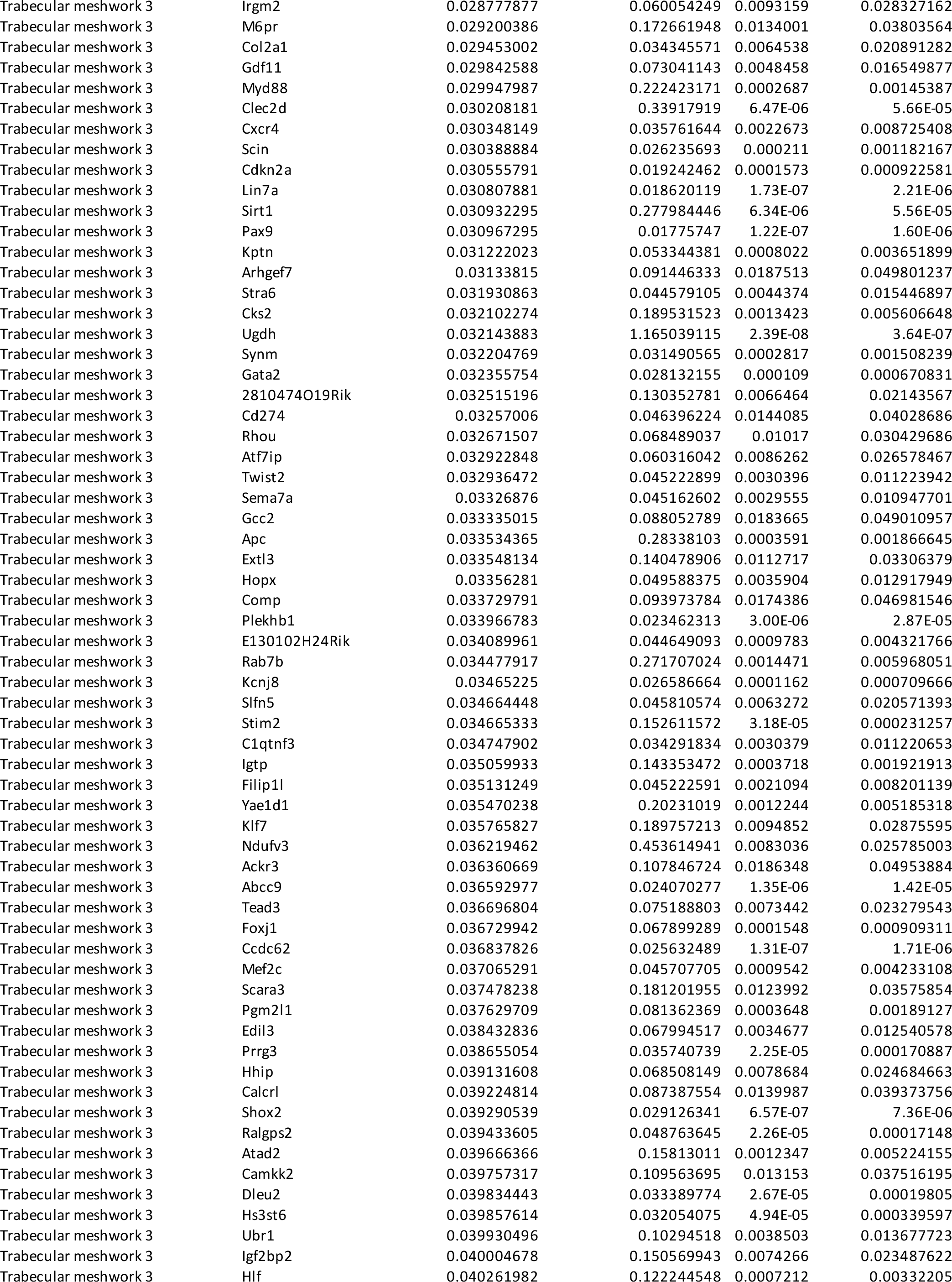
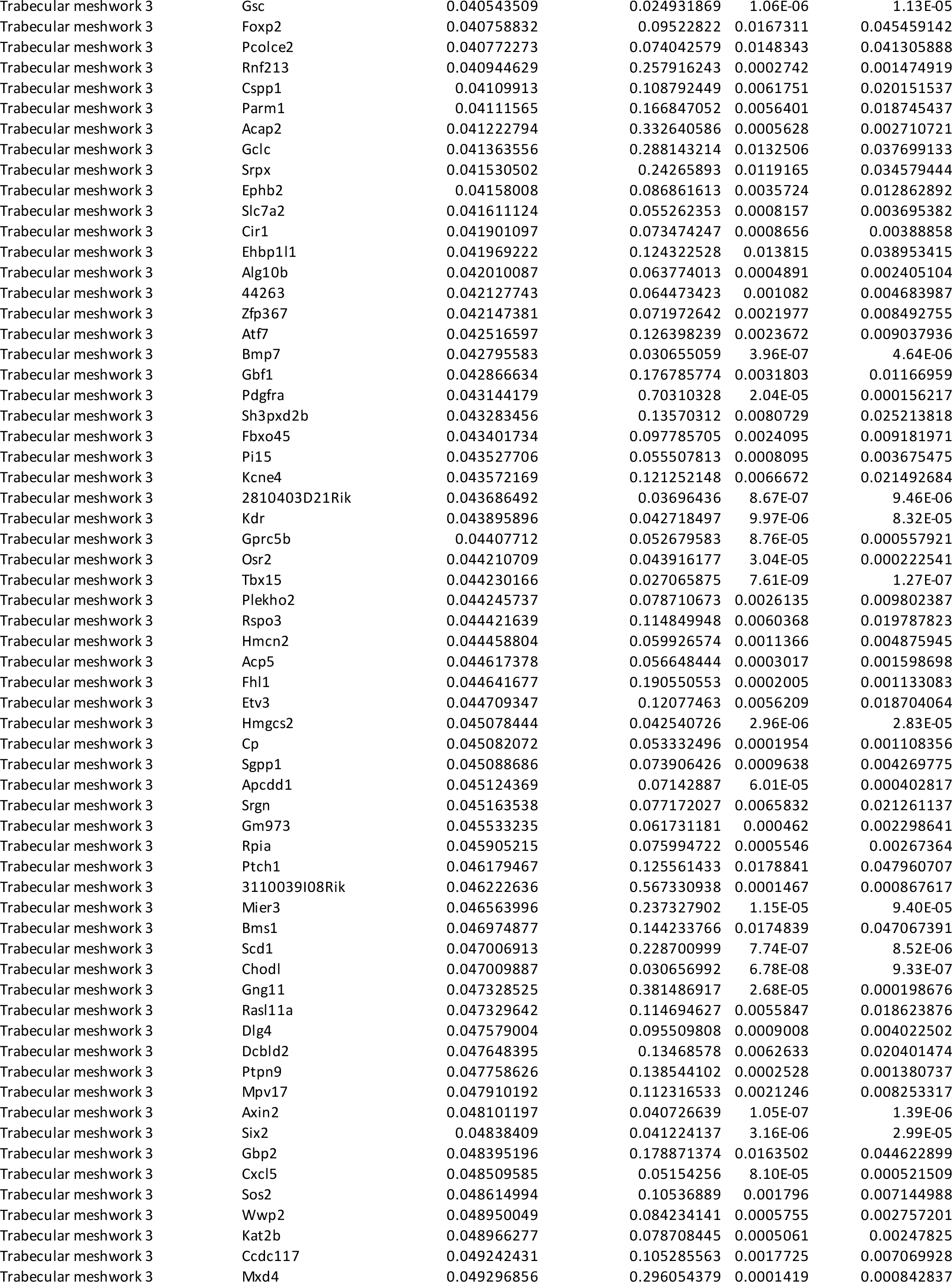
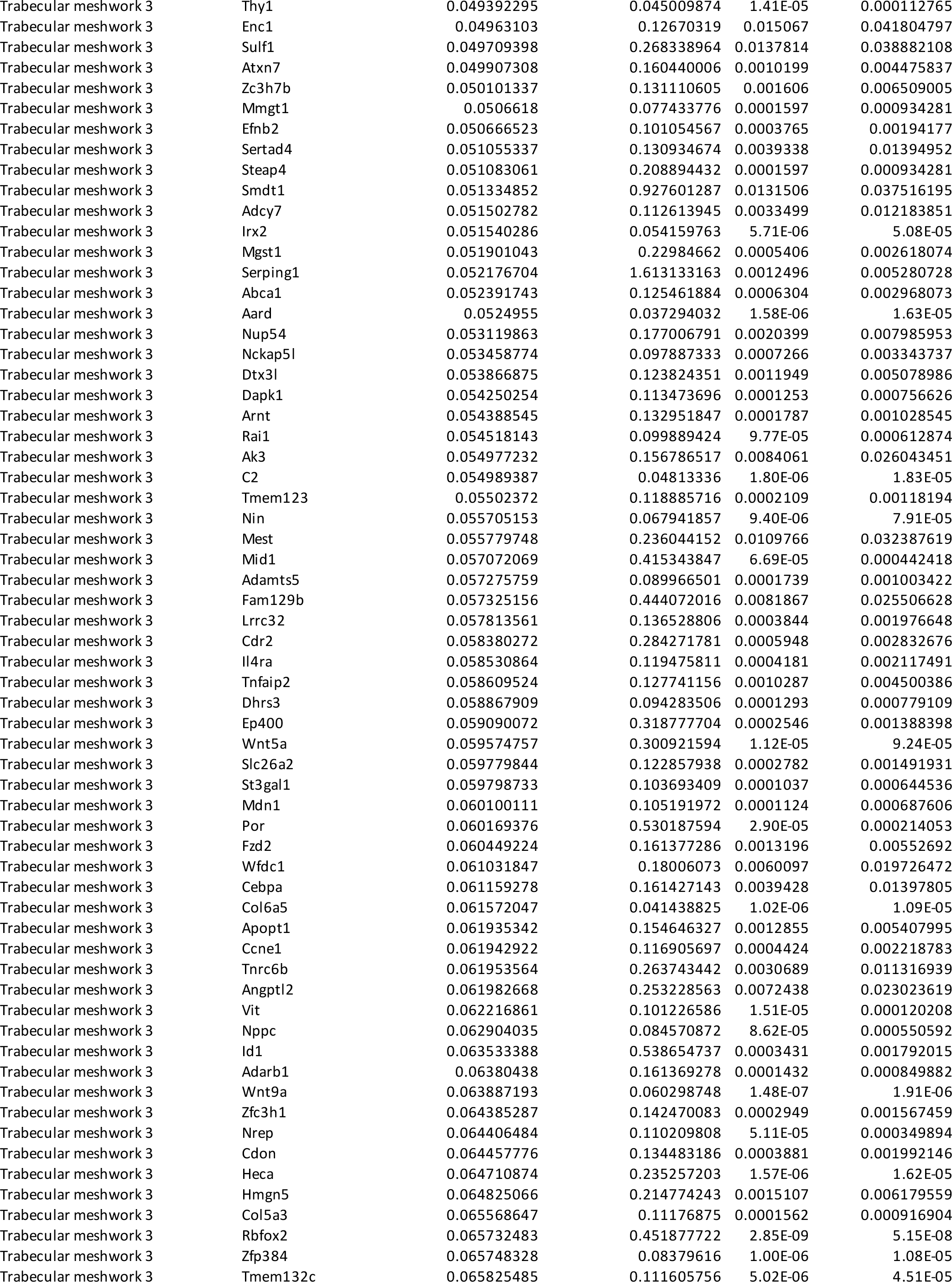
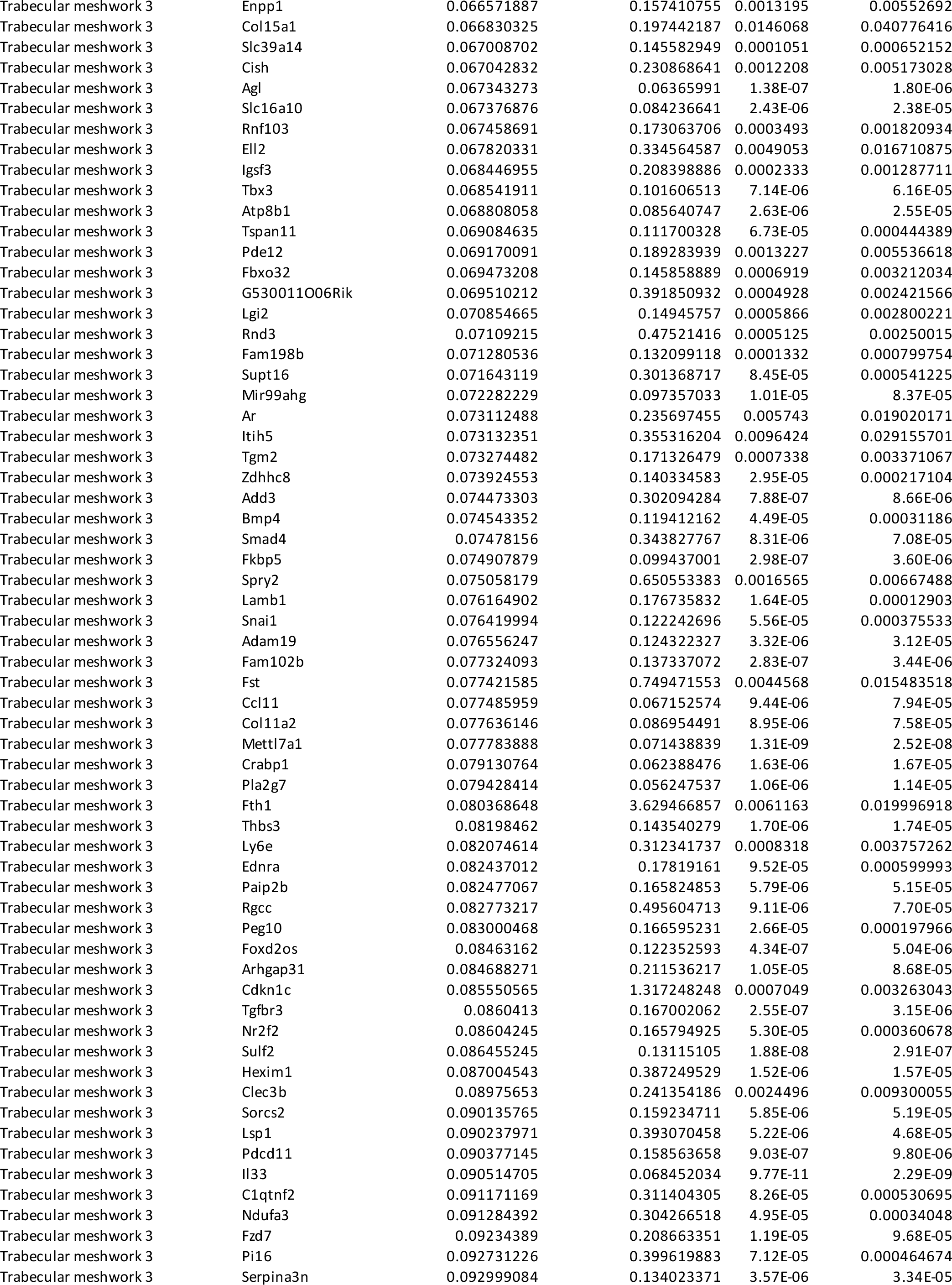
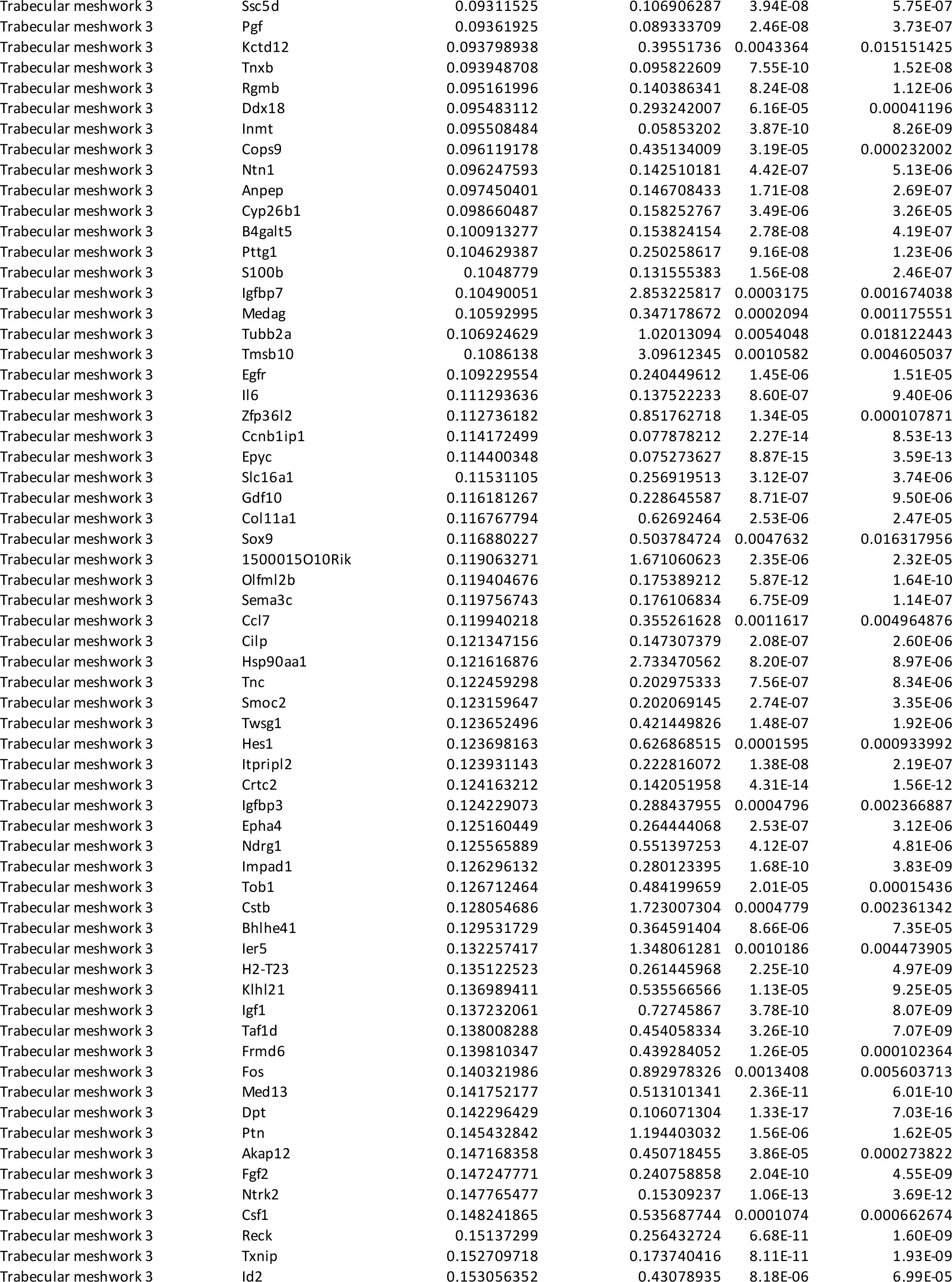
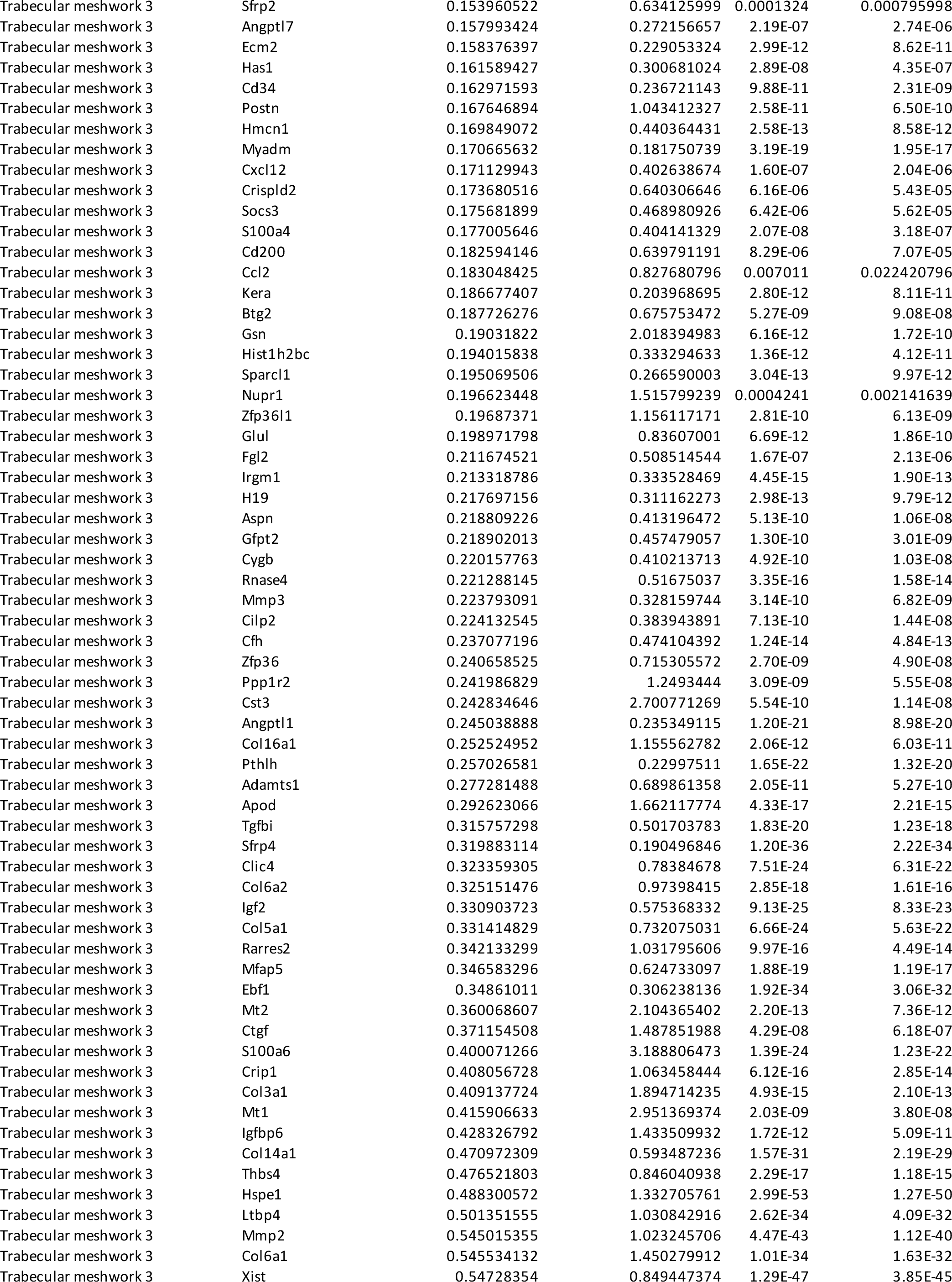
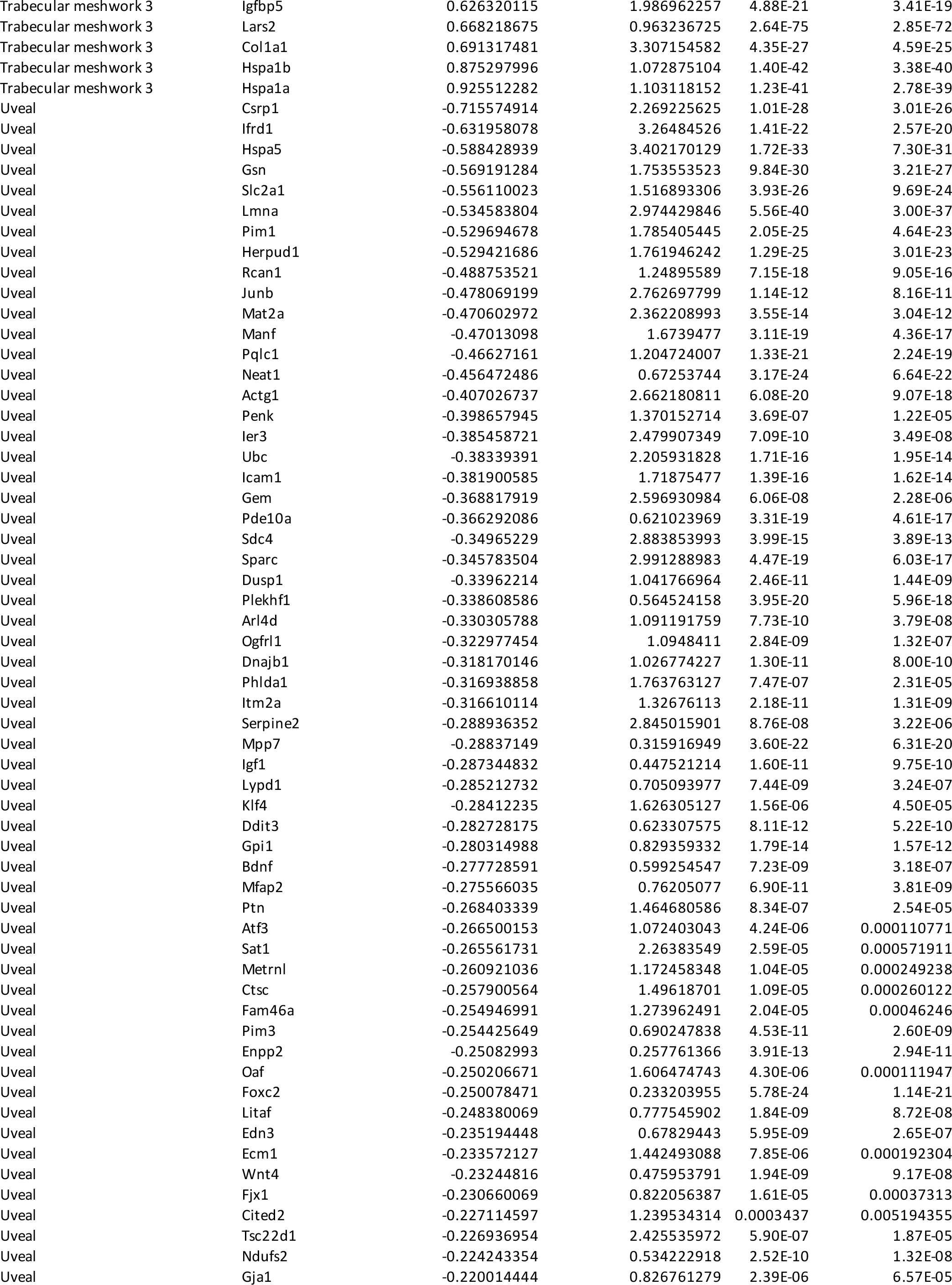
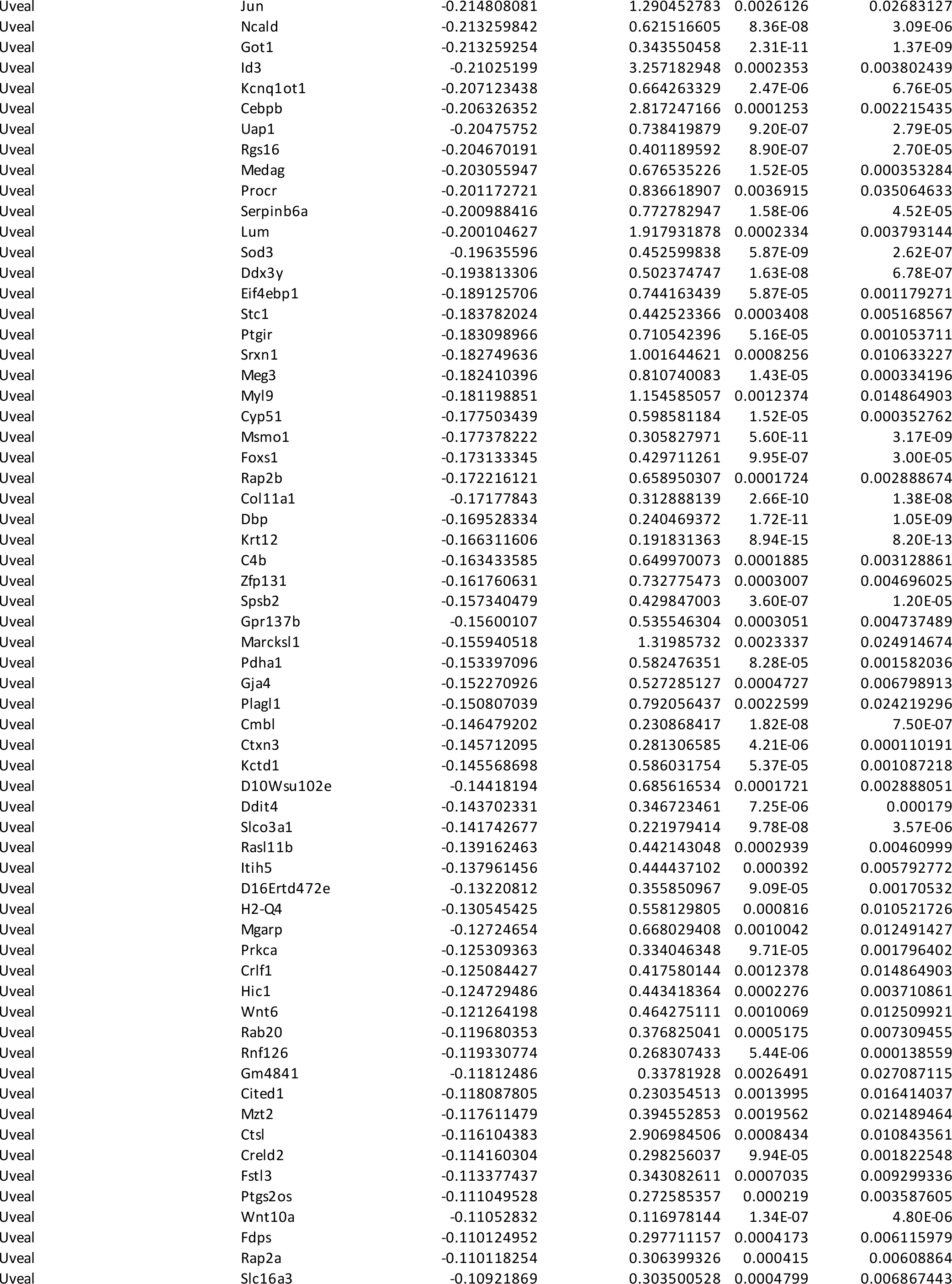
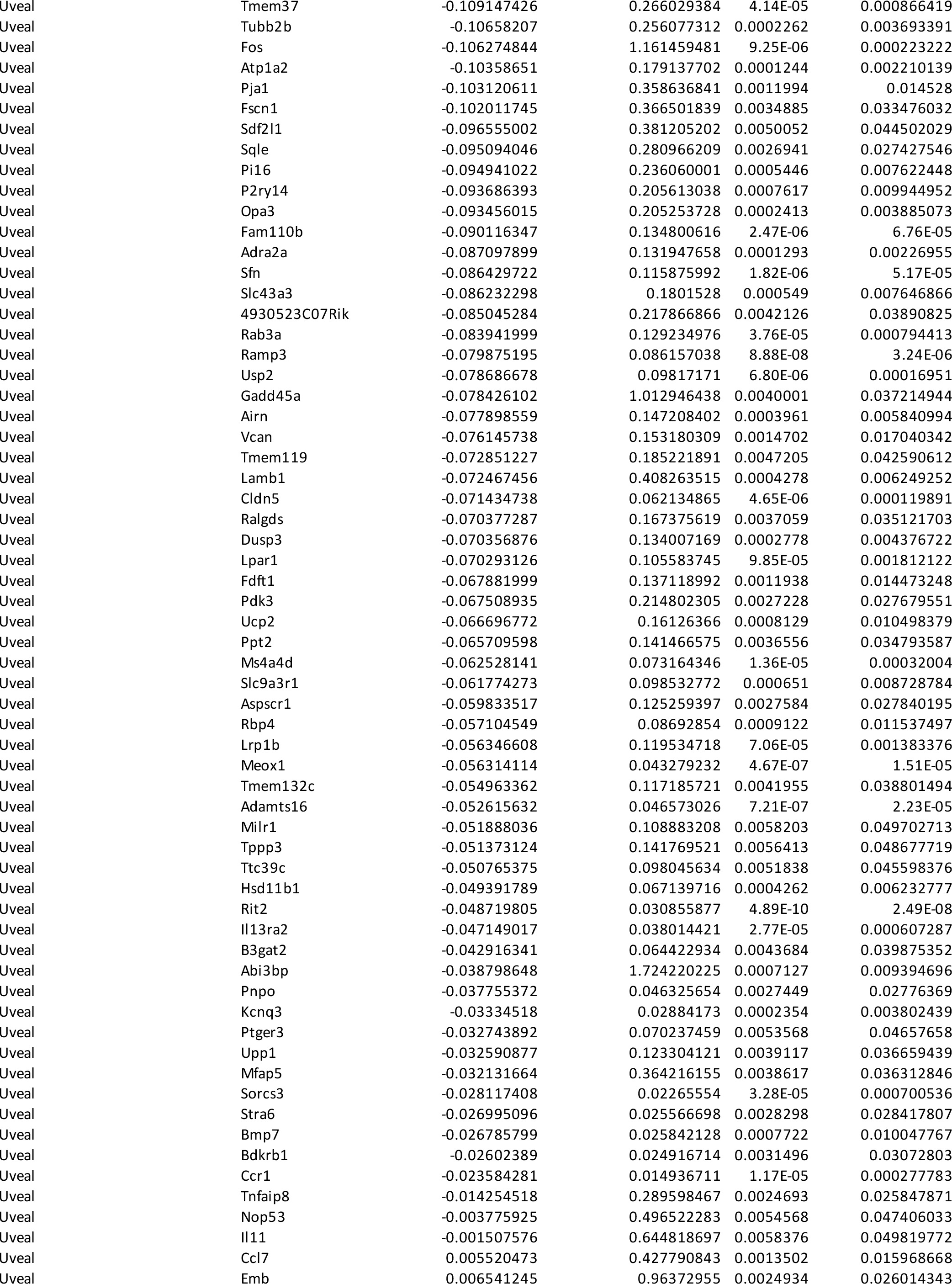
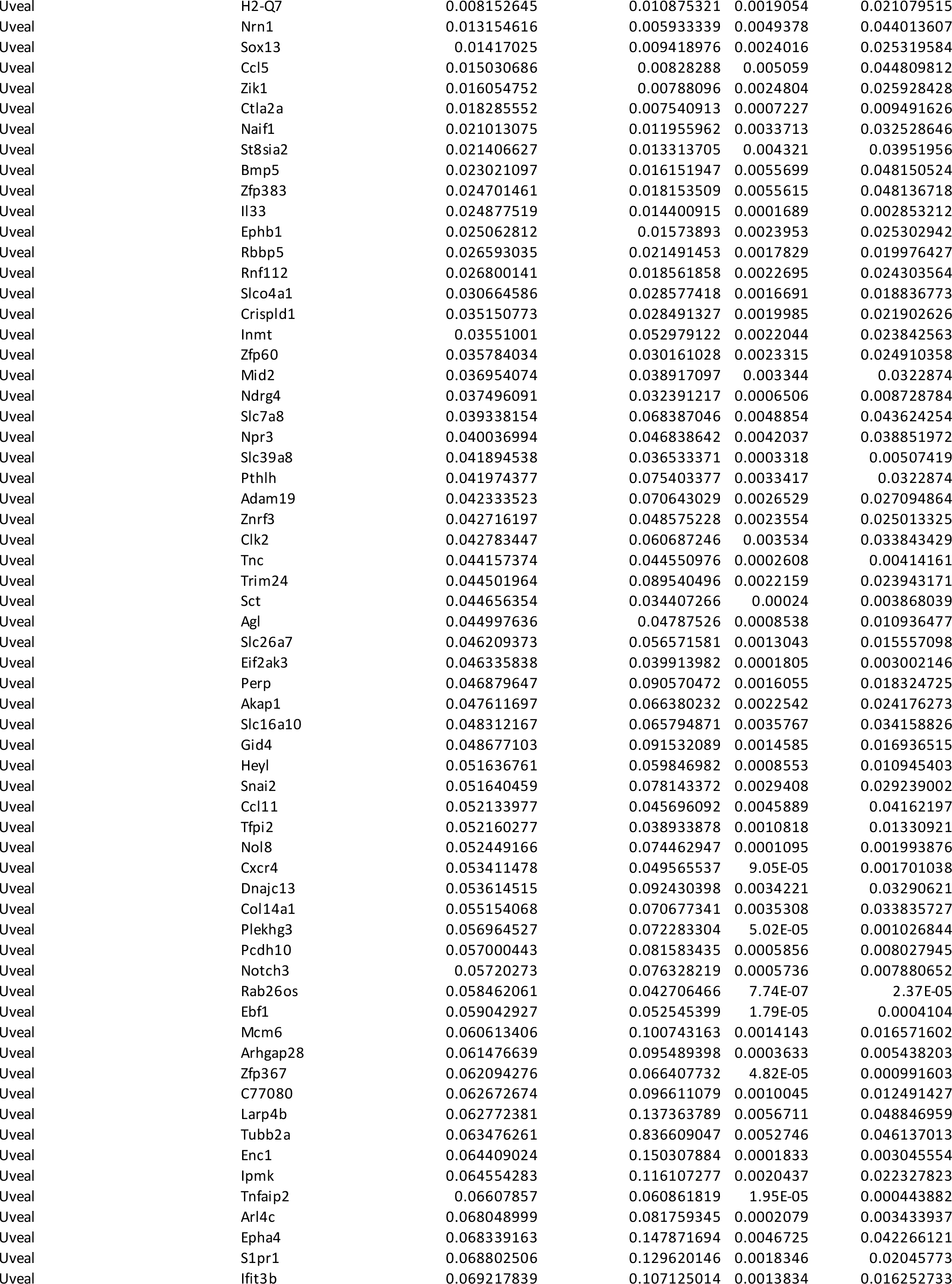
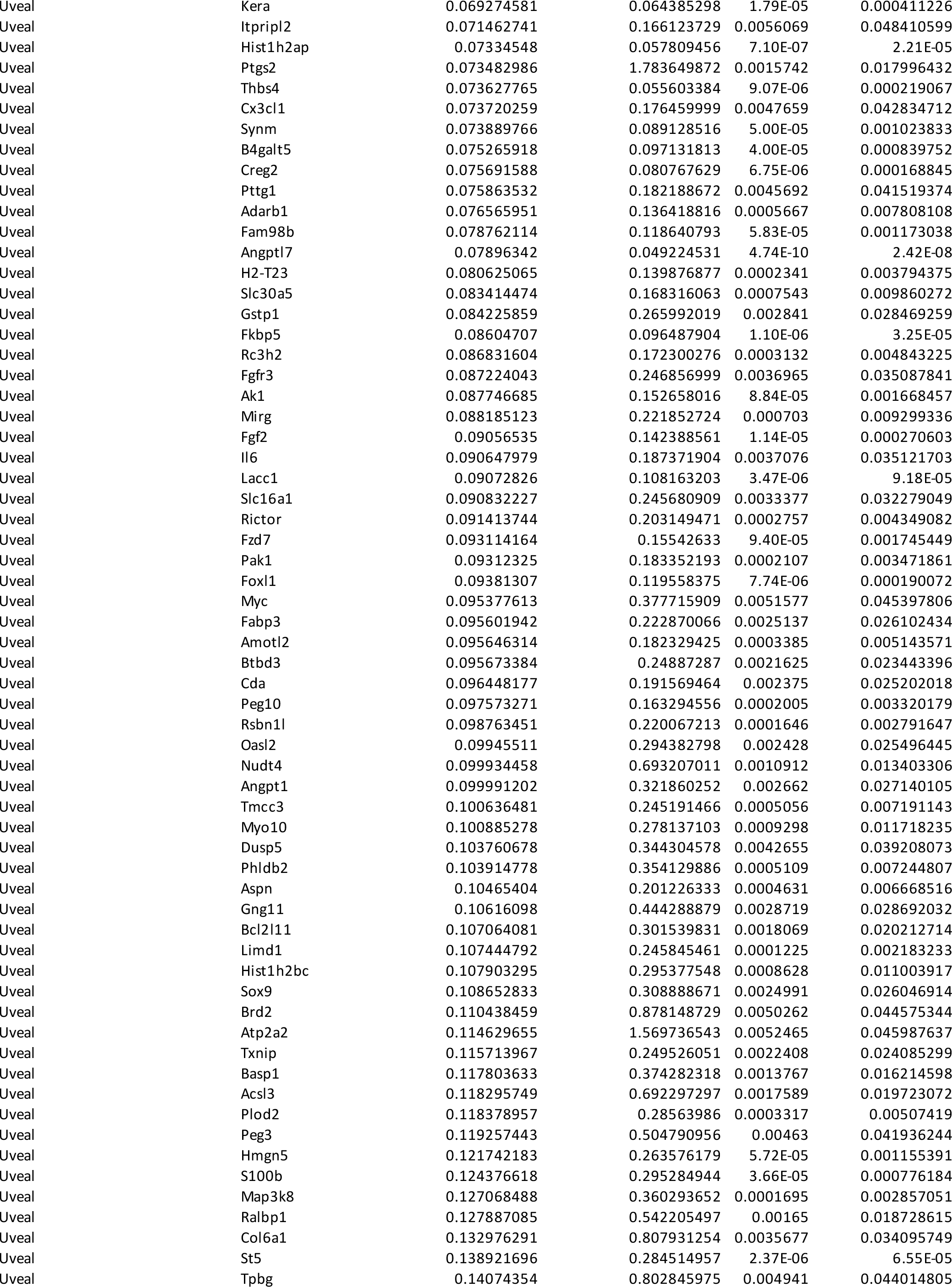
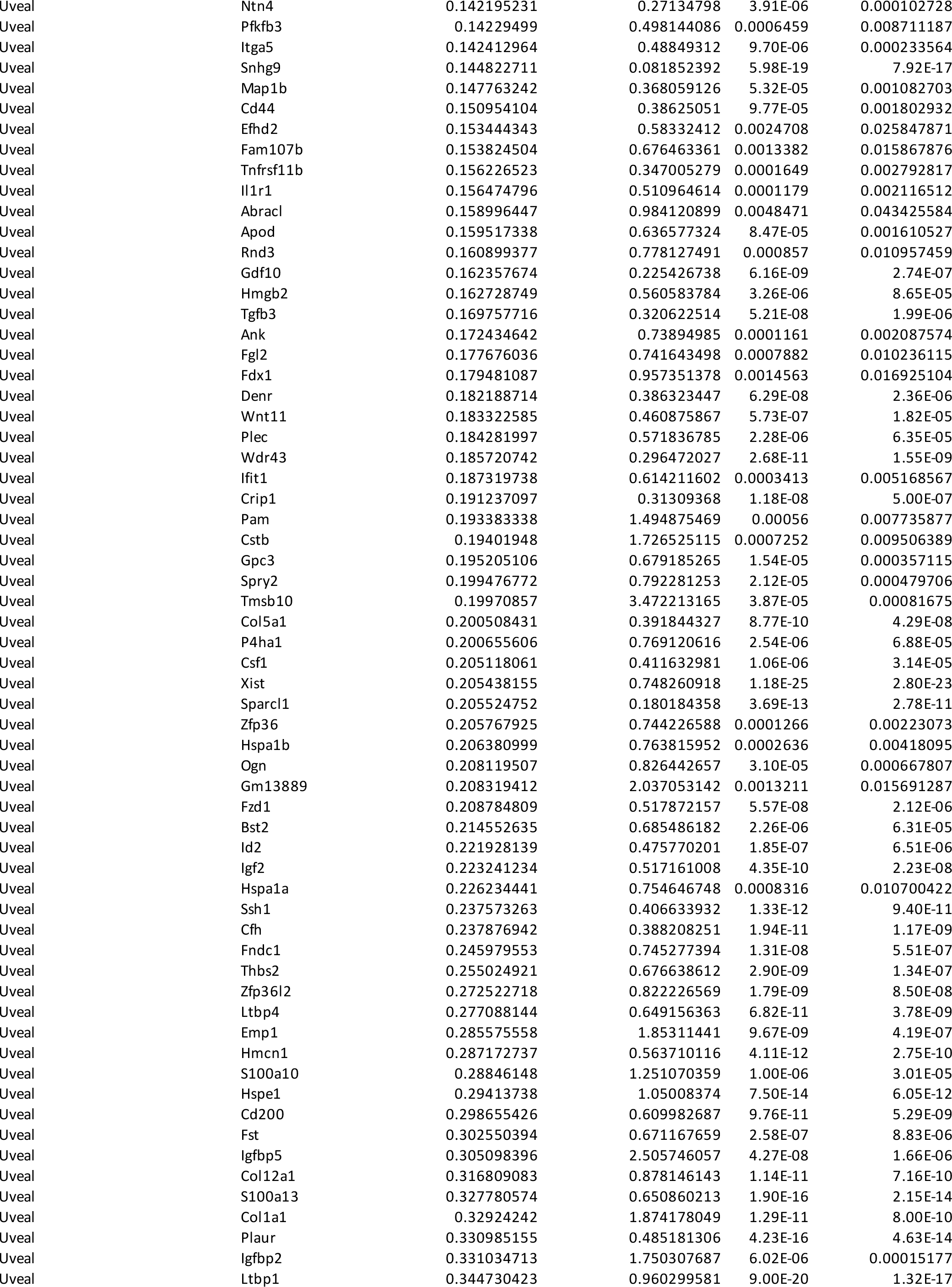
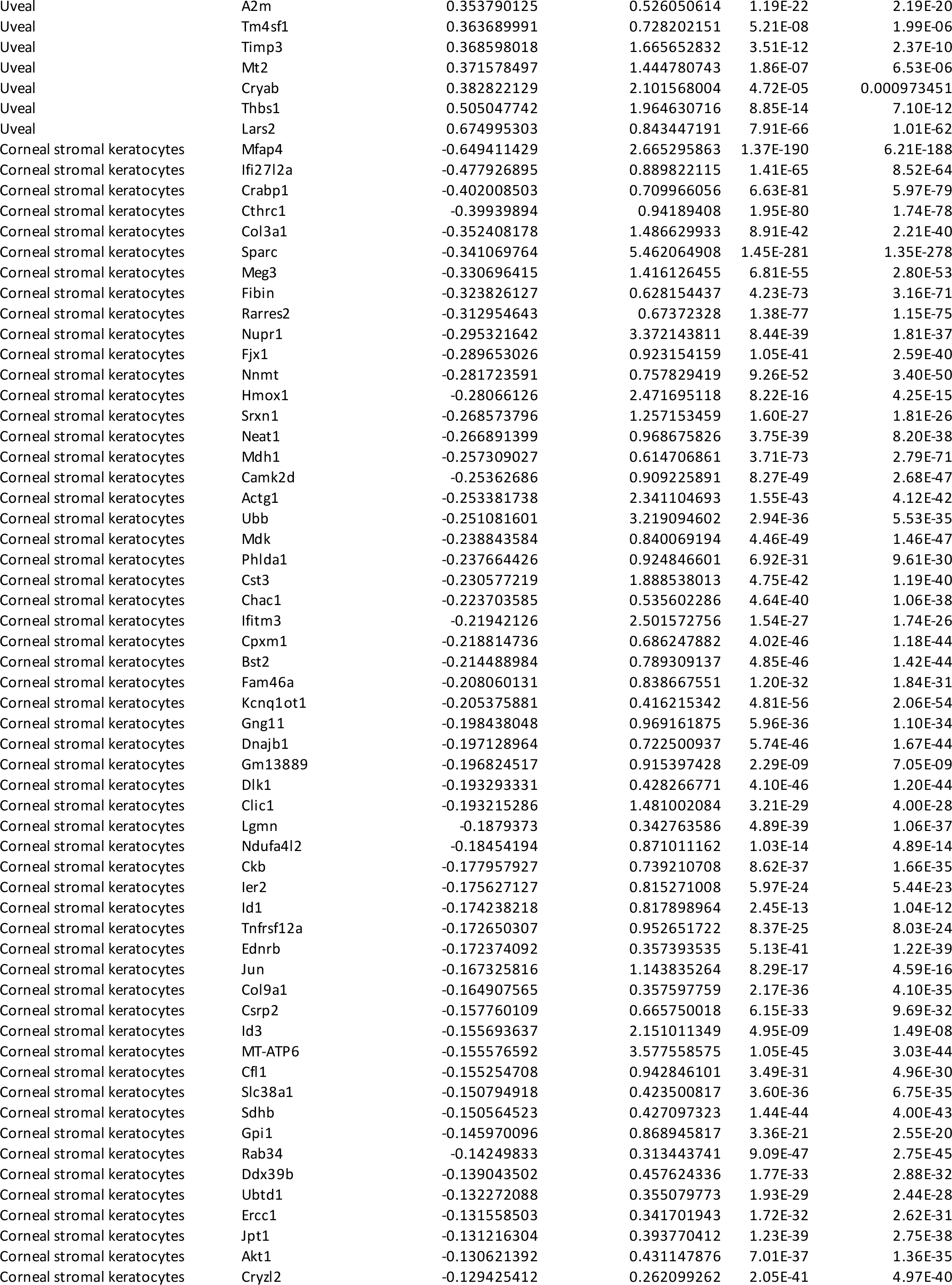
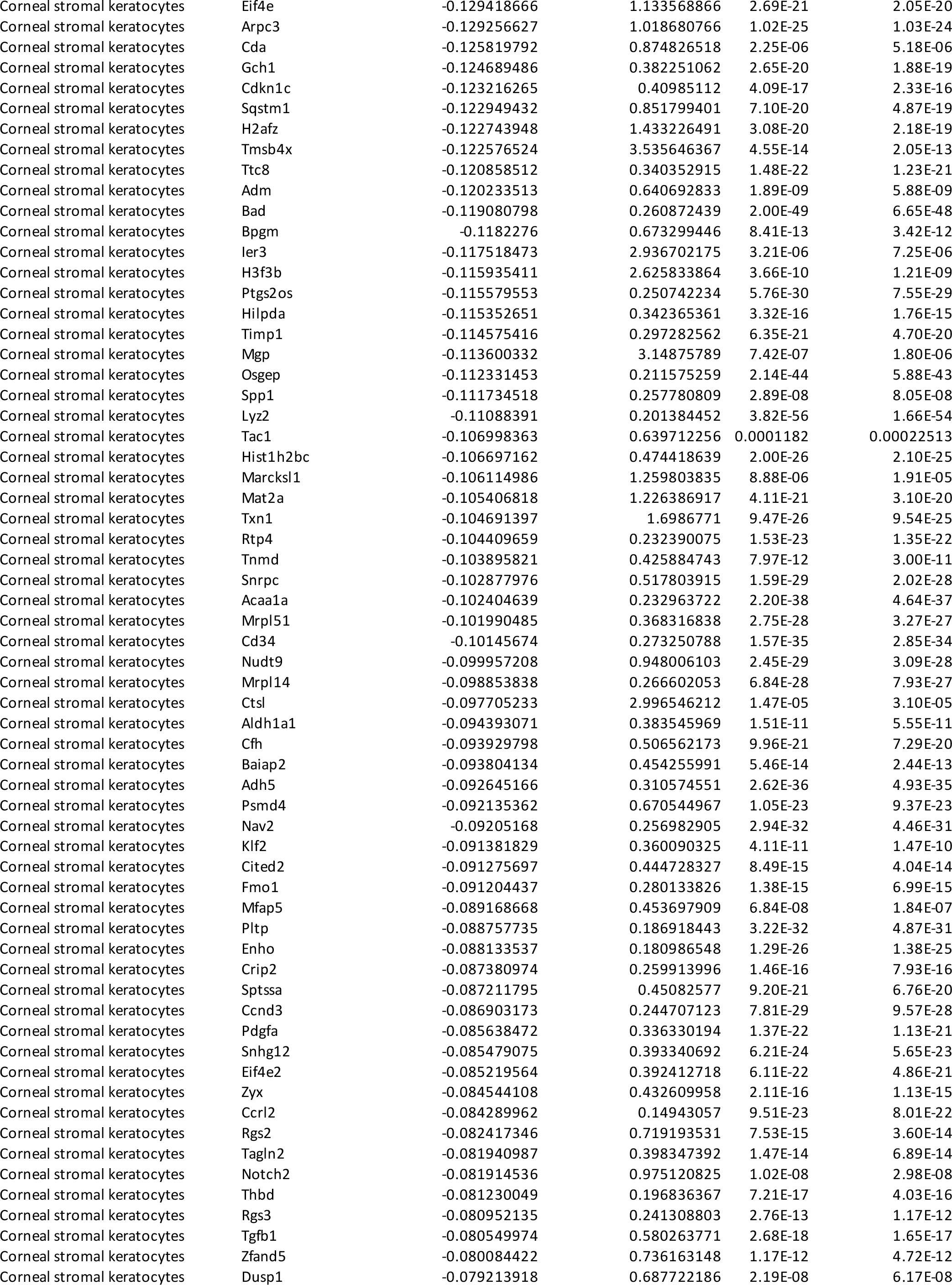
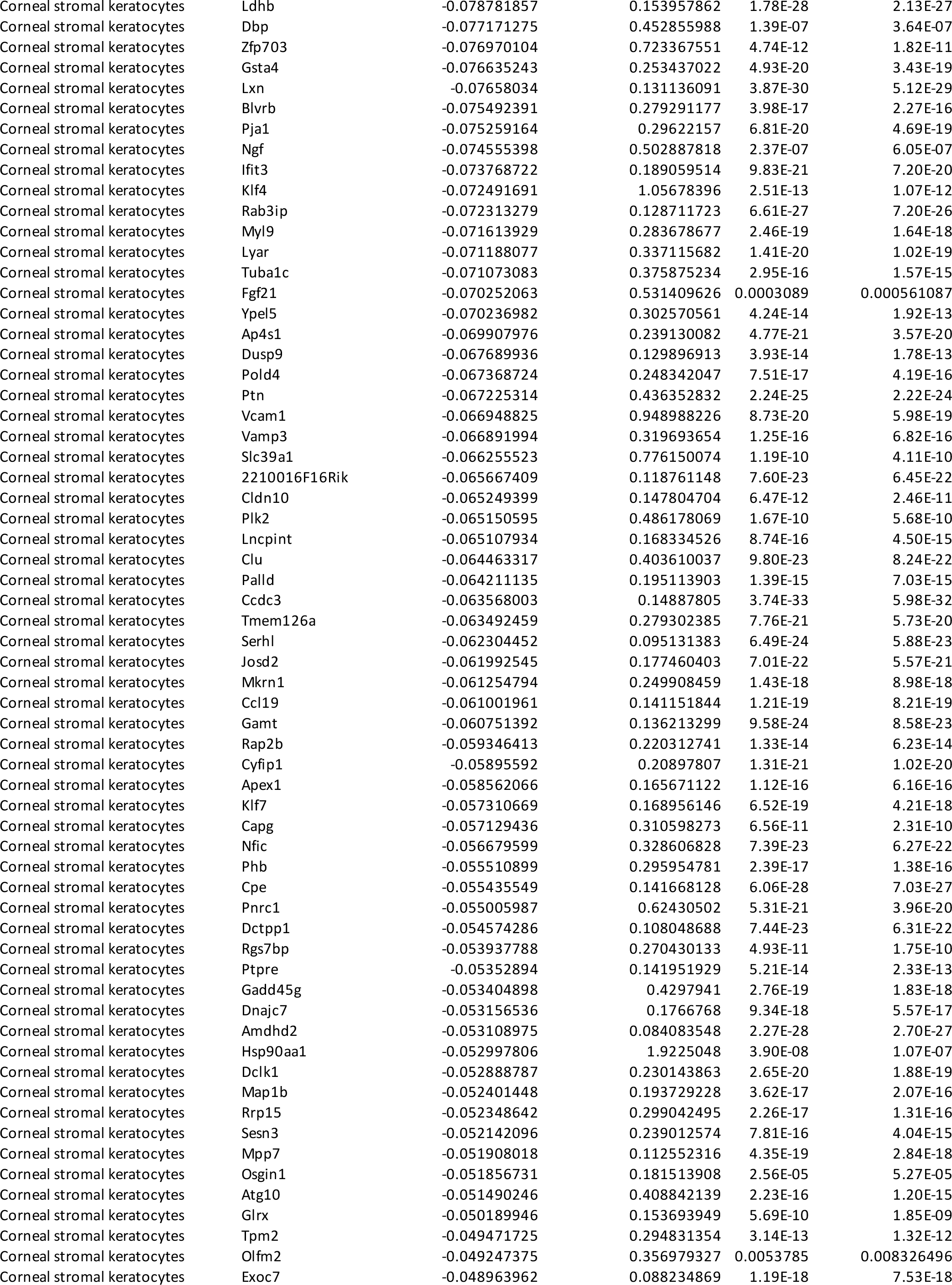
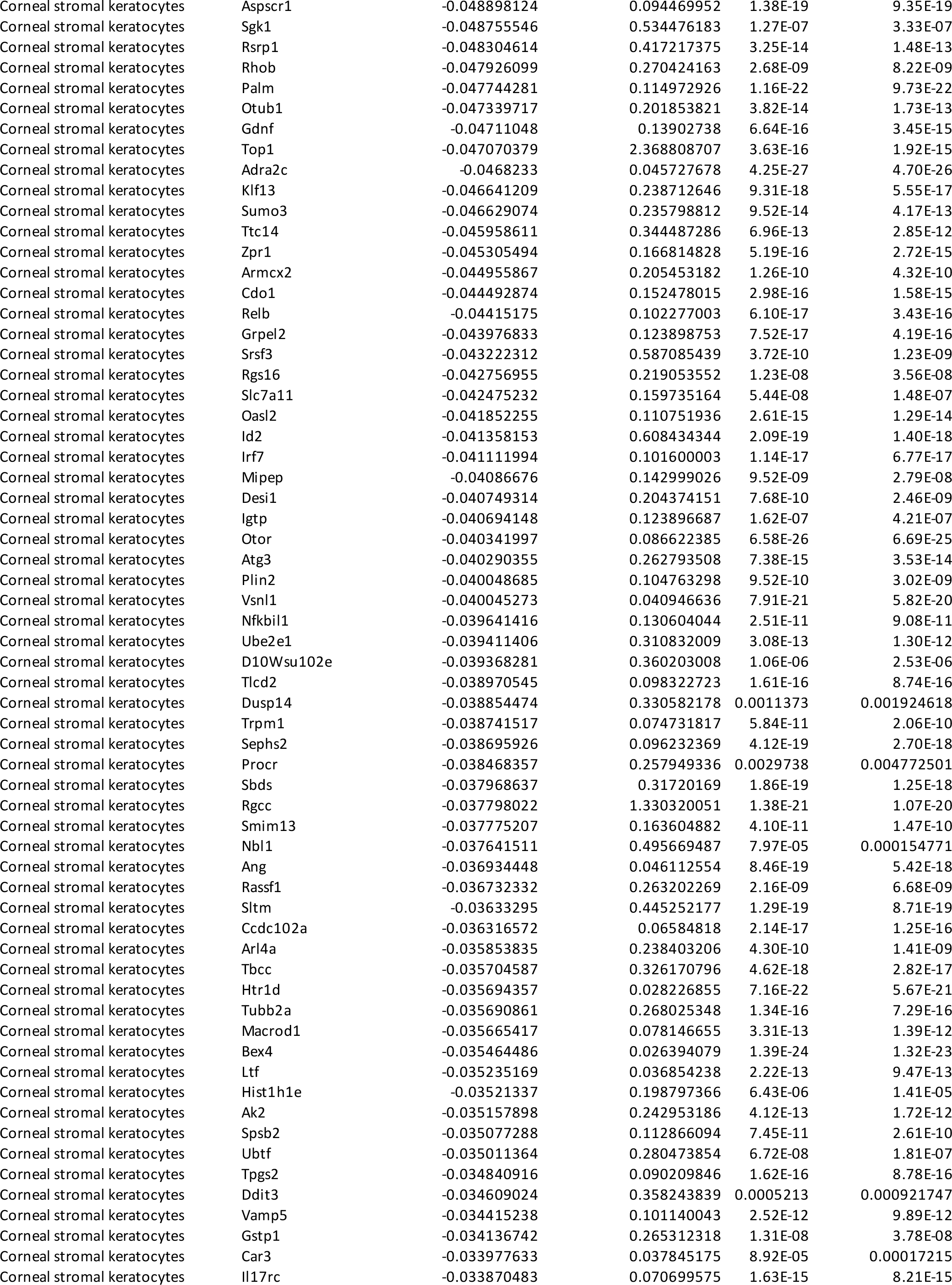
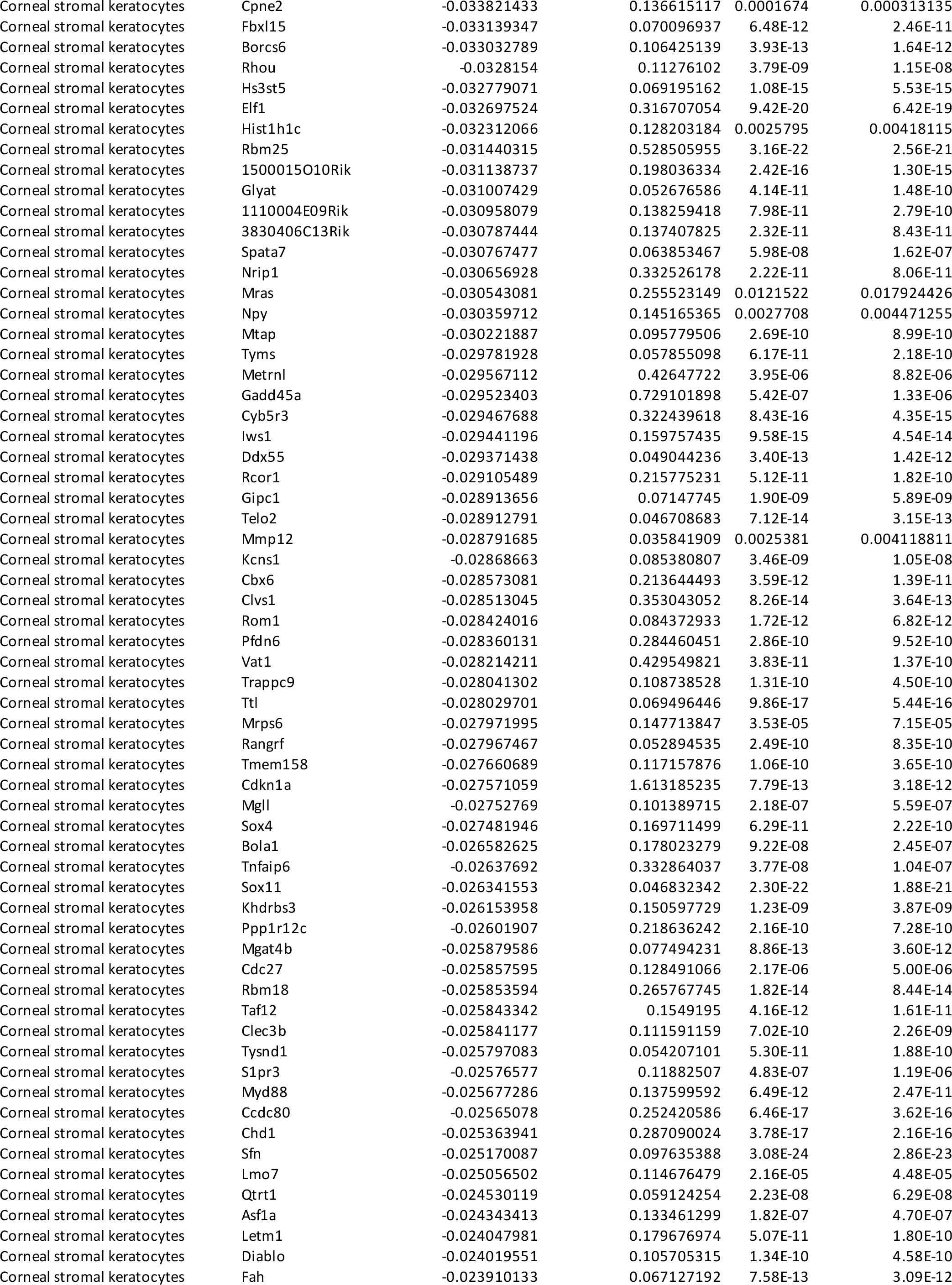
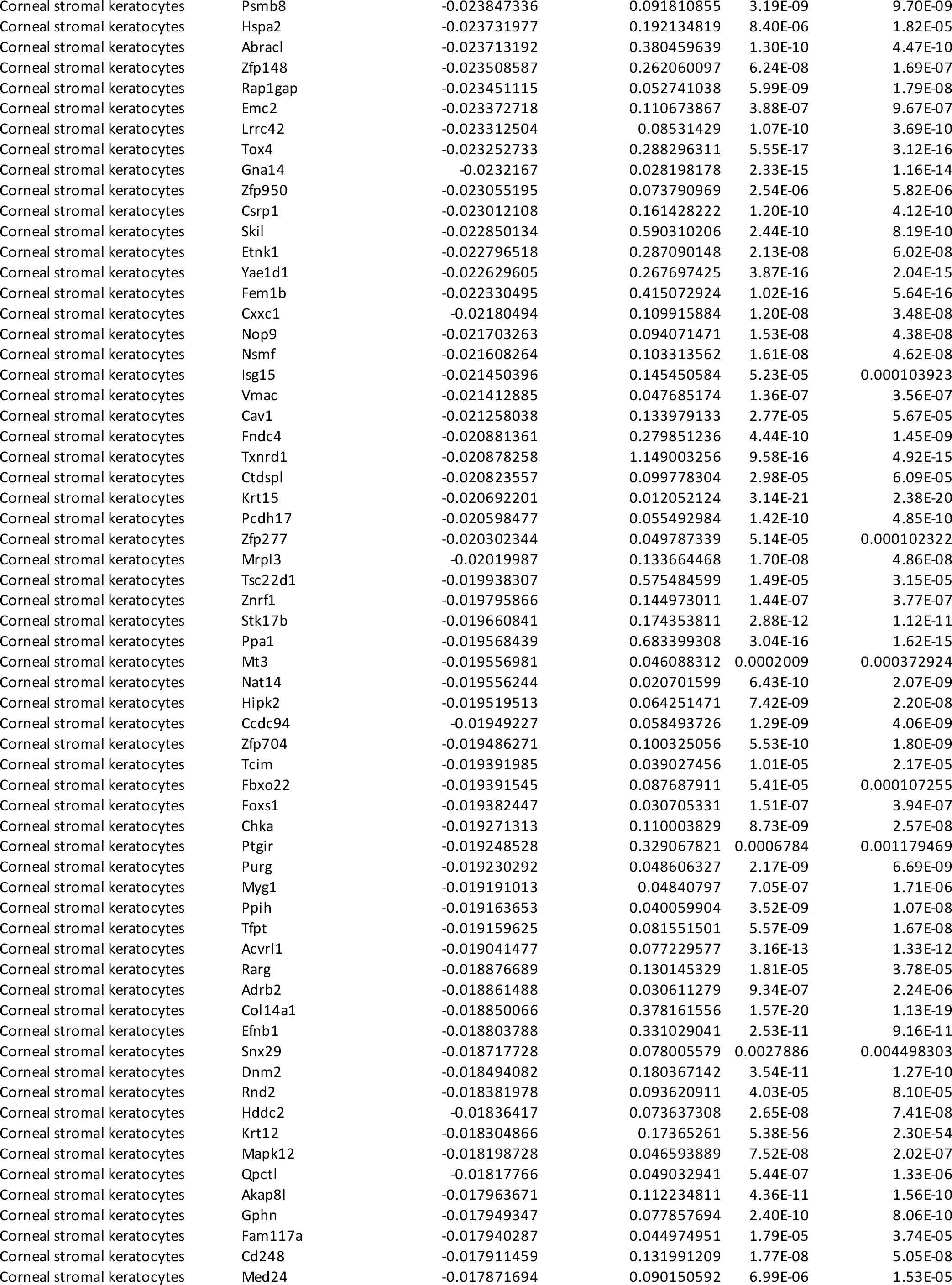
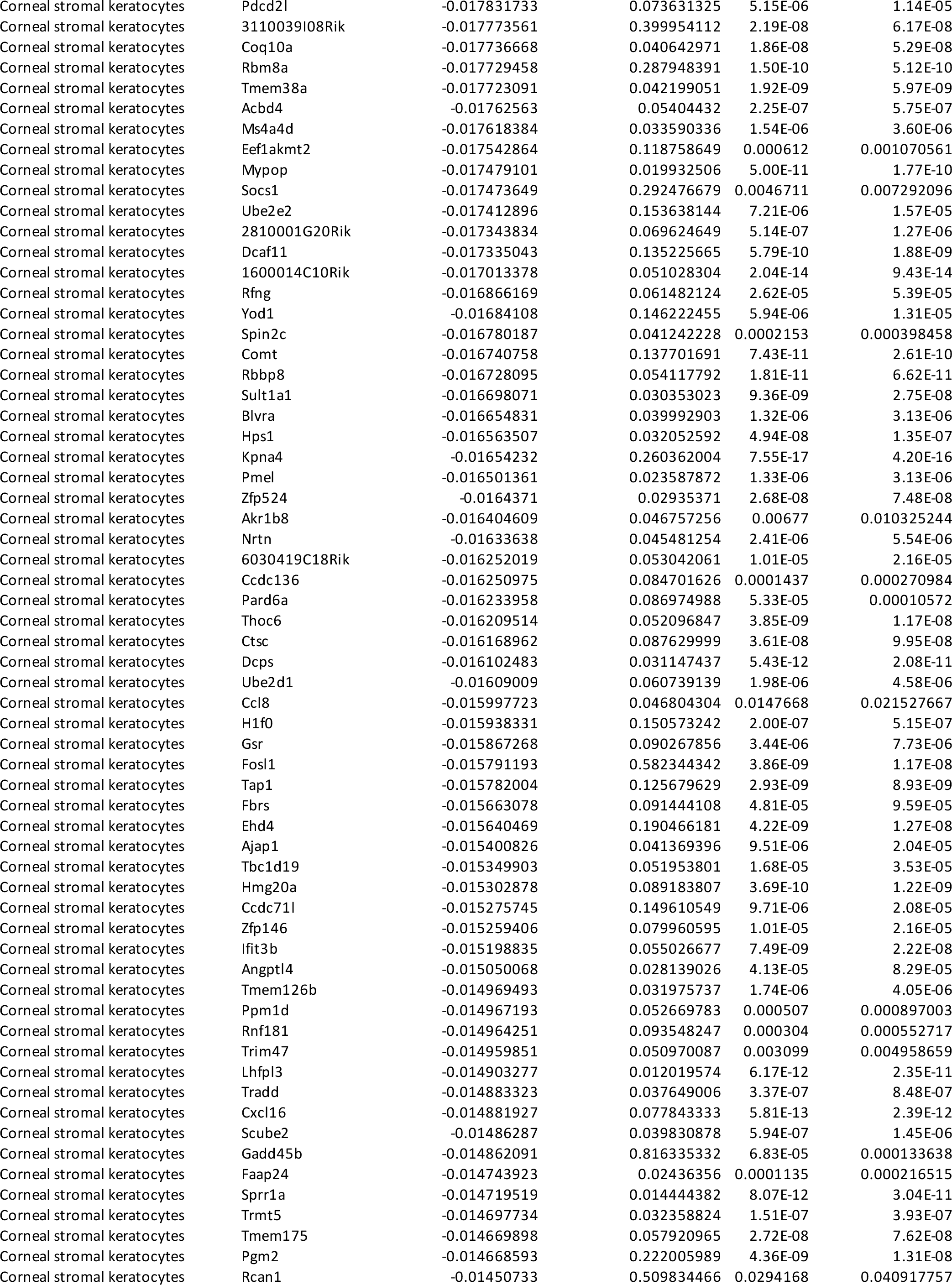
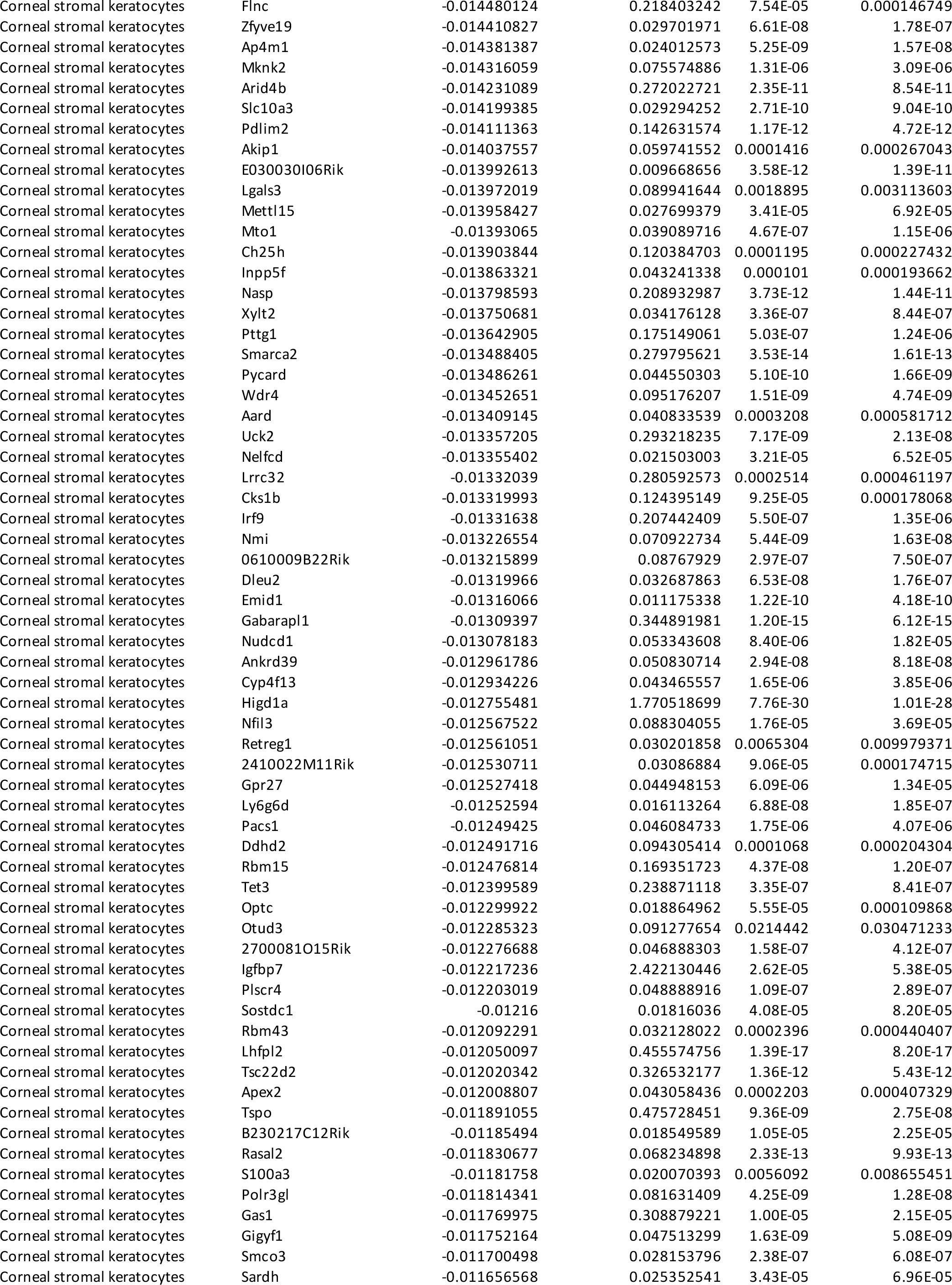
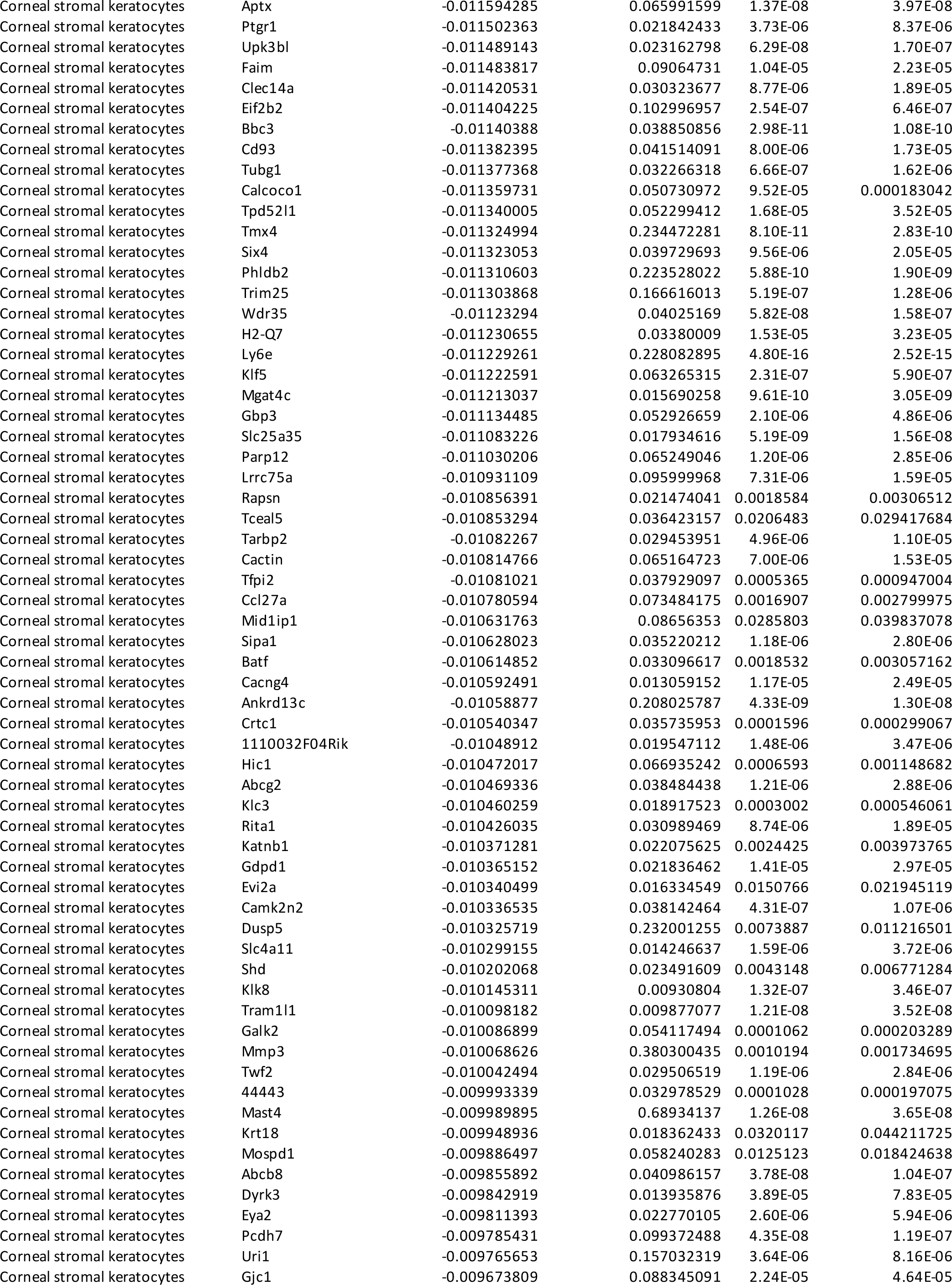
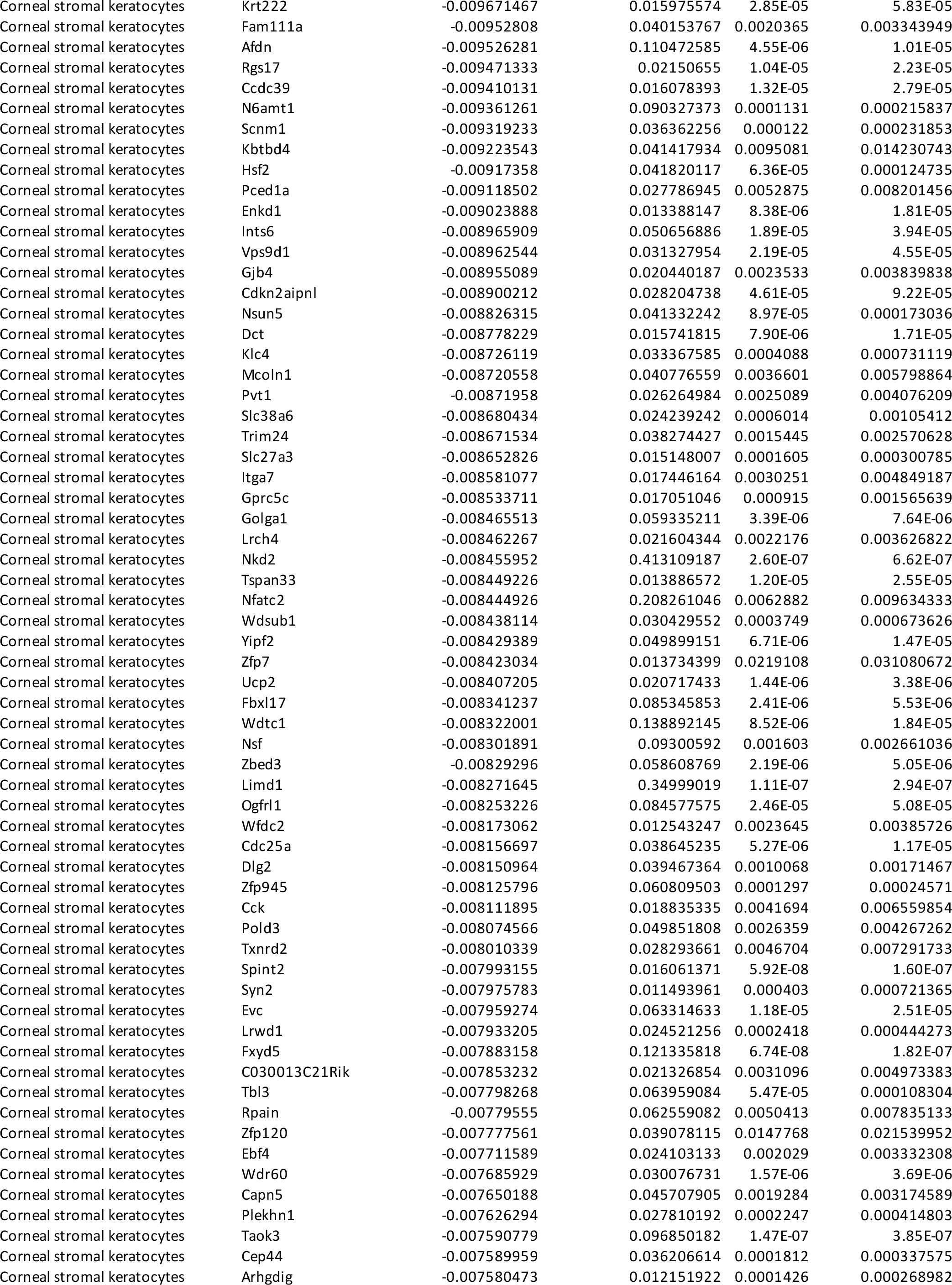
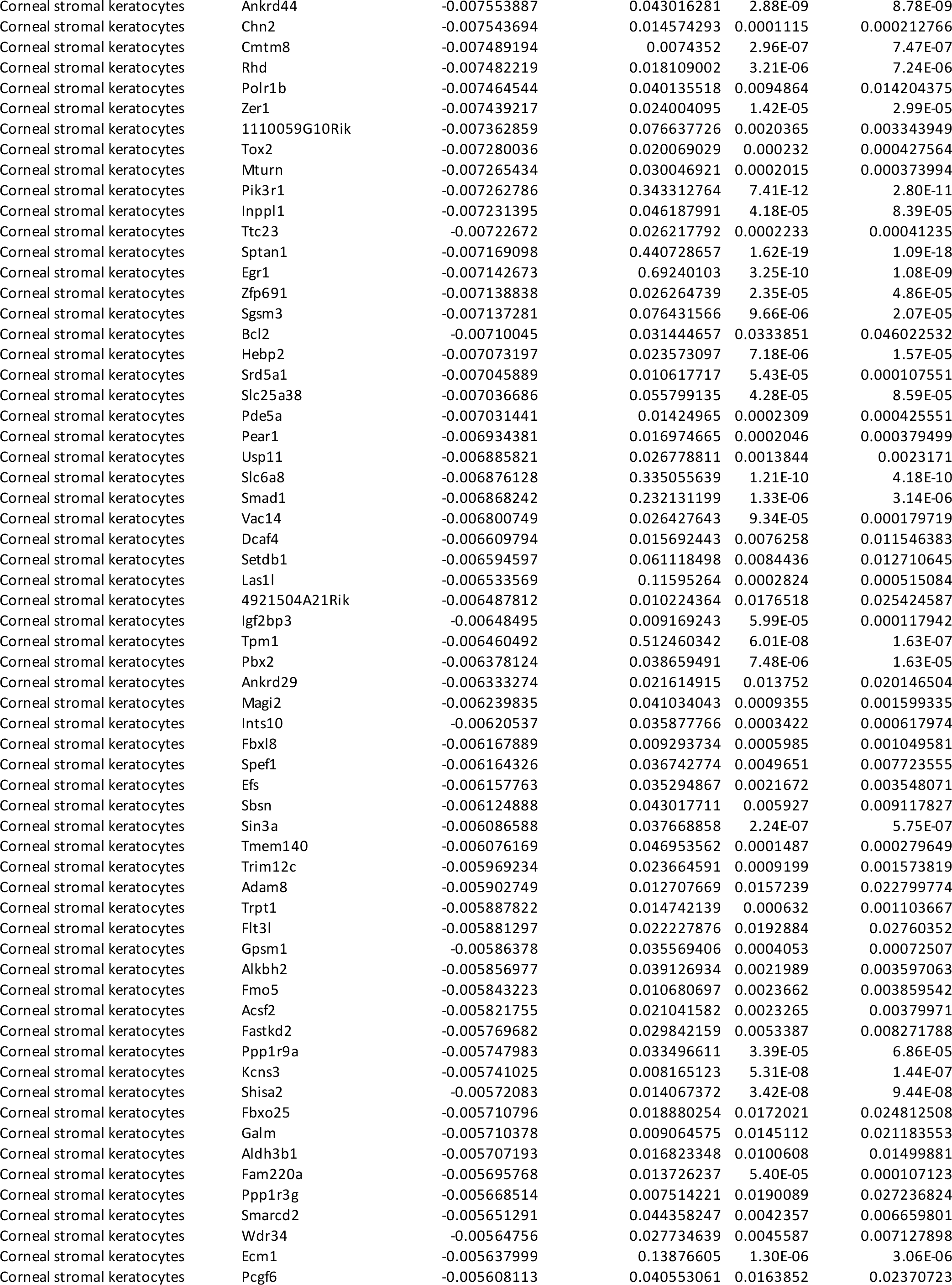
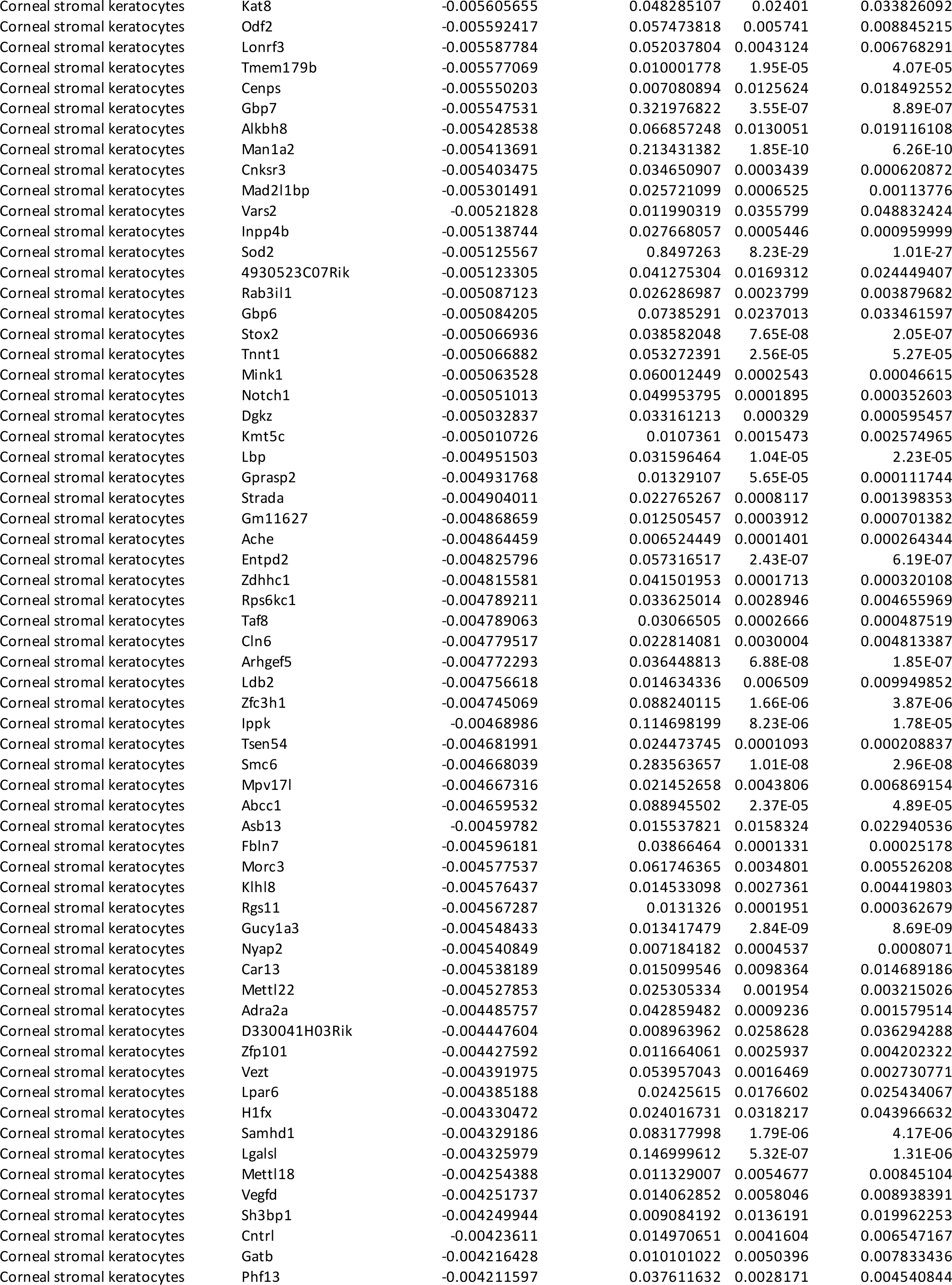
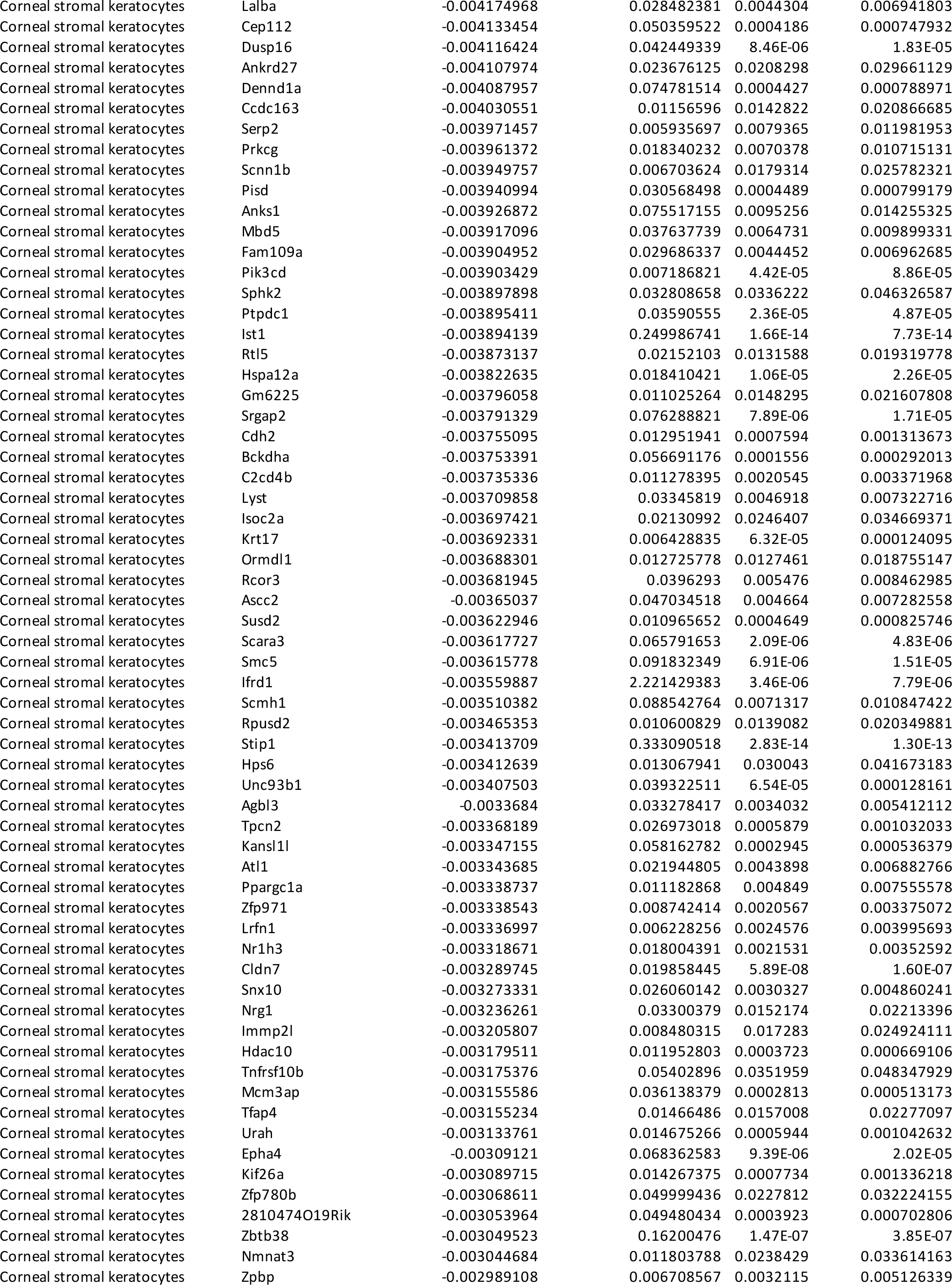
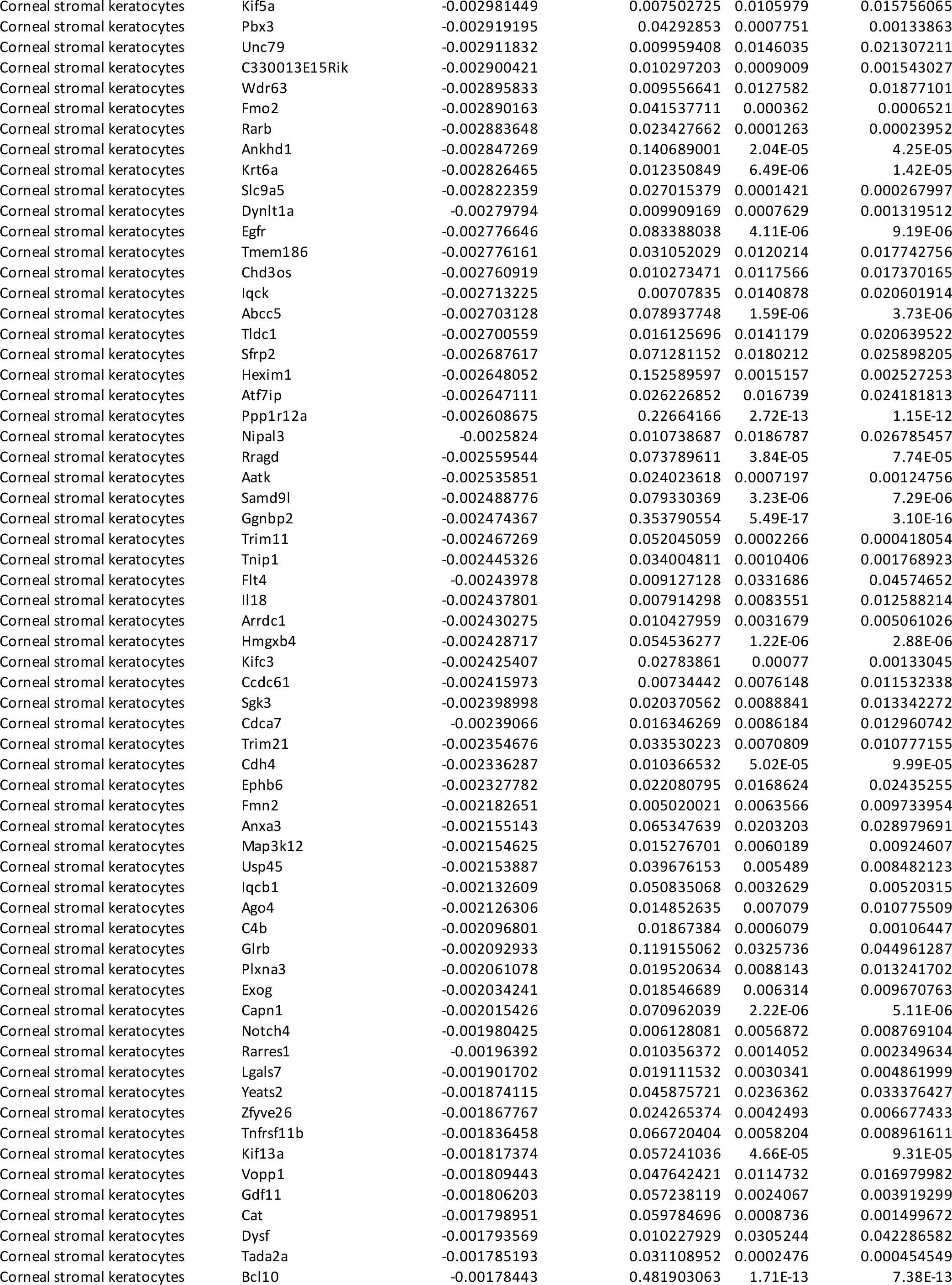
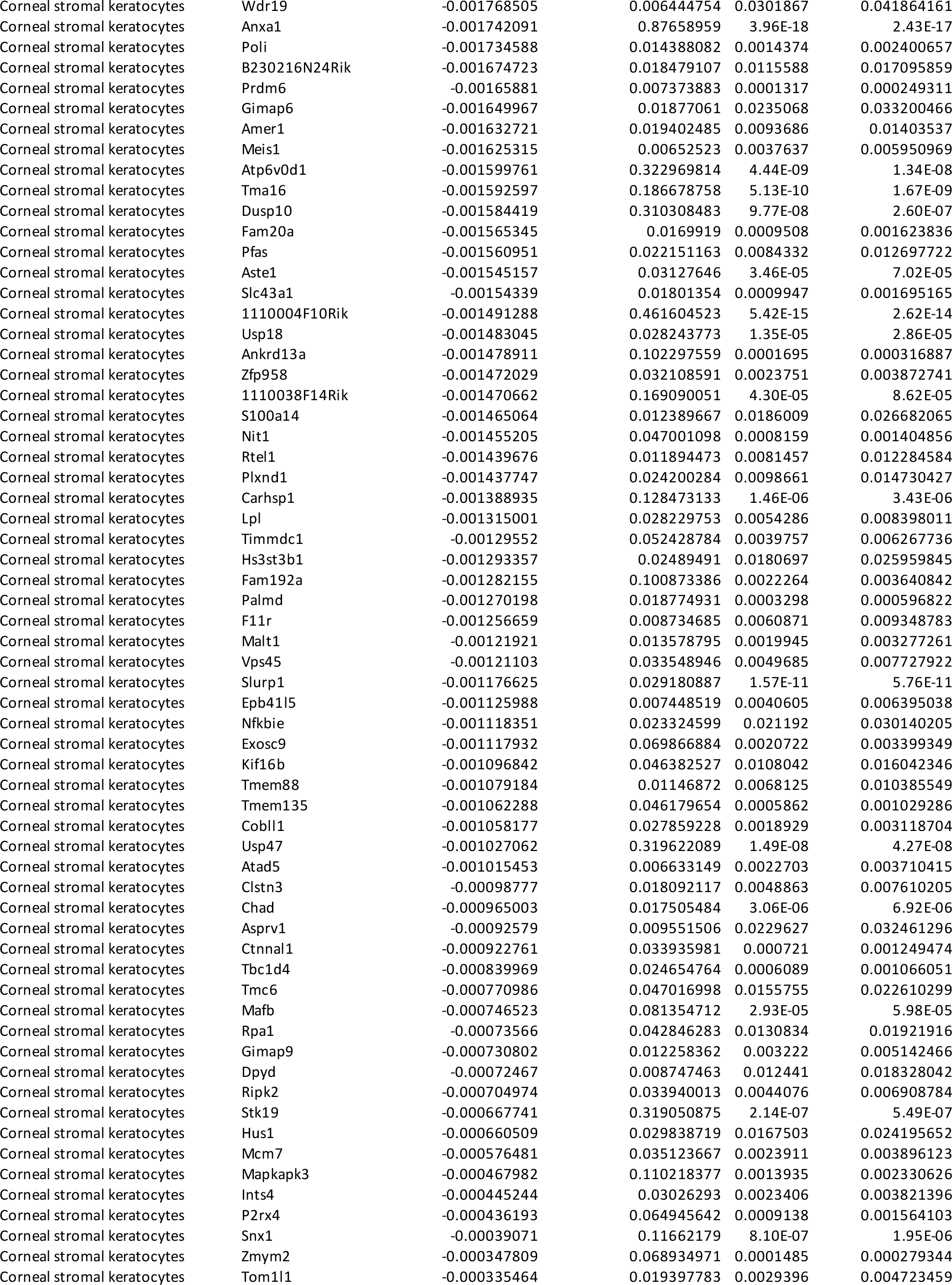
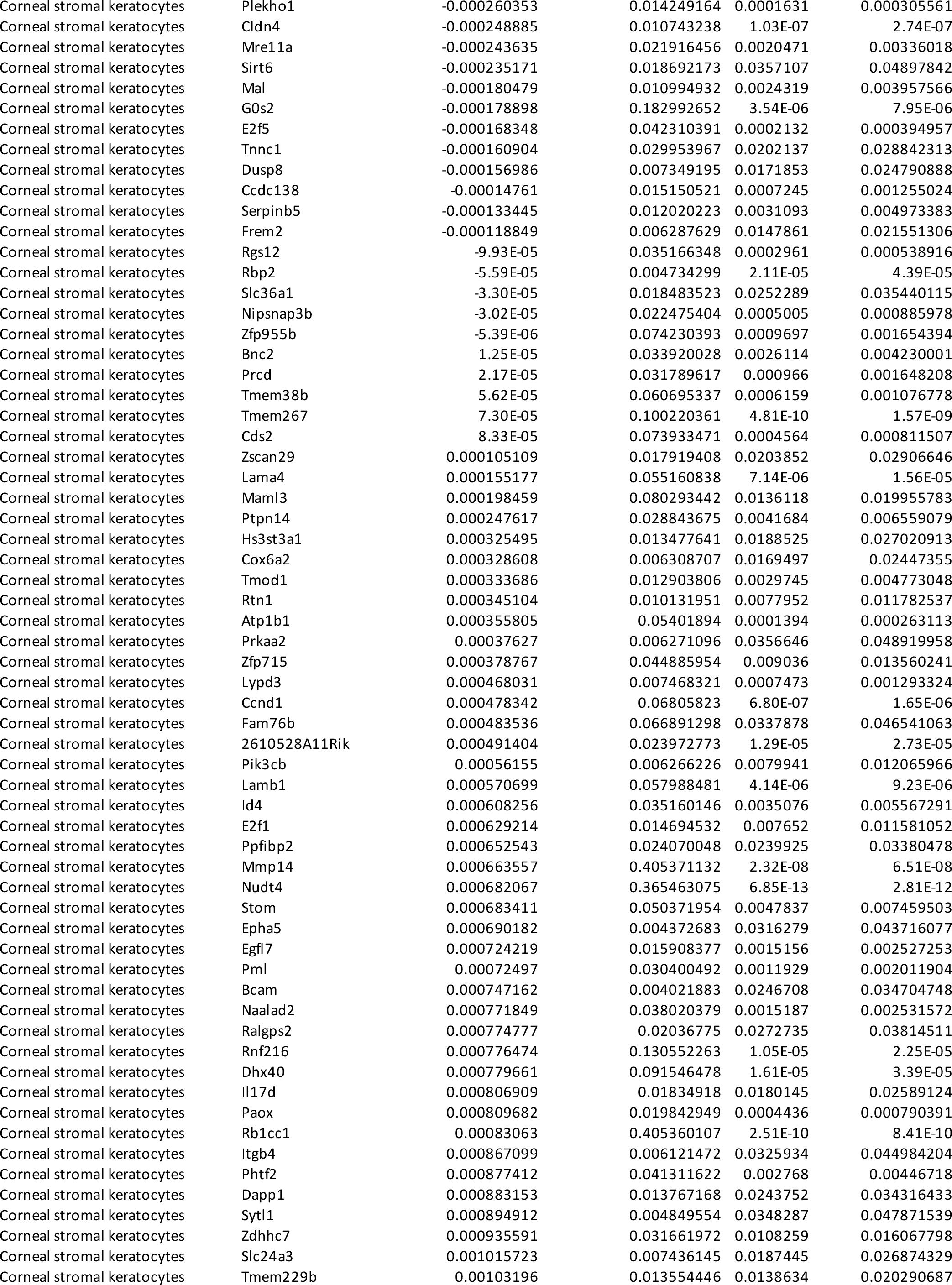
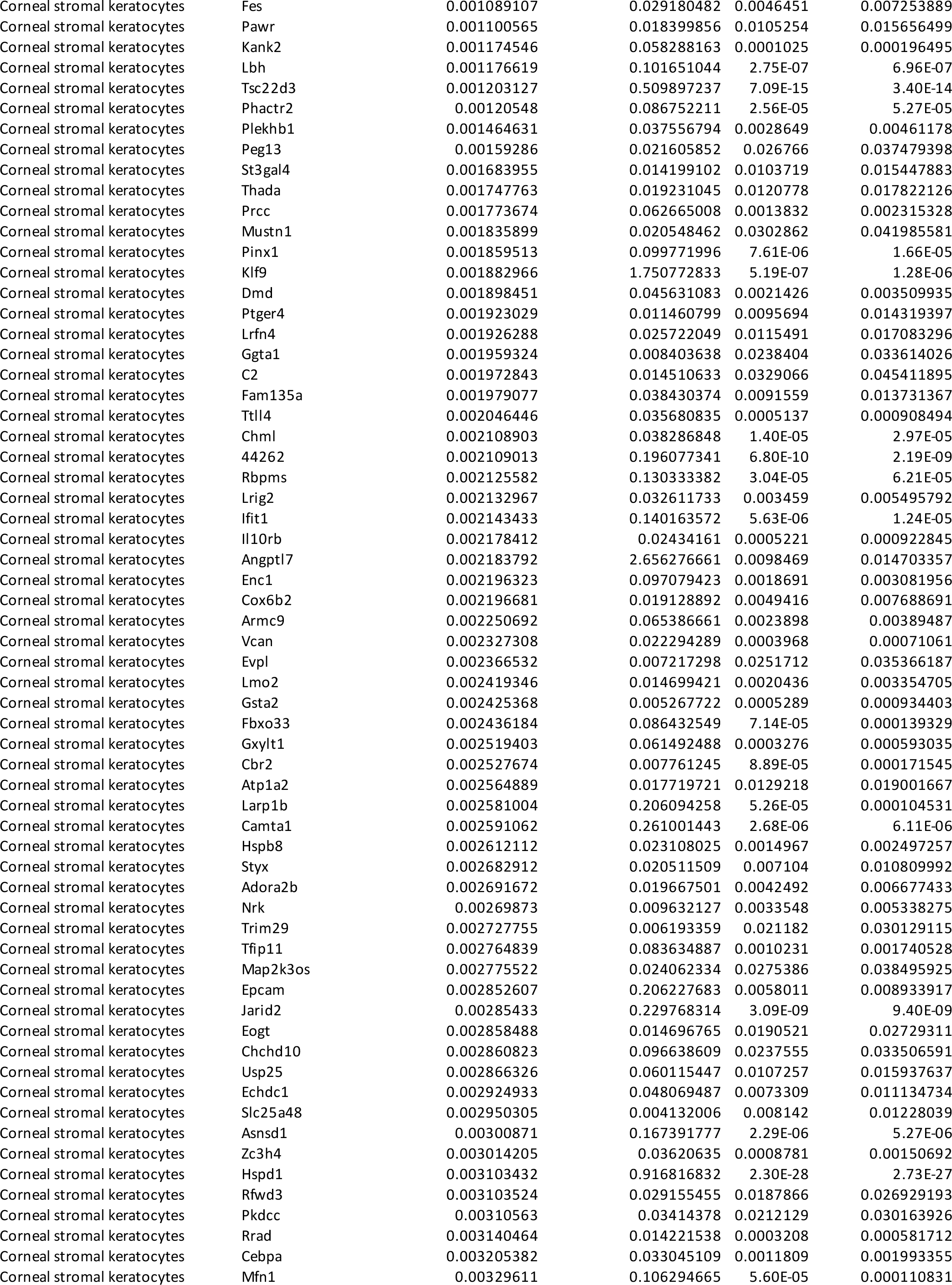
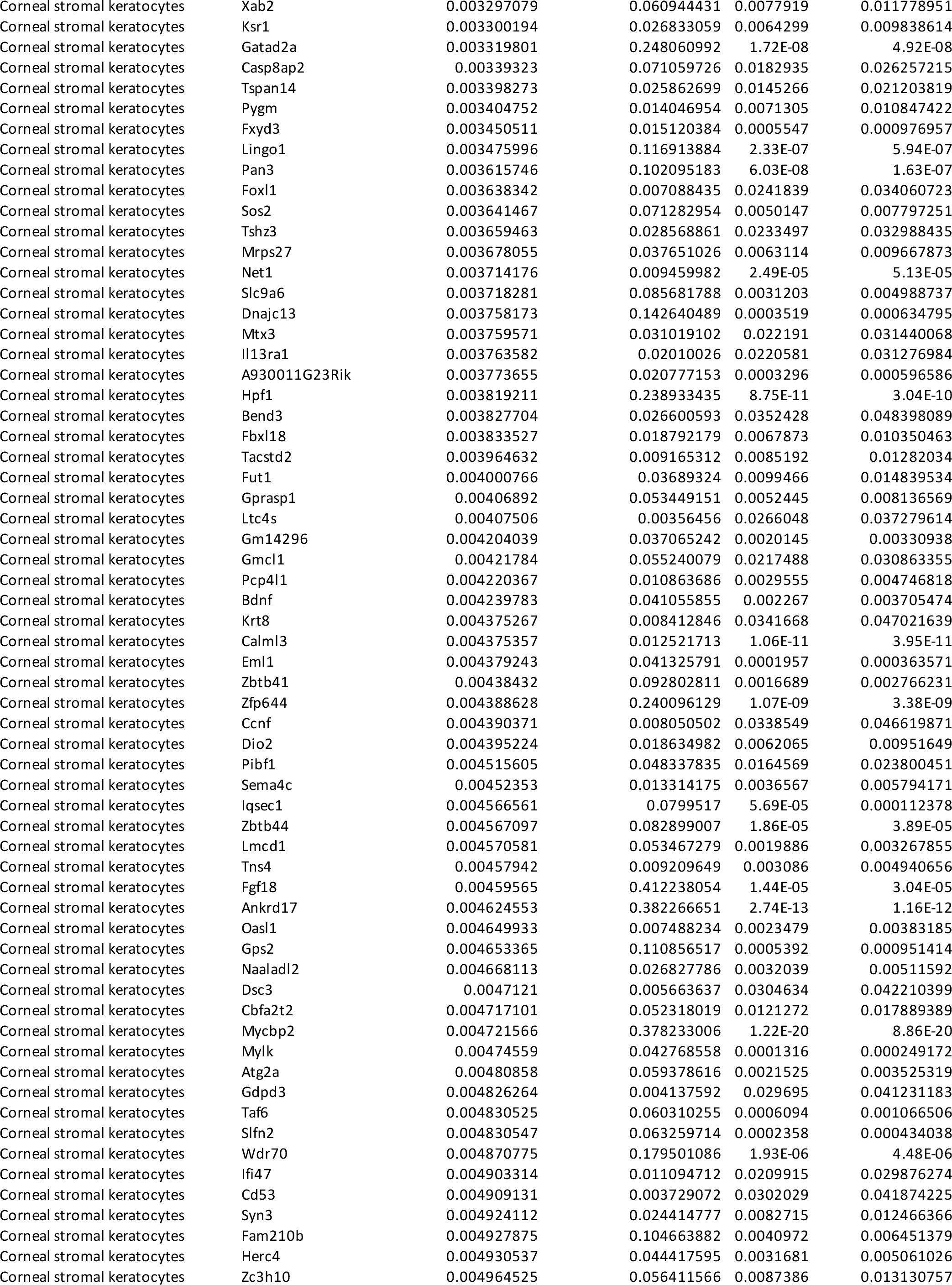
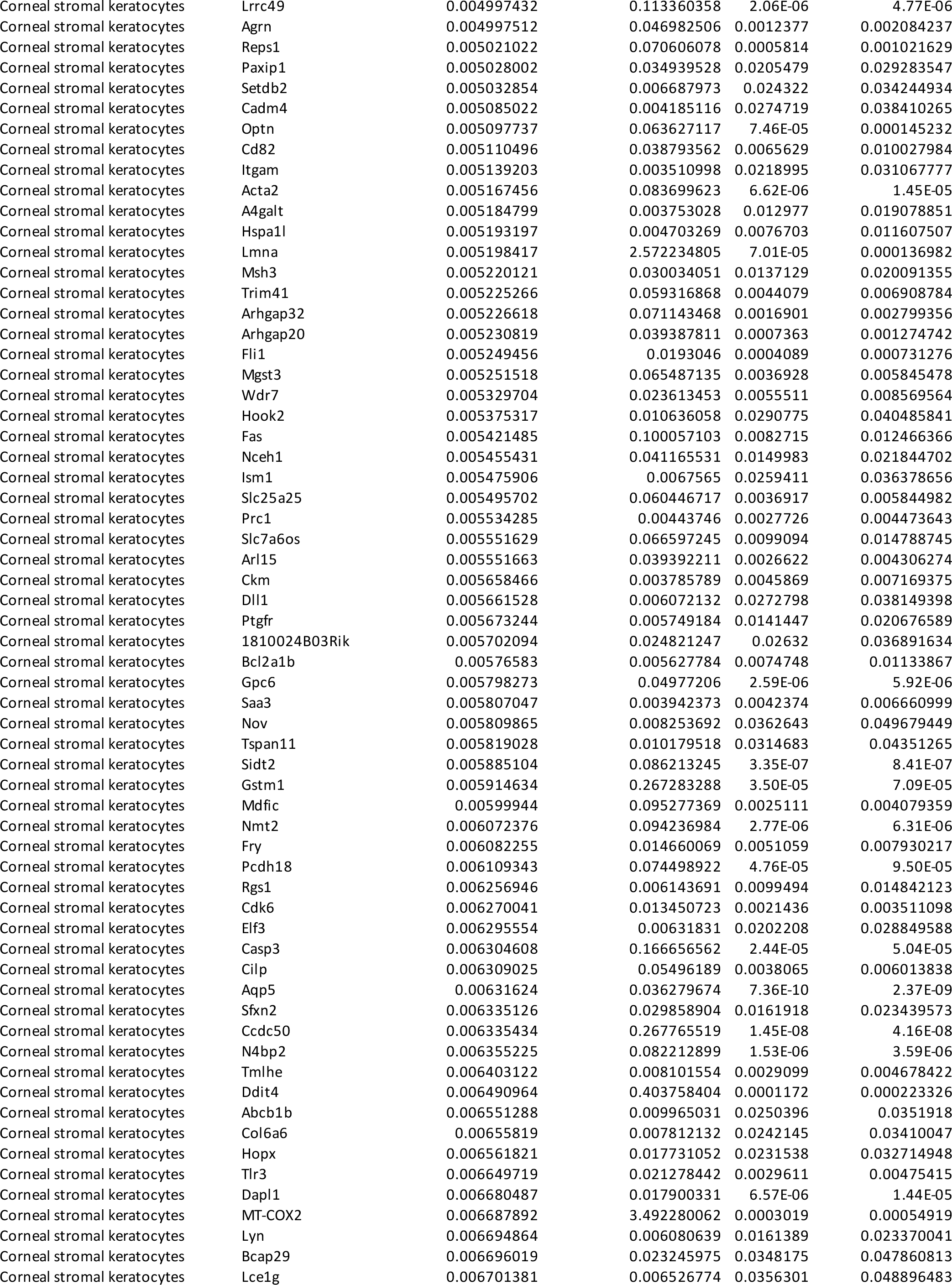
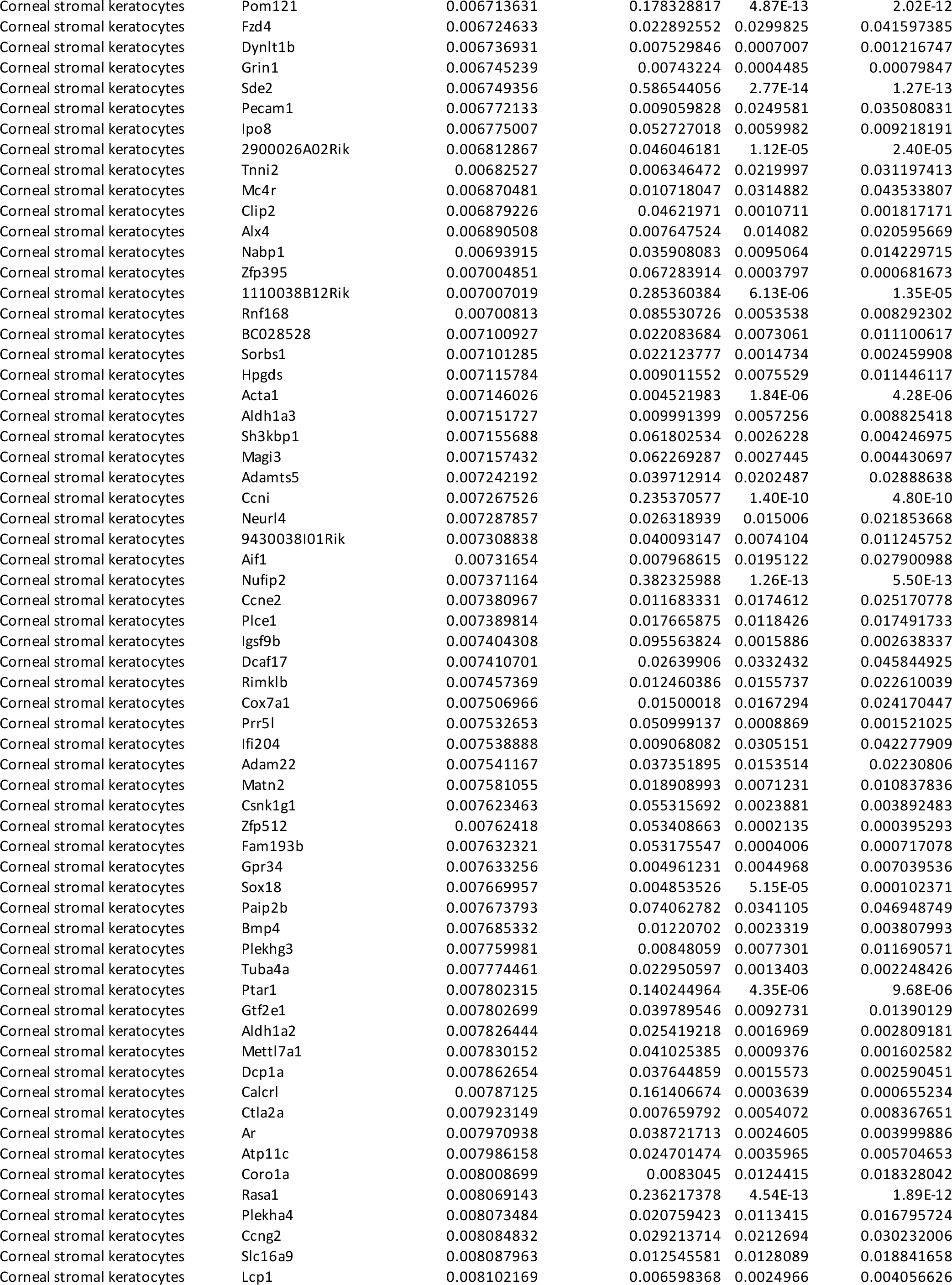
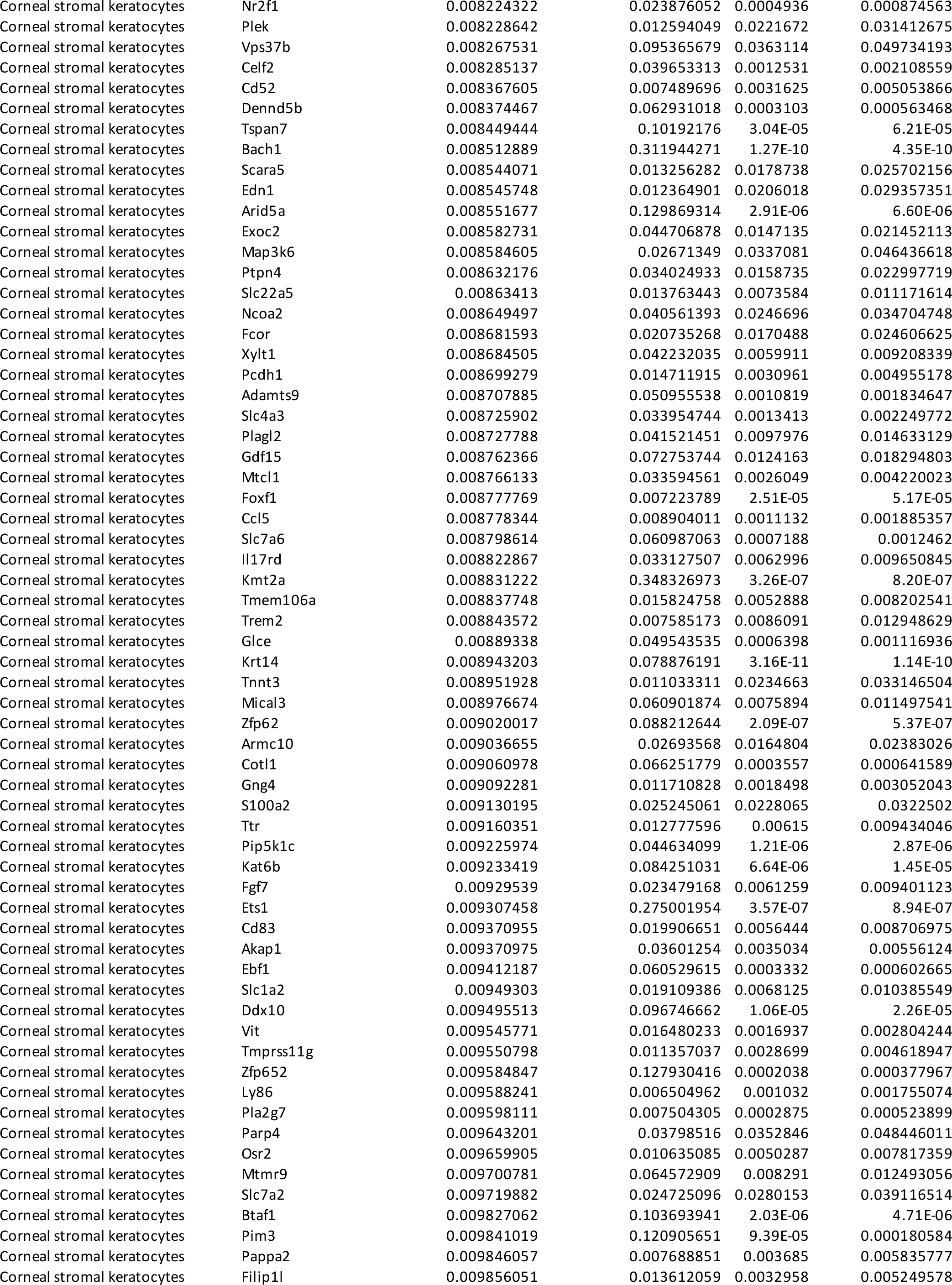
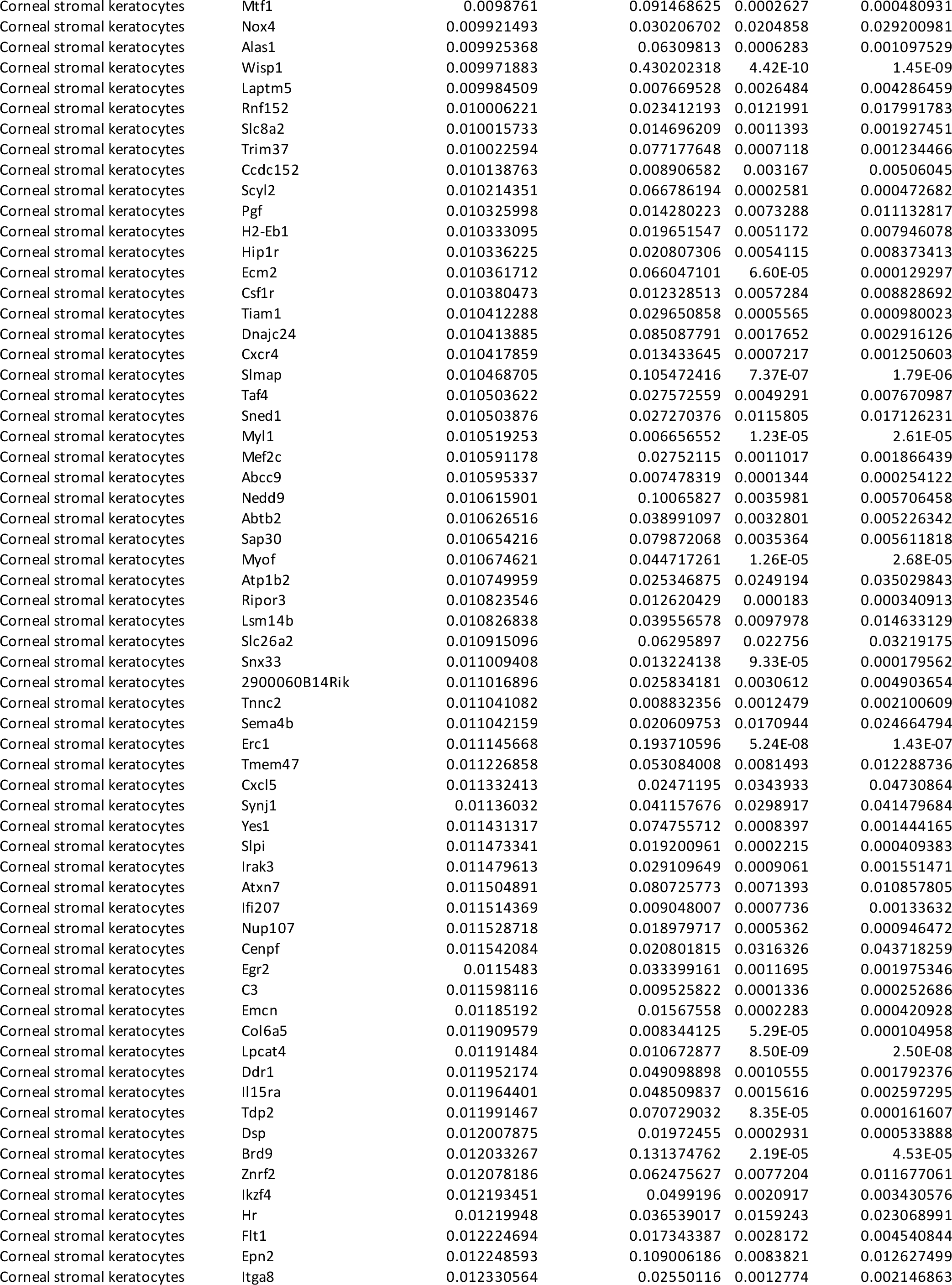
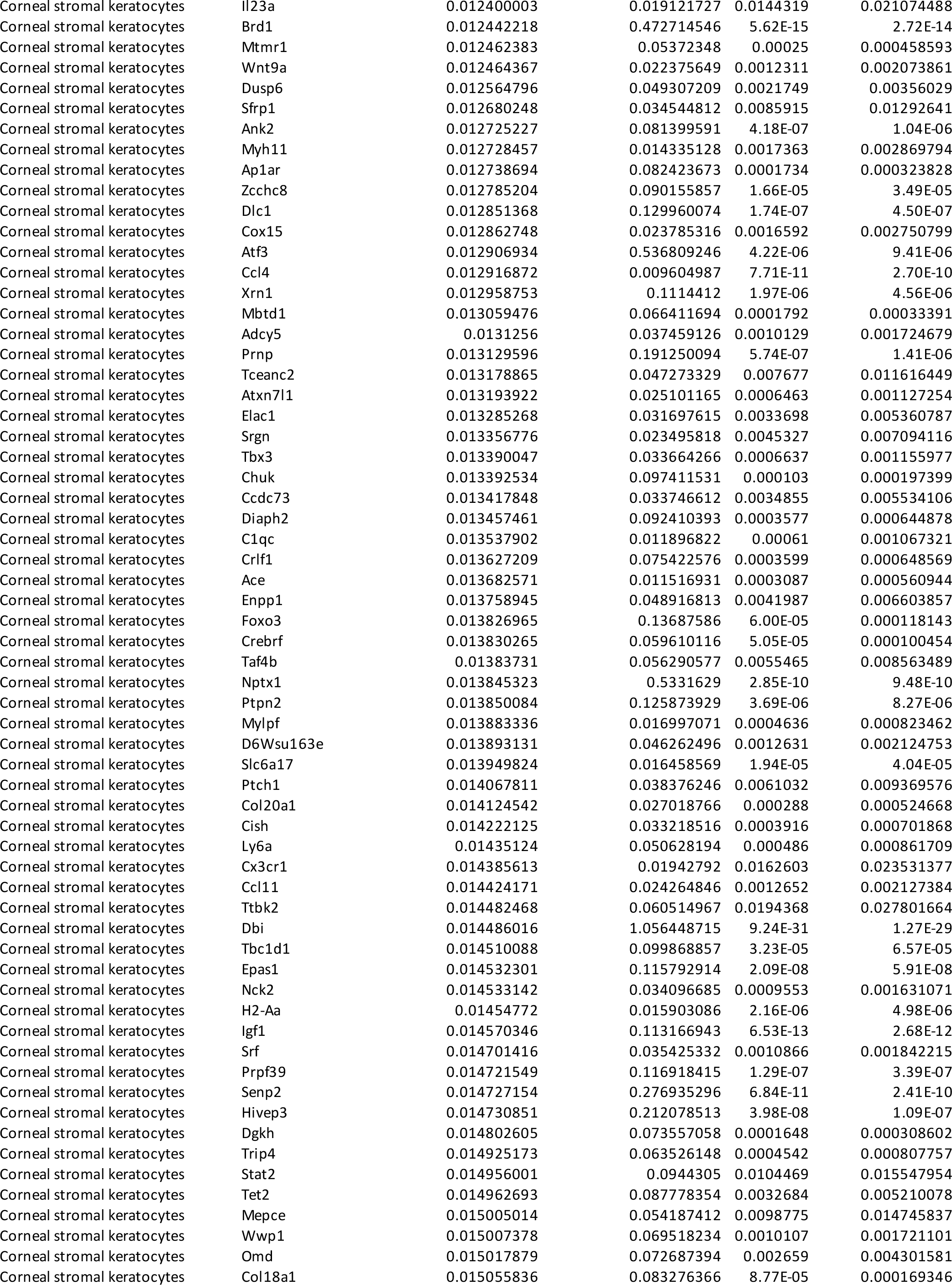
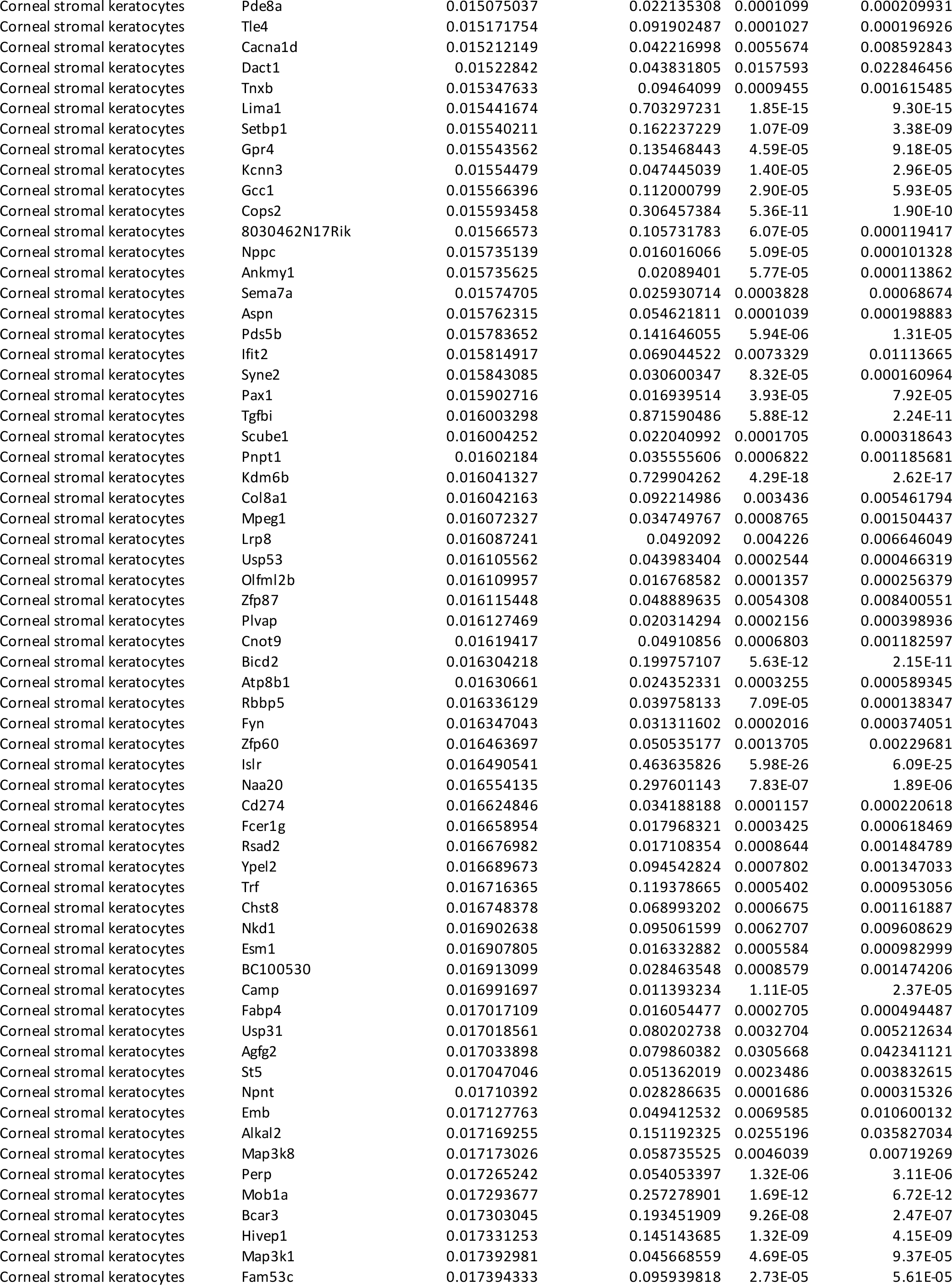
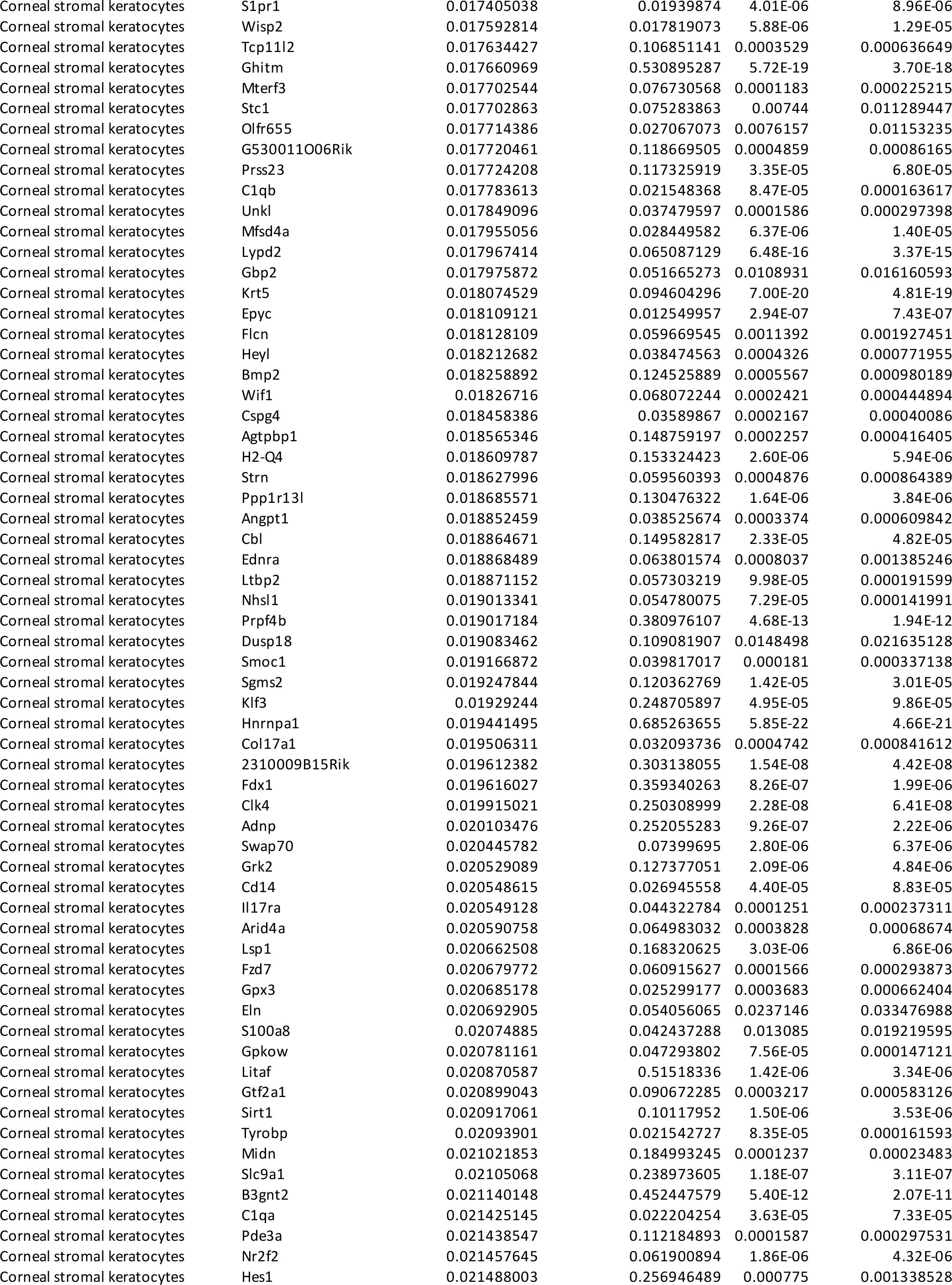
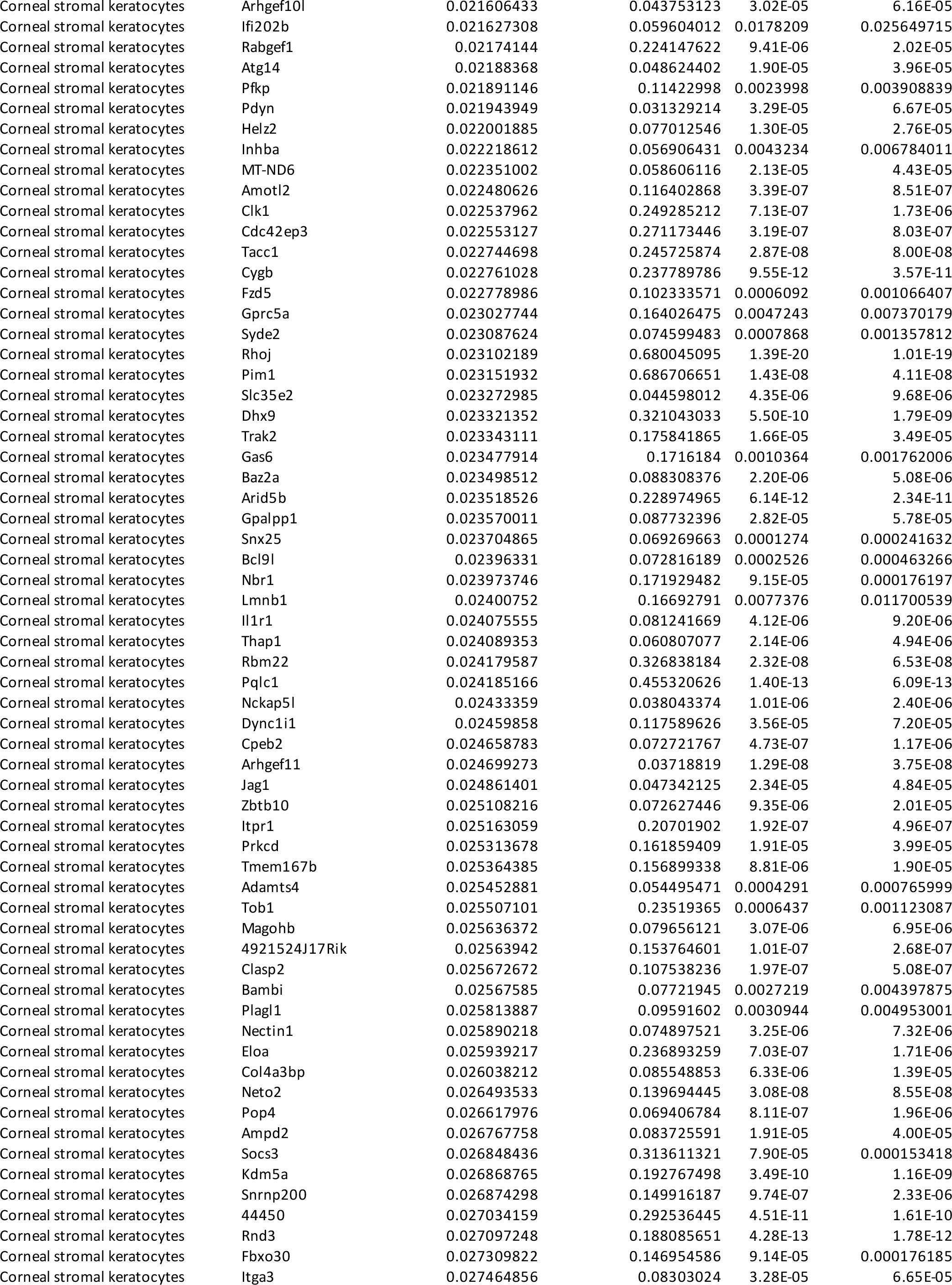
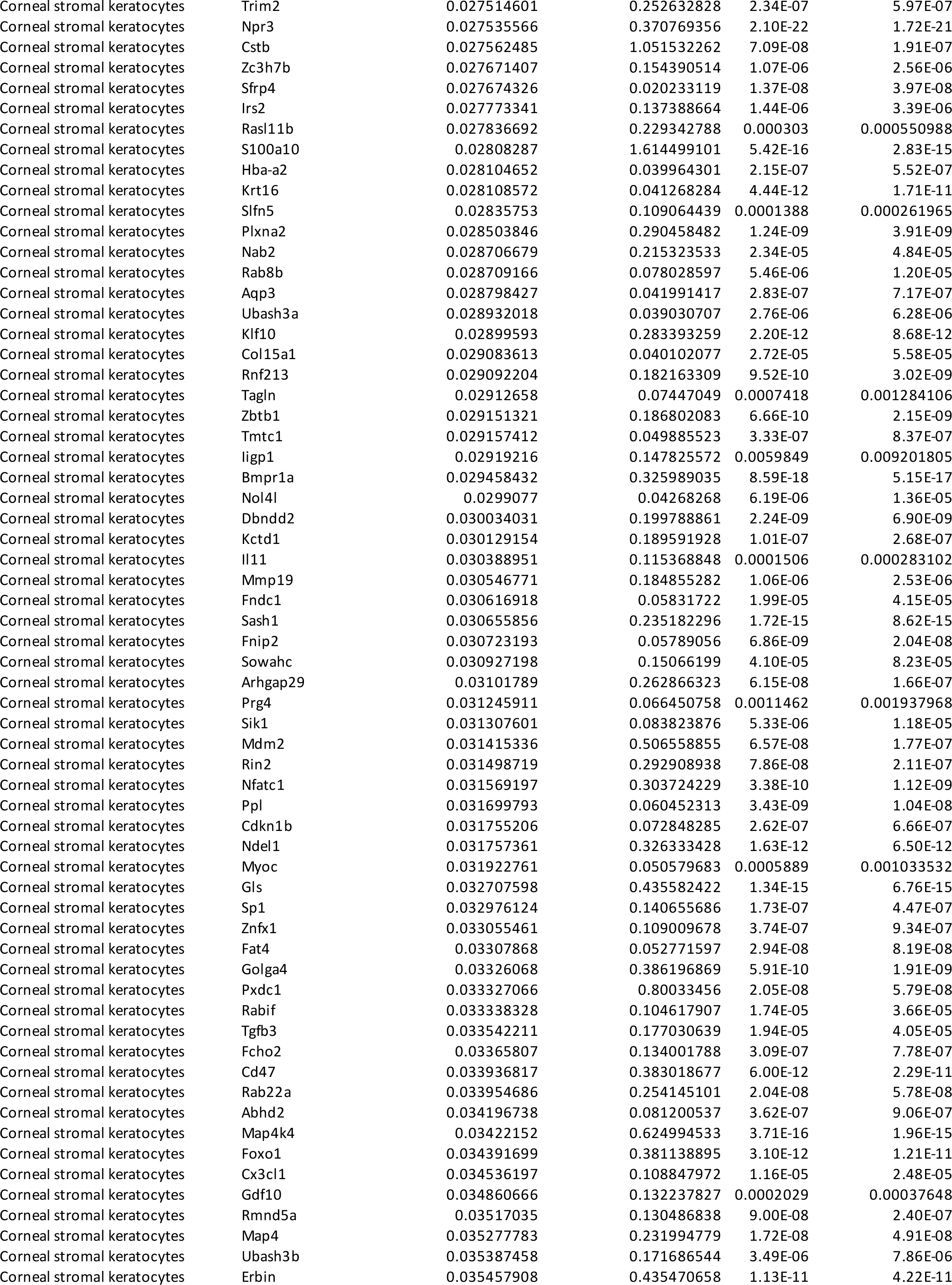
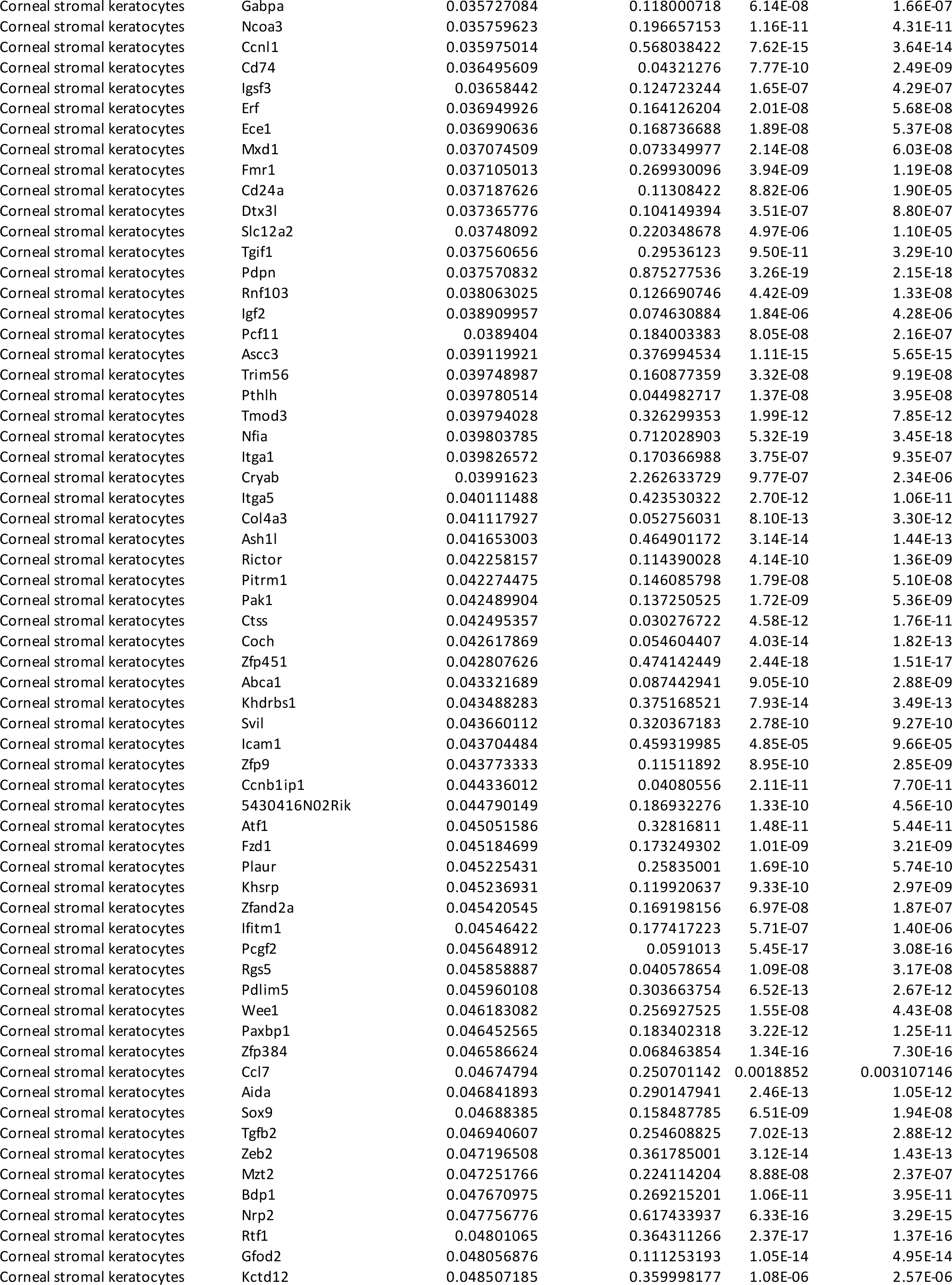
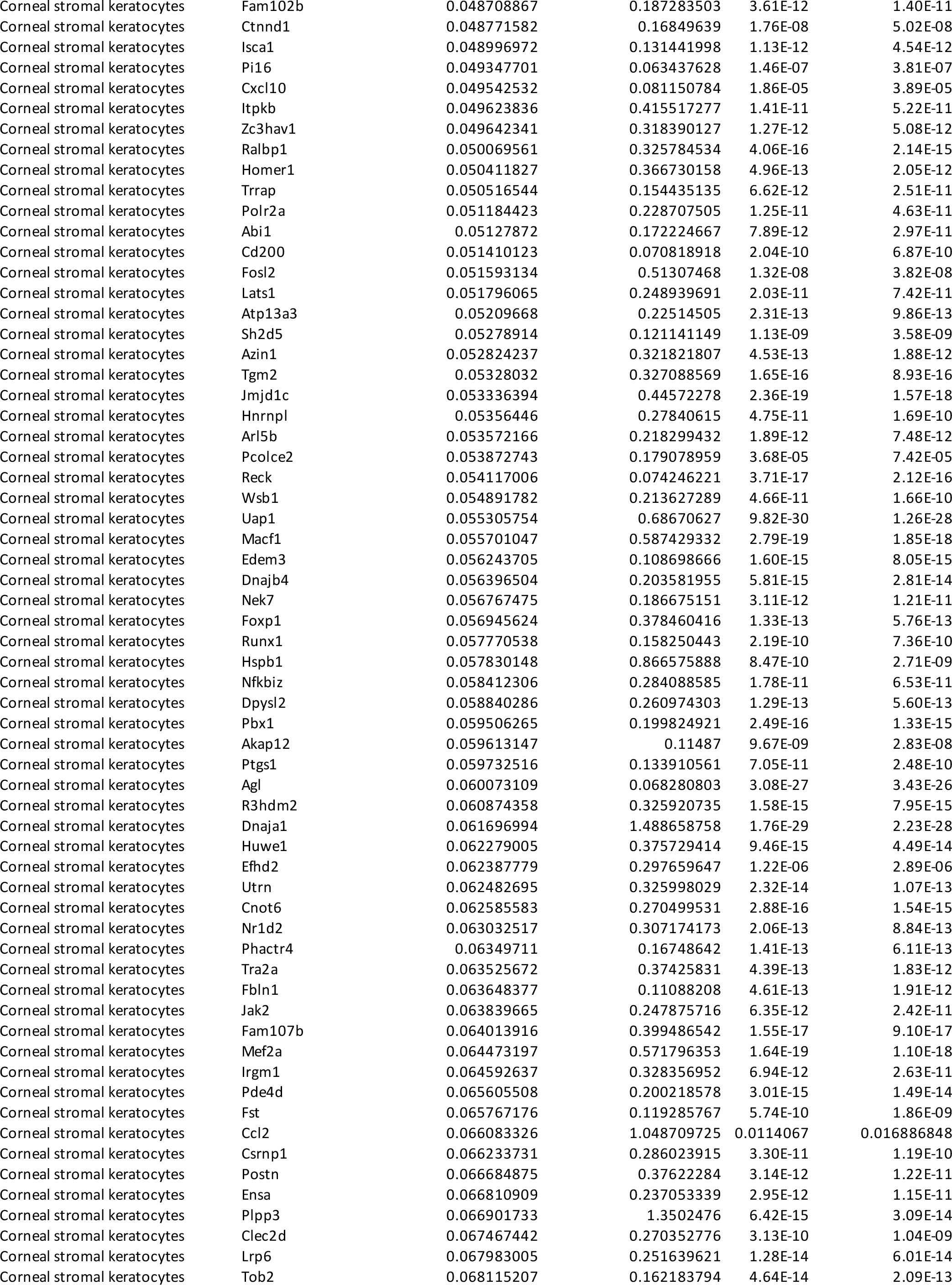
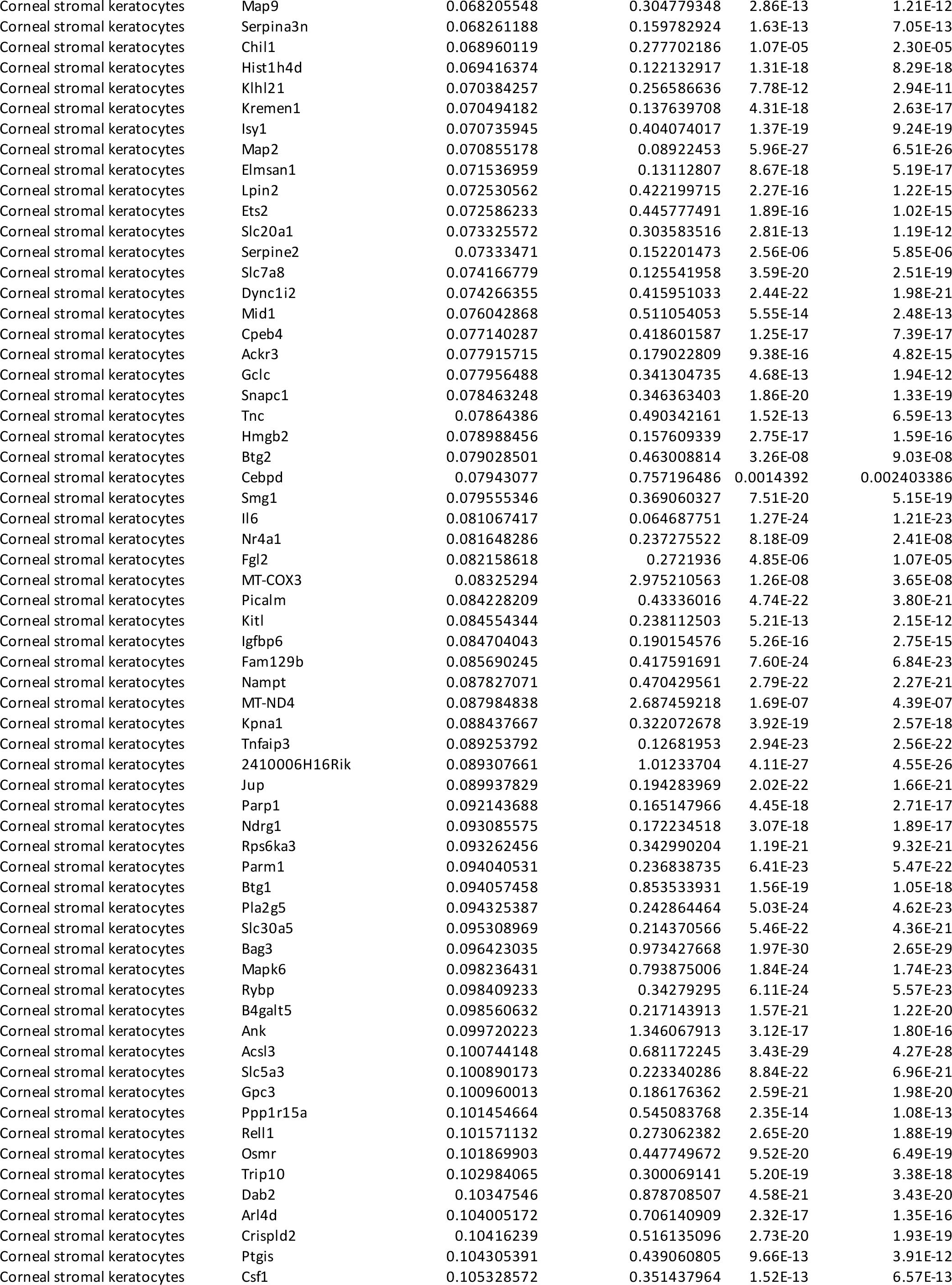
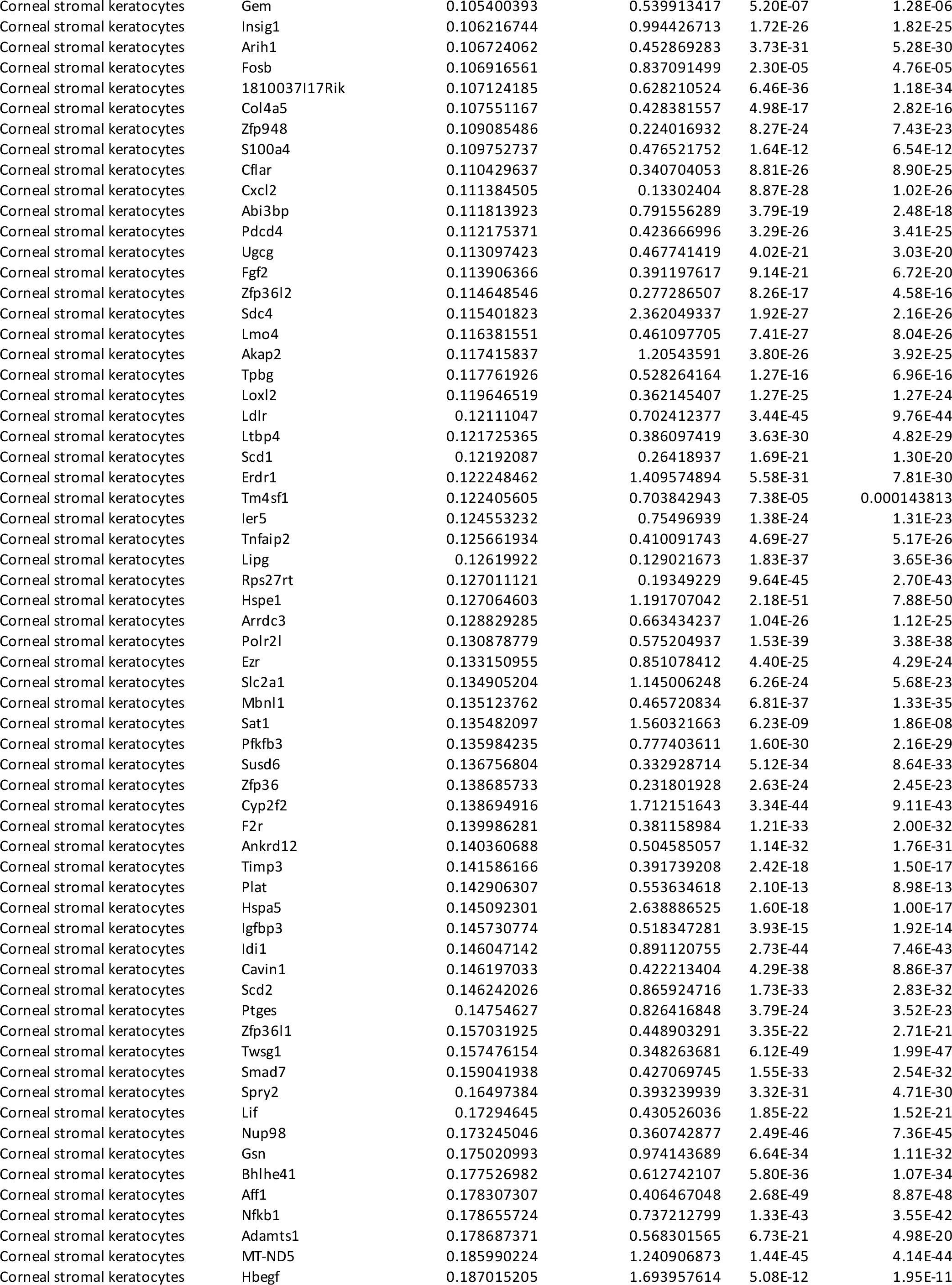
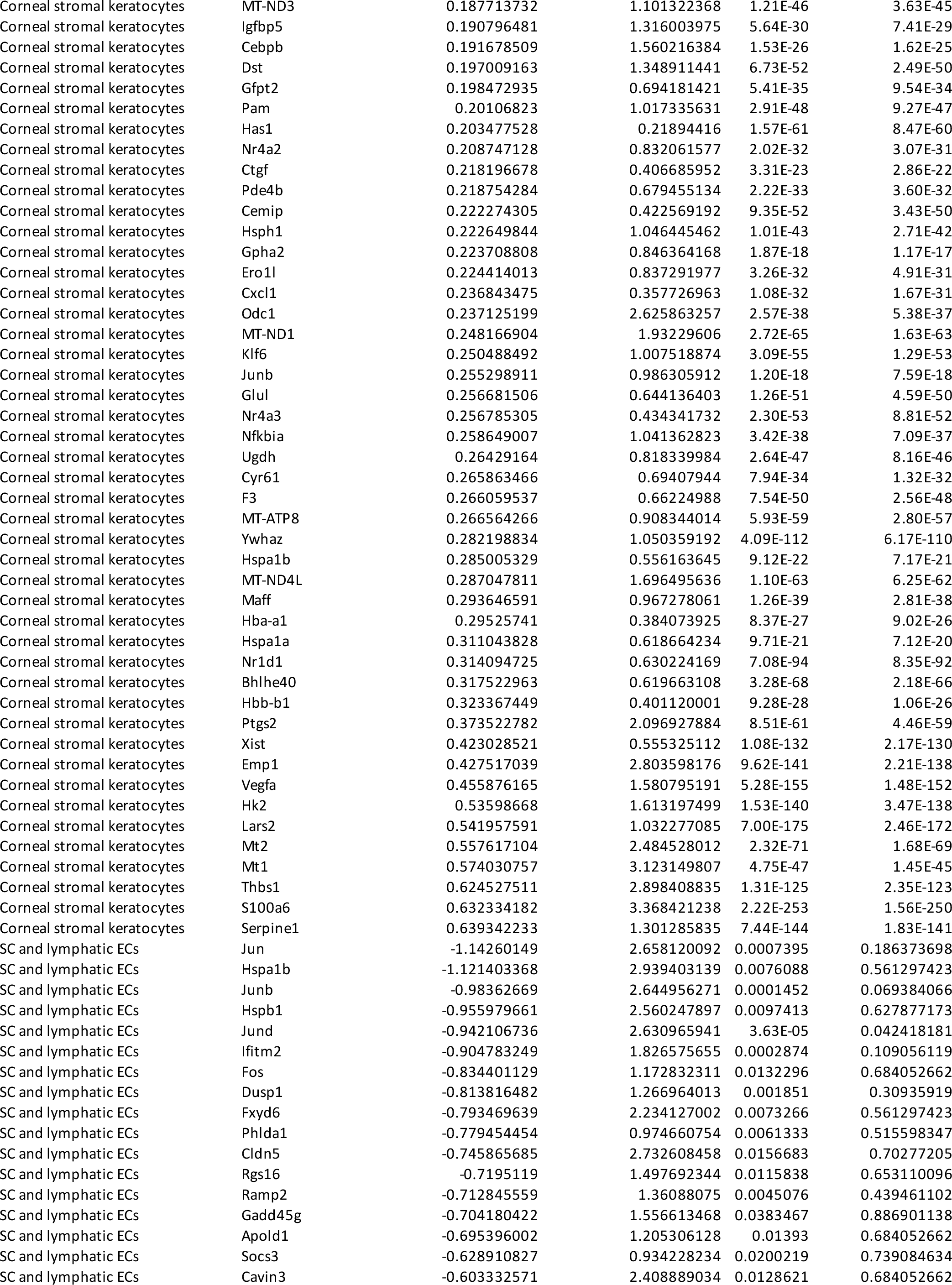
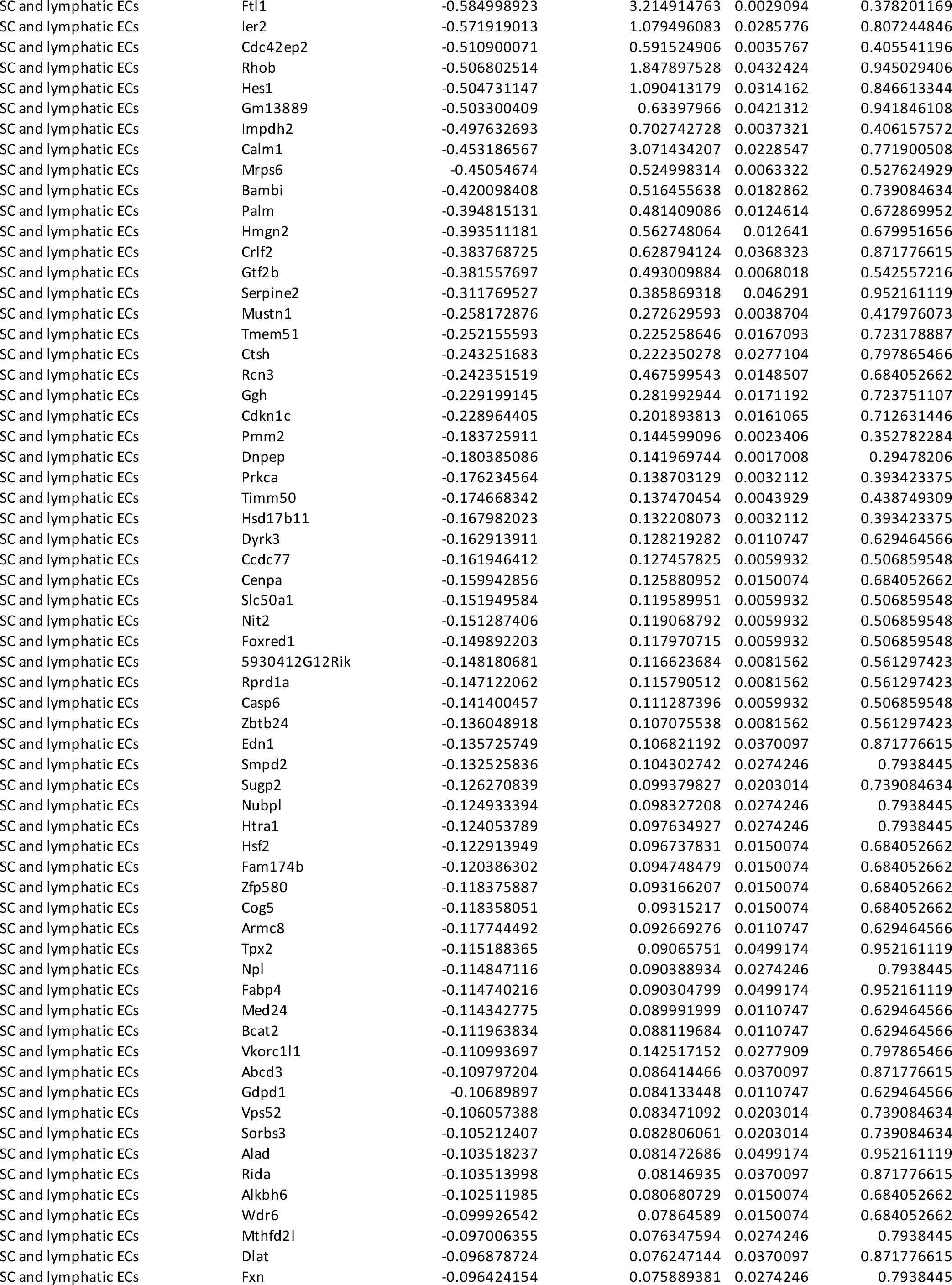
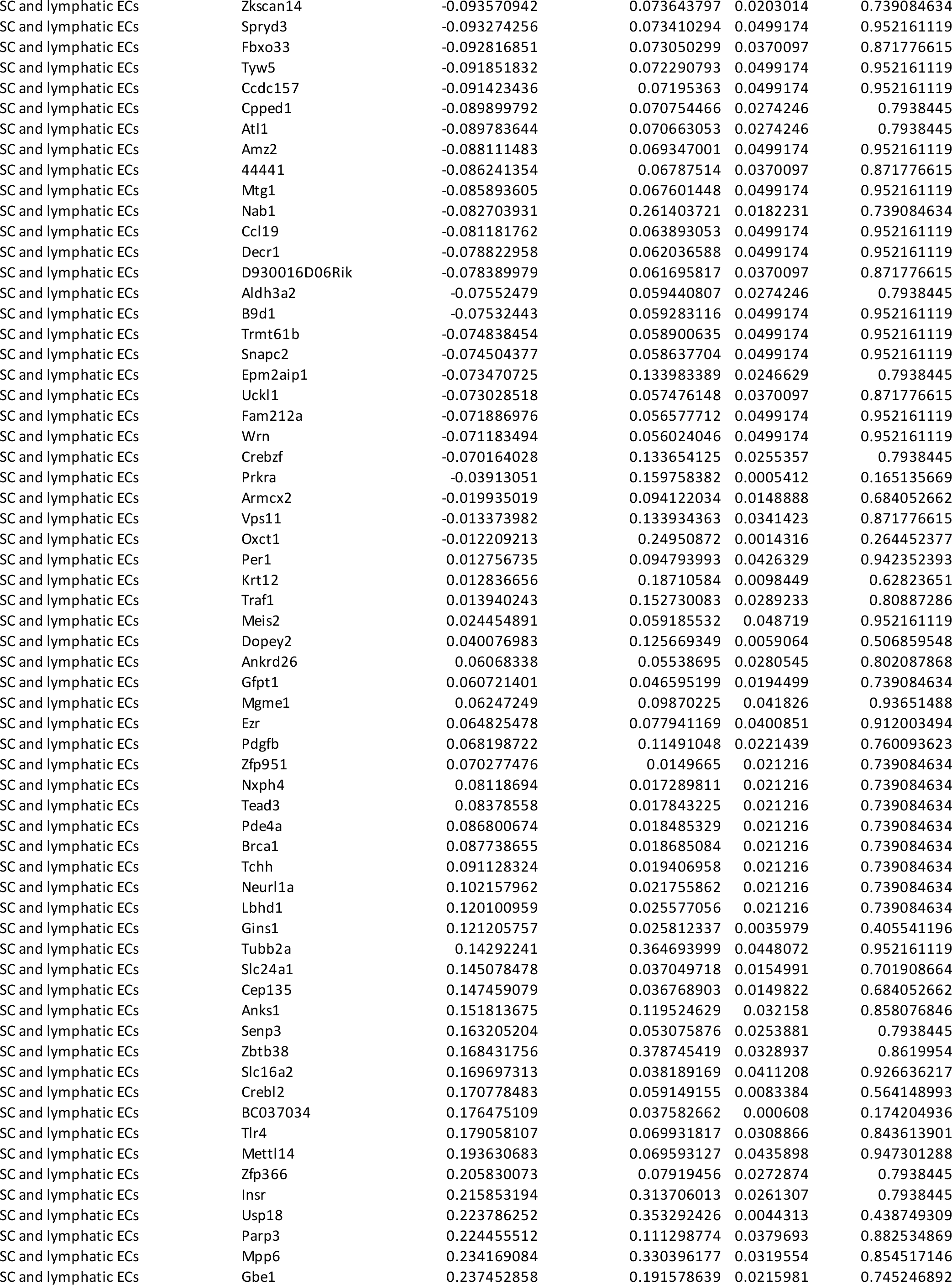
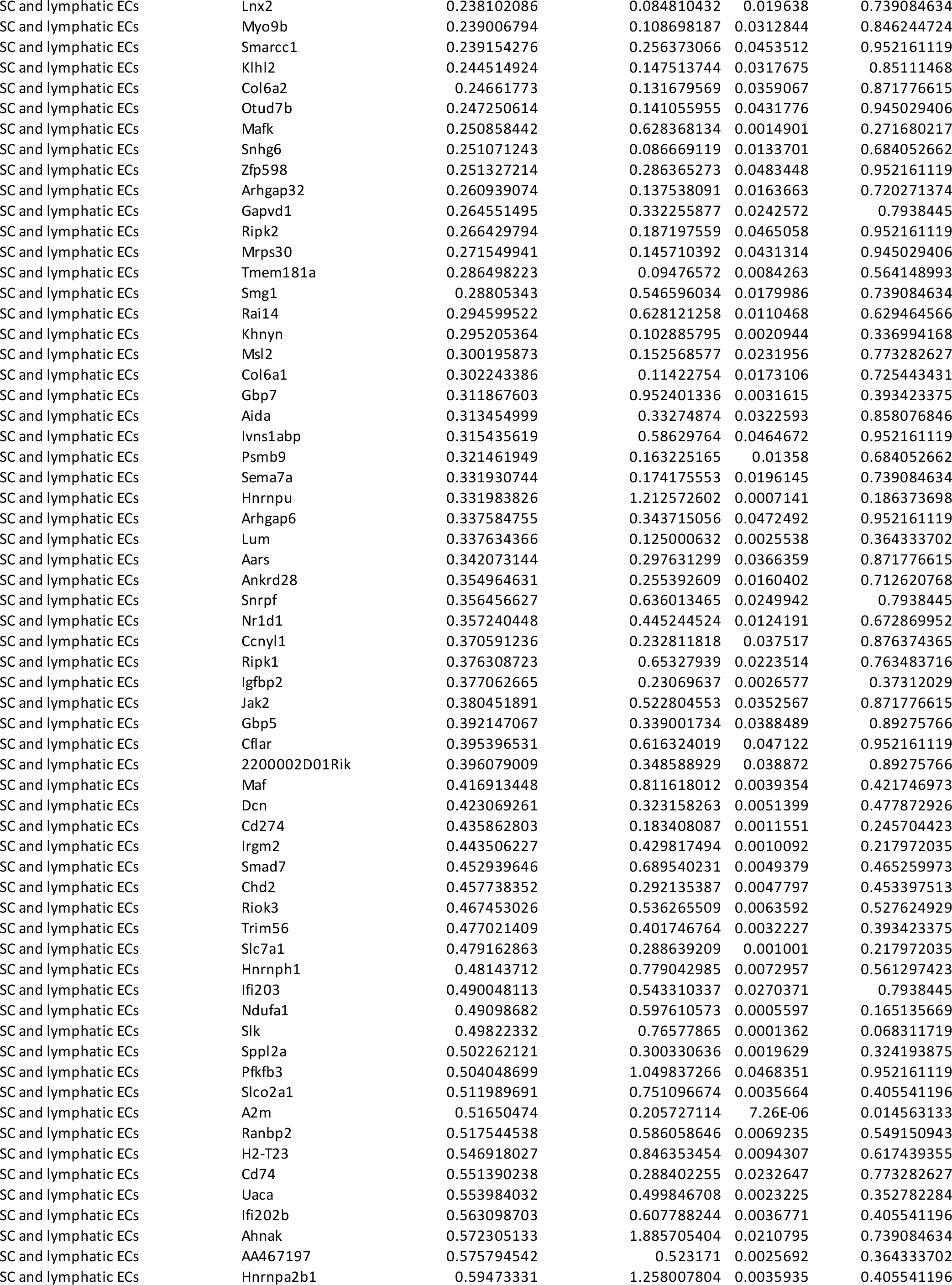
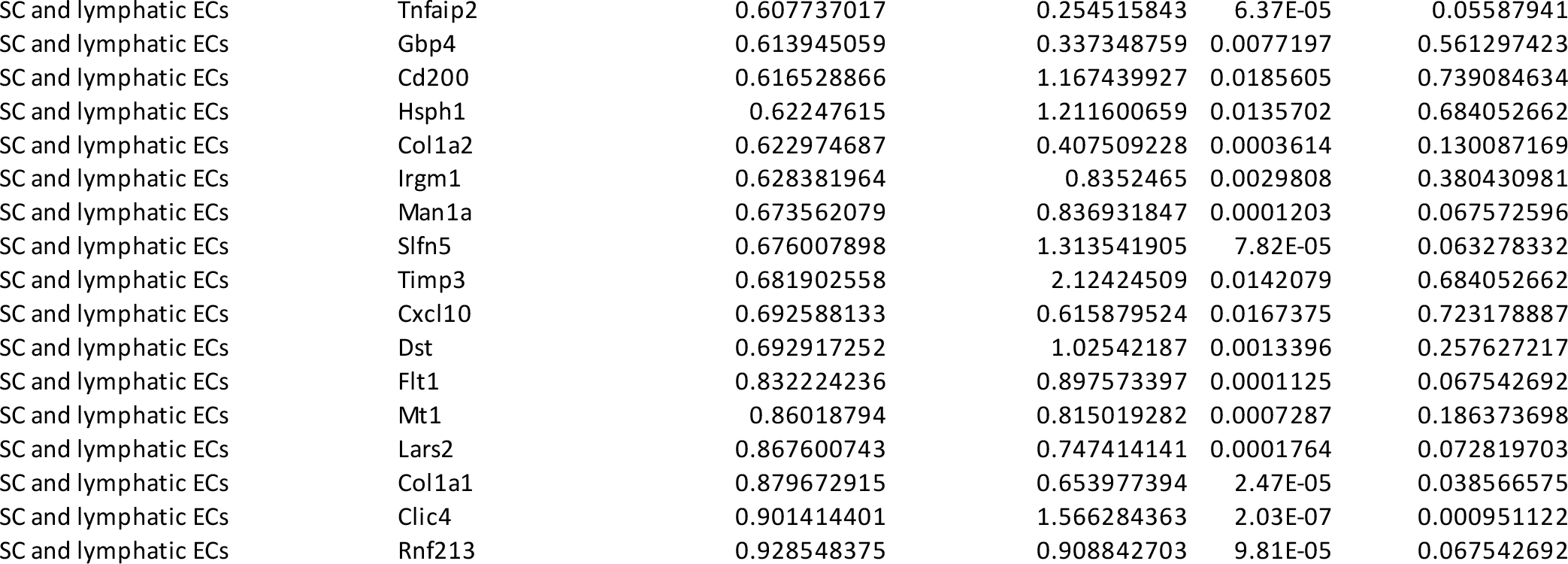
Differentially expressed genes (DEGs) in selected cell clusters from scRNA-seq analysis in *Foxc2^fl/fl^* and NC-*Foxc2^-/-^* mice.

**Supplementary Table 2.** Antibodies and dyes used for immunohistochemistry analysis

| Antigen | Reactivity | Host Species | Origin |
| --- | --- | --- | --- |
| **Primary antibodies** |  |  |  |
| CD31/PECAM1 | Mouse | Rat | BD Pharmingen<br>553370 |
| Endomucin | Mouse | Rat | Abcam ab106100 |
| MMP2 | Human/Mouse/Rat | Rabbit | Proteintech<br>10373-A2-AP |
| PDGFRβ | Mouse | Rat | eBioscience<br>14-1402-81 |
| PROX1 | Human | Goat | R&D AF2727 |
| TIE2 | Mouse/Rat | Goat | R&D AF762 |
| VEGFR-3 | Mouse | Goat | R&D AF743 |
| **Secondary Antibodies** |  |  |  |
| Alexa 488-conjugated | Rabbit/Rat/<br>Goat/Mouse | Donkey | Thermo Fisher |
| Alexa 568-conjugated | Goat/Rat | Goat/Donkey | Thermo Fisher |
| Alexa 647-conjugated | Goat/Mouse/Rabbit<br>Rat | Donkey | Thermo Fisher |
| **Dyes** |  |  |  |
| Hoechst 33342 | - | - | Thermo Fisher |

## References

1. R. Bourne et al., Trends in prevalence of blindness and distance and near vision impairment over 30 years: an analysis for the Global Burden of Disease Study. The Lancet Global Health 9, e130–e143 (2021).

2. J. D. Steinmetz et al., Causes of blindness and vision impairment in 2020 and trends over 30 years, and prevalence of avoidable blindness in relation to VISION 2020: the Right to Sight: an analysis for the Global Burden of Disease Study. The Lancet Global Health 9, e144–e160 (2021).

3. K. Allison, D. Patel, O. Alabi, Epidemiology of Glaucoma: The Past, Present, and Predictions for the Future. Cureus 12, e11686 (2020).

4. C. J. Lewis, et al., Primary congenital and developmental glaucomas. Hum Mol Genet 26, R28–R36 (2017).

5. M. Goel, R. G. Picciani, R. K. Lee, S. K. Bhattacharya, Aqueous humor dynamics: a review. Open Ophthalmol J 4, 52–59 (2010).

6. W. D. Stamer, T. S. Acott, Current understanding of conventional outflow dysfunction in glaucoma. Curr Opin Ophthalmol 23, 135–143 (2012).

7. R. N. Weinreb, T. Aung, F. A. Medeiros, The pathophysiology and treatment of glaucoma: a review. JAMA 311, 1901–1911 (2014).

8. M. H. Strungaru, I. Dinu, M. A. Walter, Genotype-phenotype correlations in Axenfeld-Rieger malformation and glaucoma patients with FOXC1 and PITX2 mutations. Invest Ophthalmol Vis Sci 48, 228–237 (2007).

9. E. I. Traboulsi et al., Lymphedema-distichiasis syndrome and FOXC2 gene mutation. Am J Ophthalmol 134, 592–596 (2002).

10. A. Sabine et al., FOXC2 and fluid shear stress stabilize postnatal lymphatic vasculature. The Journal of Clinical Investigation 125, 3861–3877 (2015).

11. A. González-Loyola et al., FOXC2 controls adult lymphatic endothelial specialization, function, and gut lymphatic barrier preventing multiorgan failure. Science Advances 7, eabf4335 (2021).

12. T. Kume, The cooperative roles of Foxc1 and Foxc2 in cardiovascular development. Adv Exp Med Biol 665, 63–77 (2009).

13. A. Fatima et al., Foxc1 and Foxc2 deletion causes abnormal lymphangiogenesis and correlates with ERK hyperactivation. J Clin Invest 126, 2437–2451 (2016).

14. P. R. Norden et al., Shear stimulation of FOXC1 and FOXC2 differentially regulates cytoskeletal activity during lymphatic valve maturation. Elife 9 (2020).

15. S. Seo et al., Foxc1 and Foxc2 in the Neural Crest Are Required for Ocular Anterior Segment Development. Investigative ophthalmology & visual science 58, 1368–1377 (2017).

16. C. Medina-Trillo et al., Role of FOXC2 and PITX2 rare variants associated with mild functional alterations as modifier factors in congenital glaucoma. PloS one 14, e0211029 (2019).

17. R. S. Smith et al., Haploinsufficiency of the transcription factors FOXC1 and FOXC2 results in aberrant ocular development. Human Molecular Genetics 9, 1021–1032 (2000).

18. A. Aspelund et al., The Schlemm’s canal is a VEGF-C/VEGFR-3-responsive lymphatic-like vessel. J Clin Invest 124, 3975–3986 (2014).

19. K. Kizhatil, M. Ryan, J. K. Marchant, S. Henrich, S. W. John, Schlemm’s canal is a unique vessel with a combination of blood vascular and lymphatic phenotypes that forms by a novel developmental process. PLoS Biol 12, e1001912 (2014).

20. D. Y. Park et al., Lymphatic regulator PROX1 determines Schlemm’s canal integrity and identity. J Clin Invest 124, 3960–3974 (2014).

21. B. R. Thomson et al., A lymphatic defect causes ocular hypertension and glaucoma in mice. J Clin Invest 124, 4320–4324 (2014).

22. Y. Wu et al., Organogenesis and distribution of the ocular lymphatic vessels in the anterior eye. JCI Insight 5 (2020).

23. T. Souma et al., Angiopoietin receptor TEK mutations underlie primary congenital glaucoma with variable expressivity. J Clin Invest 126, 2575–2587 (2016).

24. J. Kim et al., Impaired angiopoietin/Tie2 signaling compromises Schlemm’s canal integrity and induces glaucoma. J Clin Invest 127, 3877–3896 (2017).

25. B. R. Thomson et al., Angiopoietin-1 is required for Schlemm’s canal development in mice and humans. The Journal of clinical investigation 127, 4421–4436 (2017).

26. B. R. Thomson et al., Angiopoietin-1 Knockout Mice as a Genetic Model of Open- Angle Glaucoma. Transl Vis Sci Technol 9, 16 (2020).

27. B. R. Thomson et al., Targeting the vascular-specific phosphatase PTPRB protects against retinal ganglion cell loss in a pre-clinical model of glaucoma. Elife 8 (2019).

28. G. Li et al., A Small Molecule Inhibitor of VE-PTP Activates Tie2 in Schlemm’s Canal Increasing Outflow Facility and Reducing Intraocular Pressure. Investigative Ophthalmology & Visual Science 61, 12–12 (2020).

29. C. M. Findley, M. J. Cudmore, A. Ahmed, C. D. Kontos, VEGF Induces Tie2 Shedding via a Phosphoinositide 3-Kinase/Akt–Dependent Pathway to Modulate Tie2 Signaling. Arteriosclerosis, Thrombosis, and Vascular Biology 27, 2619–2626 (2007).

30. M. B. Amin et al., Foxc2(CreERT2) knock-in mice mark stage-specific Foxc2- expressing cells during mouse organogenesis. Congenit Anom (Kyoto) 57, 24–31 (2017).

31. R. S. Smith, A. Zabaleta, O. V. Savinova, S. W. John, The mouse anterior chamber angle and trabecular meshwork develop without cell death. BMC developmental biology 1, 3 (2001).

32. D. B. Gould, R. S. Smith, S. W. John, Anterior segment development relevant to glaucoma. Int J Dev Biol 48, 1015–1029 (2004).

33. X. Zhang et al., In Vivo Imaging of Schlemm’s Canal and Limbal Vascular Network in Mouse Using Visible-Light OCT. Investigative ophthalmology & visual science 61, 23 (2020).

34. L. Kagemann et al., IOP elevation reduces Schlemm’s canal cross-sectional area. Invest Ophthalmol Vis Sci 55, 1805–1809 (2014).

35. C. Ding, P. Wang, N. Tian, Effect of general anesthetics on IOP in elevated IOP mouse model. Exp Eye Res 92, 512–520 (2011).

36. N. Kaplan et al., Single-Cell RNA Transcriptome Helps Define the Limbal/Corneal Epithelial Stem/Early Transit Amplifying Cells and How Autophagy Affects This Population. Invest Ophthalmol Vis Sci 60, 3570–3583 (2019).

37. G. Patel et al., Molecular taxonomy of human ocular outflow tissues defined by single-cell transcriptomics. Proc Natl Acad Sci U S A 117, 12856–12867 (2020).

38. T. van Zyl et al., Cell atlas of aqueous humor outflow pathways in eyes of humans and four model species provides insight into glaucoma pathogenesis. Proc Natl Acad Sci U S A 117, 10339–10349 (2020).

39. J. Collin, et al., A single cell atlas of human cornea that defines its development, limbal progenitor cells and their interactions with the immune cells. The Ocular Surface https://doi.org/10.1016/j.jtos.2021.03.010 (2021).

40. B. R. Thomson et al., Cellular crosstalk regulates the aqueous humor outflow pathway and provides new targets for glaucoma therapies. Nature Communications 12, 6072 (2021).

41. E. R. Tamm, The trabecular meshwork outflow pathways: structural and functional aspects. Exp Eye Res 88, 648–655 (2009).

42. I. Grierson, W. R. Lee, S. Abraham, R. C. Howes, Associations between the cells of the walls of Schlemm’s canal. Albrecht von Graefes Archiv für klinische und experimentelle Ophthalmologie 208, 33–47 (1978).

43. J. A. Vranka, M. J. Kelley, T. S. Acott, K. E. Keller, Extracellular matrix in the trabecular meshwork: intraocular pressure regulation and dysregulation in glaucoma. Exp Eye Res 133, 112–125 (2015).

44. T. S. Acott, M. J. Kelley, Extracellular matrix in the trabecular meshwork. Exp Eye Res 86, 543–561 (2008).

45. K. E. Keller, J. M. Bradley, T. S. Acott, Differential effects of ADAMTS-1, -4, and - 5 in the trabecular meshwork. Invest Ophthalmol Vis Sci 50, 5769-5777 (2009).

46. T. O. Idowu et al., Identification of specific Tie2 cleavage sites and therapeutic modulation in experimental sepsis. Elife 9 (2020).

47. P. R. Norden, Z. Sun, G. E. Davis, Control of endothelial tubulogenesis by Rab and Ral GTPases, and apical targeting of caveolin-1-labeled vacuoles. PLoS One 15, e0235116 (2020).

48. G. L. Lehmann et al., Single-cell profiling reveals an endothelium-mediated immunomodulatory pathway in the eye choroid. Journal of Experimental Medicine 217, e20190730 (2020).

49. A. S. Shirali et al., A multi-step transcriptional cascade underlies vascular regeneration in vivo. Scientific Reports 8, 5430 (2018).

50. F. Paneni et al., Deletion of the activated protein-1 transcription factor JunD induces oxidative stress and accelerates age-related endothelial dysfunction. Circulation 127, 1229–1240, e1221-1221 (2013).

51. D. R. Overby et al., Altered mechanobiology of Schlemm’s canal endothelial cells in glaucoma. Proceedings of the National Academy of Sciences of the United States of America 111, 13876–13881 (2014).

52. M. Winderlich et al., VE-PTP controls blood vessel development by balancing Tie-2 activity. J Cell Biol 185, 657–671 (2009).

53. T. Souma et al., Context-dependent functions of angiopoietin 2 are determined by the endothelial phosphatase VEPTP. Proc Natl Acad Sci U S A 115, 1298–1303 (2018).

54. S. De Val et al., Combinatorial regulation of endothelial gene expression by ets and forkhead transcription factors. Cell 135, 1053–1064 (2008).

55. Z. Tümer, D. Bach-Holm, Axenfeld-Rieger syndrome and spectrum of PITX2 and FOXC1 mutations. Eur J Hum Genet 17, 1527–1539 (2009).

56. S. Seo et al., Forkhead box transcription factor FoxC1 preserves corneal transparency by regulating vascular growth. Proceedings of the National Academy of Sciences 109, 2015 (2012).

57. J. Sanchez et al., Conditional inactivation of Foxc1 and Foxc2 in neural crest cells leads to cardiac abnormalities. Genesis 58, e23364 (2020).

58. N. Farrar, D. B. Yan, M. Johnson, Modeling the effects of glaucoma surgery on intraocular pressure. Exp Eye Res 209, 108620 (2021).

59. G. Li et al., In vivo measurement of trabecular meshwork stiffness in a corticosteroid-induced ocular hypertensive mouse model. Proc Natl Acad Sci U S A 116, 1714–1722 (2019).

60. H. W. Chung, J. H. Park, C. Yoo, Y. Y. Kim, Effects of Trabecular Meshwork Width and Schlemm’s Canal Area on Intraocular Pressure Reduction in Glaucoma Patients. Korean J Ophthalmol 35, 311–317 (2021).

61. Z. Puyang, H. Chen, X. Liu, Subtype-dependent Morphological and Functional Degeneration of Retinal Ganglion Cells in Mouse Models of Experimental Glaucoma. J Nat Sci 1, e103 (2015).

62. J. A. Schaub et al., Regional Retinal Ganglion Cell Axon Loss in a Murine Glaucoma Model. Invest Ophthalmol Vis Sci 58, 2765–2773 (2017).

63. C. O. Crosby, J. Zoldan, Mimicking the physical cues of the ECM in angiogenic biomaterials. Regenerative Biomaterials 6, 61–73 (2019).

64. L. T. Edgar, C. J. Underwood, J. E. Guilkey, J. B. Hoying, J. A. Weiss, Extracellular Matrix Density Regulates the Rate of Neovessel Growth and Branching in Sprouting Angiogenesis. PLOS ONE 9, e85178 (2014).

65. L. Alderfer, E. Russo, A. Archilla, B. Coe, D. Hanjaya-Putra, Matrix stiffness primes lymphatic tube formation directed by vascular endothelial growth factor-C. FASEB J 35, e21498 (2021).

66. M. Frye et al., Matrix stiffness controls lymphatic vessel formation through regulation of a GATA2-dependent transcriptional program. Nature Communications 9, 1511 (2018).

67. L. De Groef, I. Van Hove, E. Dekeyster, I. Stalmans, L. Moons, MMPs in the Trabecular Meshwork: Promising Targets for Future Glaucoma Therapies? Investigative Ophthalmology & Visual Science 54, 7756–7763 (2013).

68. W. G. Stetler-Stevenson, Matrix metalloproteinases in angiogenesis: a moving target for therapeutic intervention. J Clin Invest 103, 1237–1241 (1999).

69. I. A. Carota et al., Targeting VE-PTP phosphatase protects the kidney from diabetic injury. The Journal of experimental medicine 216, 936–949 (2019).

70. S. Chen et al., Imaging hemodynamic response after ischemic stroke in mouse cortex using visible-light optical coherence tomography. Biomed Opt Express 7, 3377–3389 (2016).

71. F. Taverna et al., BIOMEX: an interactive workflow for (single cell) omics data interpretation and visualization. Nucleic acids research 48, W385–W394 (2020).

72. R. Satija, J. A. Farrell, D. Gennert, A. F. Schier, A. Regev, Spatial reconstruction of single-cell gene expression data. Nat Biotechnol 33, 495–502 (2015).

73. J. Kalucka et al., Single-Cell Transcriptome Atlas of Murine Endothelial Cells. Cell 180, 764–779 e720 (2020).

74. J. W. Shin et al., Prox1 promotes lineage-specific expression of fibroblast growth factor (FGF) receptor-3 in lymphatic endothelium: a role for FGF signaling in lymphangiogenesis. Mol Biol Cell 17, 576–584 (2006).

75. Y. Zhou et al., Metascape provides a biologist-oriented resource for the analysis of systems-level datasets. Nat Commun 10, 1523 (2019).

